# Ontology-guided harmonization enables unified discovery of public metabolomics studies within and across repositories

**DOI:** 10.64898/2026.07.23.740366

**Authors:** Shayantan Banerjee, Pratik Jalan, Rudrapratap Chinhara, Chinmayee Kalle, Pramod P Wangikar, Kshitij Jadhav

## Abstract

Public metabolomics repositories contain thousands of studies, but differences in metadata structure, vocabulary, and repository-specific terms still limit reliable search, comparison, and reuse within and across databases. Here we present HARMONY, an ontology-based framework and web platform that harmonizes study-level metadata and metabolite information across Metabolomics Workbench and MetaboLights studies. HARMONY resolves eight biological and analytical metadata nodes, including species, sample source, disease, analytical technique, separation method, ion polarity, ionization source, and mass analyzer type, while preserving the original deposited terms as evidence. A ninth node, metabolite identity, maps metabolite entities to RefMet across both repositories. HARMONY uses a two-step workflow: Multi-source extraction retrieves records missed by single-field lookups, and ontology mapping then converts repository-specific labels into shared query terms, substantially closing the cross-repository retrieval gap relative to raw matching. Across the full corpus, HARMONY increased cross-repository retrievability from 75.5% to 89.6%, yielding thousands of study-node retrievals and reconnecting studies that raw-text search would have left unreachable within their own repositories. Approximately 91% of Metabolomics Workbench and 85% of MetaboLights studies had at least six of the eight nodes harmonized. The resulting platform, available at https://omicsinharmony.in, supports ontology-aware search, metadata filtering, within- and cross-repository study comparison, and metabolite-level querying, with retrieval backed by machine learning encoders that map study metadata into shared representations of biological and analytical context. HARMONY provides the metabolomics community with a shared, traceable search interface for study discovery and comparison within and across public repositories.

## Introduction

Metabolomics is the systematic measurement of small-molecule metabolites in biological systems. Over the past decade, LC-MS-based metabolomics has evolved from a niche specialty into a major omics platform for studying metabolism and broader biomolecular phenotypes, generating thousands of publicly archived datasets^1,2^. This growth has been supported by major public funding. Programs such as the NIH Common Fund’s decade-long Metabolomics Program^3^ have funded core facilities, repositories, and standards development, and the experiments themselves. Such studies can cost tens of thousands of dollars once instrument time, sample preparation, and analyst effort are considered. Yet only a small share of this work is deposited in public repositories. A recent comparison^4^ of published MS-based metabolomics articles with publicly shared datasets over the same five-year period found that fewer than 20% of articles were accompanied by a newly deposited public dataset, indicating that most published studies did not make their underlying data publicly available. Even among the deposited studies, differences in instrumentation, experimental workflows, reporting practices, and metadata representation fragment the public data landscape into incompatible “islands” that are difficult to compare or reuse^4–7^. This lack of coherence has hindered the development of large, interoperable metabolomics resources and has limited both cross-study discovery and integration with other omics disciplines.

Much of this public landscape is now concentrated in two leading repositories: Metabolomics Workbench (MW)^8^, part of the NIH Common Fund’s Metabolomics Program, and MetaboLights (ML)^9^, the EMBL-EBI repository for metabolomics studies and associated metadata. Together, they archive around 6,830 independent studies that are publicly accessible (as on 9th March 2026), covering human disease, model organisms, plant biochemistry, environmental science, and food chemistry. If these studies could be queried through shared biological and analytical concepts, they would provide a foundation for cross-repository discovery^4,10^, including assembling comparable, analysis-ready cohorts across databases^11^, comparing metabolic phenotypes across species and sample sources, identifying recurring metabolite–phenotype patterns, evaluating how analytical platforms shape reported findings, and exposing technical or biological confounders that remain invisible within any single repository.

This possibility, however, is limited by a persistent interoperability problem that is already visible in the deposited data^4,7^. The Metabolomics Standards Initiative (MSI), established in 2005, published reporting guidelines in 2007 to define minimum descriptions for metabolomics experiments, including experimental design, biological context, chemical analysis, and data processing^12^, an early effort toward interoperable reporting. The FAIR principles, later formalized, set the expectation that scientific data should be Findable, Accessible, Interoperable, and Reusable, and helped shape the data-sharing practices adopted by repositories such as MW and ML. Of these four principles, accessibility is largely determined by repository policy and infrastructure, which fall outside the scope of metadata harmonization. The reusability of deposited tabular data for machine learning applications has been addressed separately through source-aware readiness assessment of repository-hosted matrices in MW^11^. The reusability of raw spectral data has been addressed through frameworks such as Pan-ReDU^4^ which harmonize sample metadata and index mass spectrometry files across repositories, making spectra easier to search and reuse for reanalysis. Related efforts, including denoising tools^13^, further improve the quality of raw spectra prior to annotation. The remaining two principles, findability and interoperability, stay limited by the semantic inconsistencies described below.

Findability remains constrained at the level of biological description, as shown by an audit^6^ of 399 public studies from four metabolomics repositories, in which none of the MSI biological-context standards were satisfied by all studies, and compliance with individual standards ranged from 0 to 97%. Interoperability is similarly constrained at the analytical level. A review of chromatographic metadata in public repositories reported that 70% of chromatographic setup descriptions were incomplete and another 10% were ambiguous or incorrect, leaving only about one in five records suitable for downstream uses such as retention-time prediction^14^. Tools such as MW File Status^15^ help address findability by checking file-format validity, but they do not resolve the underlying mismatch in meaning across curator strings, partial ontology identifiers, and repository-specific terms. It checks file formats, but not the semantic mismatches in metadata that still prevent interoperability across repositories. Part of the problem is that much of the needed instrument information is already in the raw files, but researchers still have to enter it manually, and limited time, unfamiliar interfaces, and resource constraints make that difficult^16^. Current efforts, such as the one presented in this study can improve search and comparability only for what is actually deposited, but accessibility still depends on what is made public in the first place. Within individual repositories, controlled vocabularies and ontologies such as Chemical Methods Ontology (CHMO)^17^, the HUPO-PSI mass spectrometry ontology (PSI-MS)^18^, and the NCBI Taxonomy^19^ provide the basis for structured annotation, but legacy records and uneven labeling still leave the same concept scattered across different free-text or repository-specific terms. What is still needed is a harmonization layer that maps study-level biological and analytical descriptors to shared ontology-linked concepts across MW and ML.

Here we present HARMONY (Harmonized Annotation and Resolution of Metabolomics Ontologies across Node-mapped repository data), a computational pipeline for harmonizing study-level metadata from Metabolomics Workbench and MetaboLights. By resolving repository-specific metadata into shared, ontology-linked concepts, HARMONY directly targets the “Findability” and “Interoperability” components of the FAIR framework. HARMONY is freely available online (https://omicsinharmony.in) and maps raw repository annotations to shared ontology-linked concepts across biological and analytical nodes, while retaining the original deposited terms as evidence. Across 4,118 MW and 2,712 MetaboLights studies (available as of 9th March 2026), it resolves metadata spanning species, sample source, disease, analytical technique, separation method, ion polarity, ionization source, mass analyzer type, chromatography instrument, RefMet-linked metabolite identity, and searchable differentially abundant metabolites (DEMs) across studies. We use this resource to report node-level mapping coverage, compare raw-string and ontology-based cross-repository study matching, and support ontology-aware study discovery on the HARMONY platform using machine learning encoders that retrieve studies when exact-term matches are unavailable.

## Results

### Species mapping shows broad coverage and distinct taxonomic structure across repositories

Species annotations were mapped to NCBI Taxonomy with high coverage in both repositories (Figure 1A). Unmapped records (MW:1.45%; ML: 4.64%) were manually reviewed and were retained as null species IRIs when they represented missing organism metadata, blanks, or QC samples, environmental or community descriptors, food or material mixtures, analytical controls, or cases requiring biological inference beyond the deposited term (Table S6). The mapped taxonomic space differed substantially between repositories despite comparable coverage rates. MW was concentrated around a small set of high-frequency taxa, including human, mouse, rat, and a handful of standard model organisms, with the top taxa accounting for a disproportionately large fraction of all study-level mappings, all resolved via exact OLS label matches. ML, despite having fewer studies, spanned a broader NCBITaxon vocabulary than MW and was harder to resolve. MW mappings relied entirely on exact label matches, whereas ML required exact, synonym-level, token-compatible, and manually reviewed matches to achieve similar coverage. Many taxa in ML appeared in only a single study, producing a pronounced singleton long tail (Figure 1B; Table S6). This breadth extended in taxonomic rank as well. ML contained more multi-species studies and a higher proportion of annotations resolved at genus, subspecies, or strain level rather than species rank, which is why rank-preserving mapping was necessary rather than collapsing all organism entries to species (Figure 1B, Figure S1A-B, Table S6). Overall, MW therefore represents a taxonomically concentrated resource built around a core of well-studied organisms. At the same time, ML captures a wider and more heterogeneous biological space, including plants, livestock, fungi, microbes, and non-model organisms that are underrepresented in MW.

**Figure 1:**
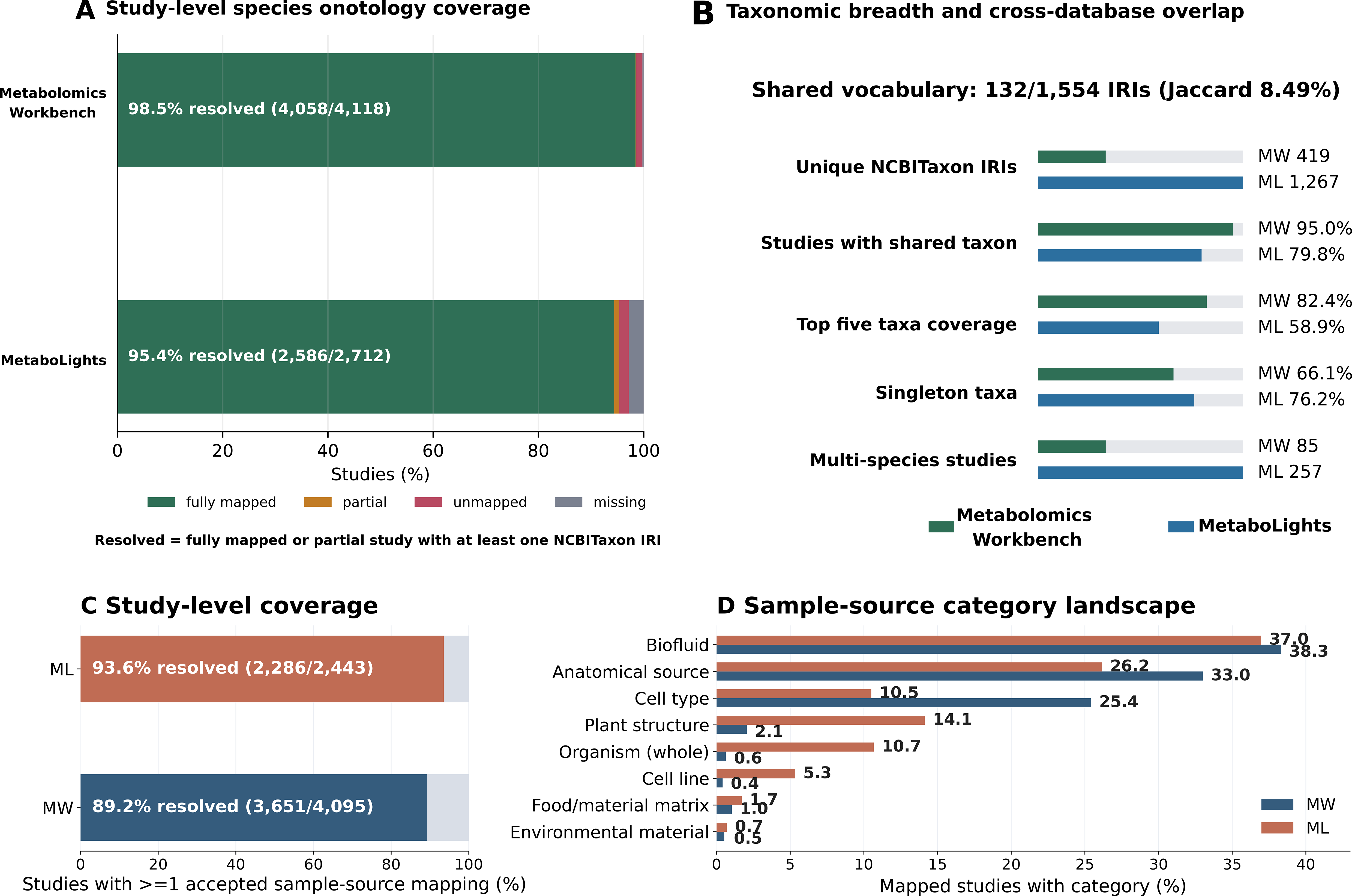
Species and sample source ontology mapping coverage, resolution evidence, and cross-database taxonomic structure. A) Study-level species mapping status; a study counts as resolved if at least one organism entry carries a valid NCBITaxon IRI. B) Taxonomic breadth and cross-repository overlap after mapping, bar length shows each metric’s value, with the grey extension indicating headroom to 100%. C) Study-level sample-source mapping status, restricted to studies with at least one non-empty raw source term. D) Distribution of mapped sample-source categories; studies contributing more than one category are counted in each.

At the vocabulary level, only 132 NCBITaxon IRIs were shared between MW and ML, giving a Jaccard overlap of 8.49%. Although low, it reflects the genuine difference in biological scope between the two repositories rather than a failure of harmonization. More importantly, these 132 shared taxa were sufficient to connect 95.0% of mapped MW studies and 79.8% of mapped ML studies to at least one counterpart in the other repository. Ontology mapping, therefore, converted a narrow shared vocabulary into a cross-repository species layer broad enough to support meaningful study retrieval across both databases.

### Sample-source mapping captures diverse specimen types across both repositories

Sample-source mapping achieved high study-level coverage across both repositories. Deposited terms spanned tissues, biofluids, cell types, cell lines, whole organisms, plant structures, environmental materials, food matrices, culture-derived materials, and experimental descriptors. This led to a broader semantic range than the species node, ultimately requiring a multi-ontology mapping strategy rather than a single anatomy-centered vocabulary (Figure 1C). Across both repositories, 955 unique accepted raw source terms resolved to 673 unique ontology IRIs (Table S7). Within each repository, a compact set of high-frequency IRIs accounted for most mapped studies: 24 IRIs covered 90.6% of MW mapped studies, and 55 IRIs covered 90.3% of ML mapped studies; the remaining ∼10% of studies were distributed across 161 (MW) and 536 (ML) lower-frequency IRIs. Biofluids dominated the mapped source landscape in both repositories, with plasma, serum, and urine the most frequently mapped specimen classes, consistent with their prevalence in clinical metabolomics^1,20^ (Figure 1D). ML showed greater representation of plant structures, whole-organism sources, cell lines, and heterogeneous material descriptors than MW, which drove the need for ontology coverage across UBERON and BTO for anatomical and biofluid terms, CL and CLO for cell-related terms, PO for plant structures, ENVO for environmental materials, and FOODON for food matrices, with reviewed secondary and fallback ontologies applied where the resolved concept represented a valid source material (Figures S2A, S2F). Most unique mapped terms were resolved by exact label matching. Smaller subsets required constrained token-compatible matching or manual curation (Figure S2E). At the row level, exact accepted mappings dominated, with constrained, caveated, rejected, unmapped, and missing rows retained as distinct decision classes to allow downstream queries to tier results by evidence quality rather than treating all non-exact mappings equally (Figure S2B). Residual null or rejected entries covered ambiguous deposits, specimen-processing descriptors, blank and QC terms, analyte or chemical labels, disease-context terms, and assay or procedure metadata (Figure S2C; Table S7), with representative curation decisions documented in Figure S2D. Cross-repository vocabulary overlap was modest (Jaccard 15.3%; 103/673 shared IRIs), but ontology-harmonized source IRIs recovered 13,880 MW–ML study pairs missed by exact raw-string matching alone.

### Conservative disease harmonization recovers shared disease concepts while preventing over-assignment in non-disease studies

Disease-node recovery reflected a fundamental structural difference between repositories (Table S27). MW exposes a dedicated REST disease endpoint, from which 2,466 of 4,118 studies carried at least one disease term; 2,458 of these received a strictly accepted disease IRI. ML required evidence-based extraction from study descriptions, design descriptors, factor names, and "Factor Value[]" fields, yielding 1,369 disease-extractable studies of 2,712, of which 1,184 carried a strictly accepted IRI, and 1,306 carried a strict or broad accepted IRI. Extractable ML studies without a safe ontology match were retained as valid disease literals rather than assigned misleading IRIs. The ML non-extractable group was explicitly curated rather than treated as unannotated (Table S28; Figure S10). It comprised method or resource workflows; plant- and food-based studies without disease endpoints; diet and exposure studies; microbial ecology and biofilm studies without host disease; normal developmental biology; environmental stress; in vitro perturbation; healthy reference metabolomes; and exercise physiology. This boundary prevented background terms, such as infection, resistance, inflammation, pathogen, tumor, biofilm, from inflating disease coverage where the study design did not support a disease assignment.

The mapped disease landscape differed between repositories (Figure 2A–B). MW was dominated by broad REST endpoint labels: cancer, diabetes mellitus, obesity, environmental exposure, malaria, bacterial infection, inflammation, and organ-system categories. ML showed a more granular structure after manual boundary review (Table S29). For visualization, ML concepts were grouped into broad disease families, while the underlying record retained full audited labels, IRIs, ontology namespaces, synonyms, parents, and hierarchy summaries. Cross-repository overlap was modest but meaningful. After cross-ontology equivalence cleanup, MW contained 233 strict disease concepts and ML 605, with 123 shared across a union of 715 (Jaccard 17.2%). The gap reflects MW’s broad endpoint labels against ML’s more specific subtypes and clinical conditions. However, the shared concepts covered major disease areas: diabetes, obesity, COVID-19, malaria, metabolic syndrome, fatty liver disease, sepsis, neurodegeneration, inflammatory bowel disease, cardiovascular disease, and kidney disease.

**Figure 2:**
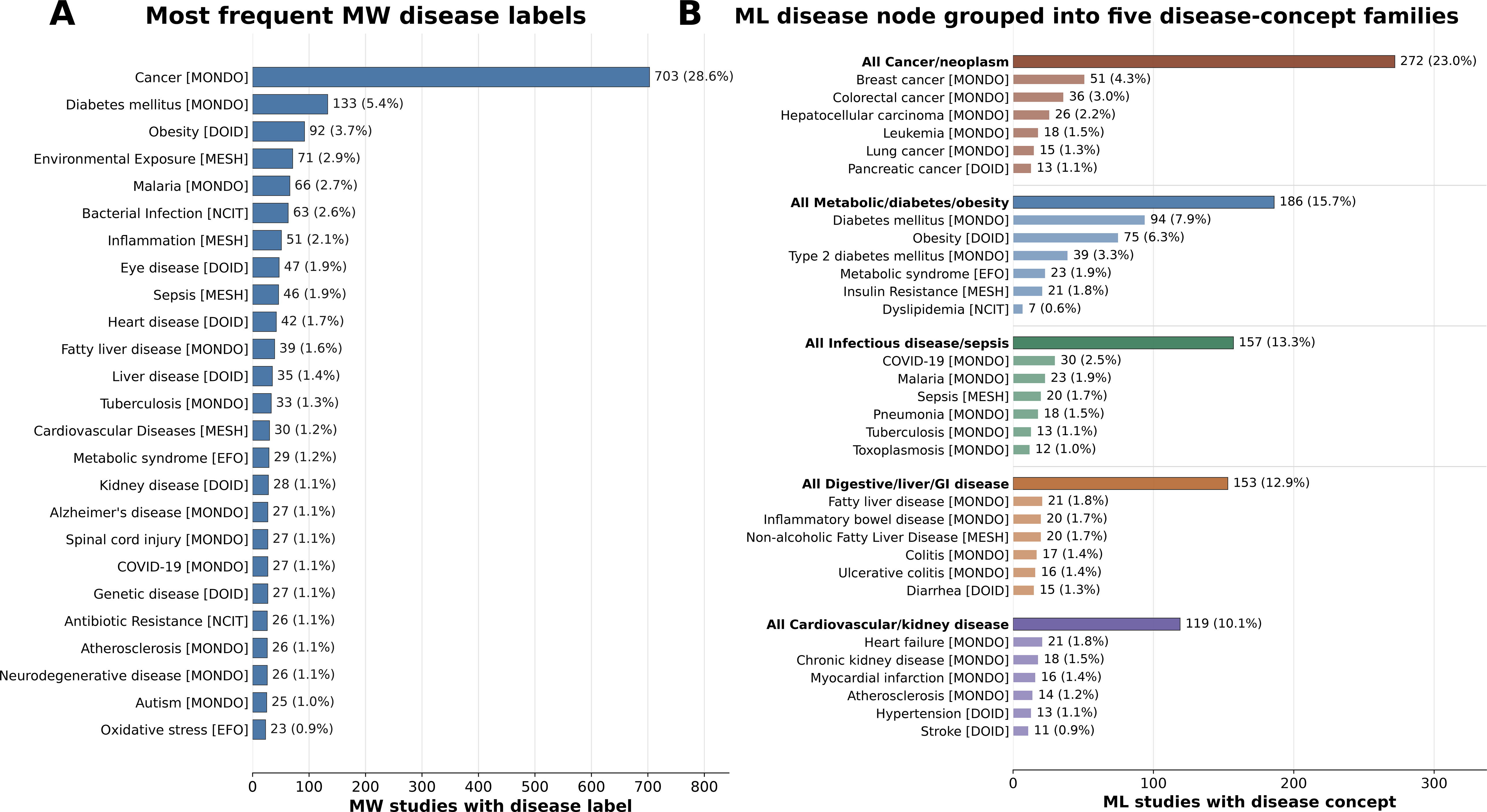
Disease-node mapping landscape across MW and ML. A) Most frequent disease labels in MW. B) Disease concepts in ML are grouped into five broad families for display. Only strictly accepted mappings are shown, and the underlying data retain full labels and ontology provenance.

### Analytical technique harmonization reveals a compact shared platform vocabulary across repositories

Analytical technique mapping achieved near-complete study-level coverage across both repositories (6,826/6,827 studies), reflecting the relatively limited vocabulary used to describe metabolomics assay platforms. A key structural consideration is that these are assay-level rather than study-level attributes; many studies contained multiple assay records, representing genuine multi-platform designs^21,22^ (Figure 3A). Collapsing each study into a single technique label would mask valid within-study heterogeneity, particularly given that combining complementary platforms increases metabolite coverage and the likelihood of detecting metabolites of interest relative to single-technique approaches^22–25^. Mappings were therefore preserved throughout the assay. Because MW and ML structure their assay metadata differently, term extraction followed repository-specific evidence rather than a single unified rule (Figure S4B). In ML, analytical techniques were recovered primarily from the "Assay Type Label" field, with platform, filename, and study design descriptor fields used only for generic or ambiguous entries. In MW, the node was reconstructed from the “/analysis” REST endpoint, where "chromatography_type" provided most LC/GC/CE evidence, supplemented by MS, detector, instrument, and NMR fields for specific methods (Supplementary Methods 3). The resolved vocabulary was compact and well-shared. LC-MS dominated both repositories, followed by GC-MS and NMR (Figure 3B). MW mapped to 15 unique IRIs and ML to 17, with nine shared, yielding a Jaccard overlap of 39.1%, which is substantially higher than for more open-ended biological nodes (species, sample source, or disease). These shared terms covered 98.6% of MW studies and 97.9% of ML studies for cross-repository linkage. Most mappings resolved to CHMO, which covers the major metabolomics platforms; secondary ontologies (TFO/TransformON, PSI-MS) were used only where CHMO lacked a suitable term (Table S8; Figure S4A). Multi-IRI records reflected genuine multi-platform study designs rather than mapping ambiguity, with LC-MS plus GC-MS the most common combination (Figure S4C). Low-frequency modalities, including GC-FID, LC-DAD, GCxGC-MS, DI-MS, FIA-MS, and MS imaging, were retained as distinct terms rather than collapsed into generic MS or LC-MS (Figure S4D).

**Figure 3:**
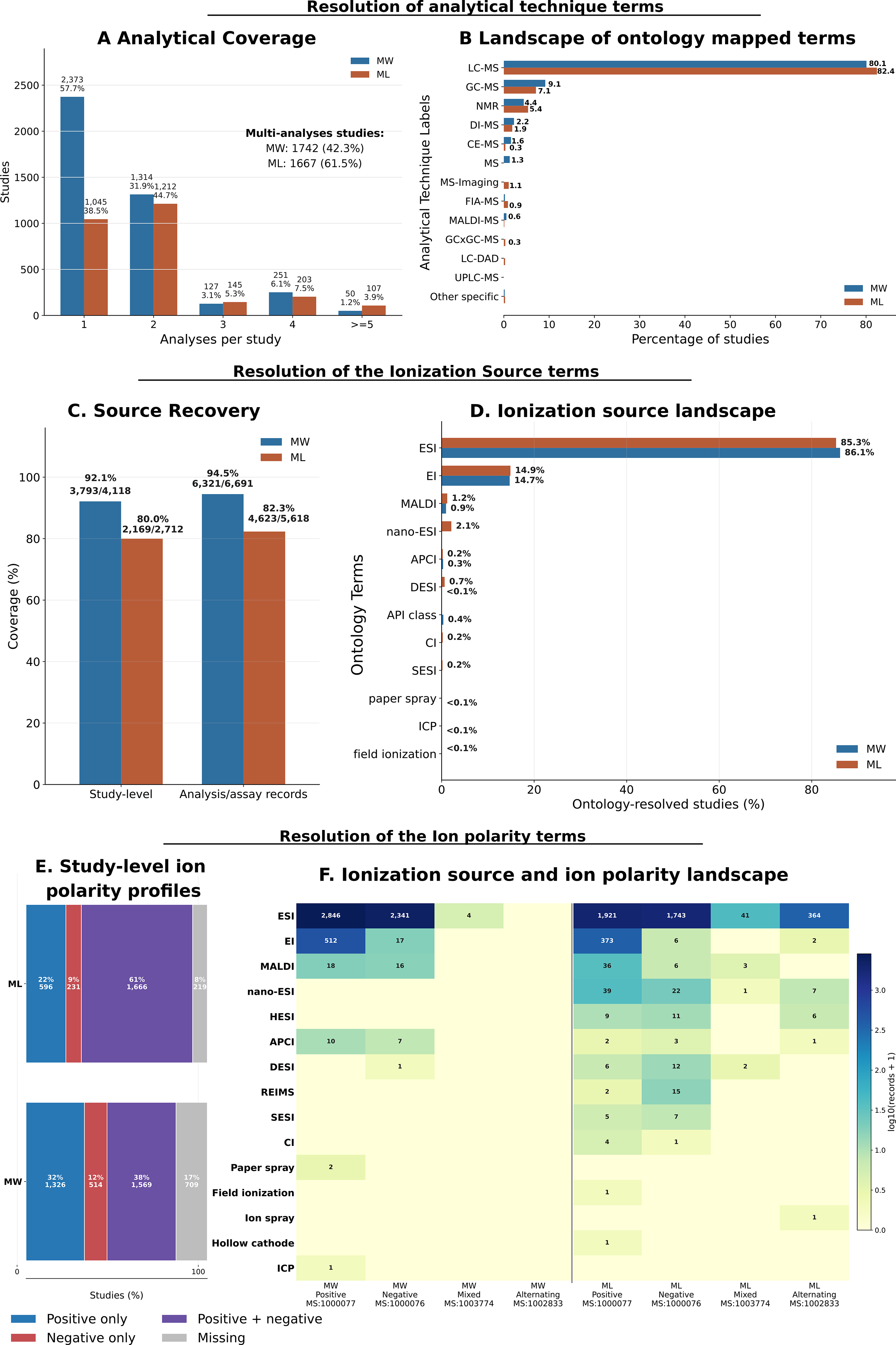
Analytical technique, ionization source, and acquisition polarity harmonization across Metabolomics Workbench and MetaboLights. A) Number of analyses per study after extraction. C) harmonized analytical technique labels. C) ionization-source recovery at the study and record level. D) resolved ionization-source terms. E) study-level ion-polarity profiles. F) combined ionization source and ion-polarity landscape. Across both repositories, LC-MS, GC-MS, NMR, and ESI were the most common patterns, while lower-frequency terms were retained as distinct categories.

NMR records were refined within the analytical technique node rather than treated as a separate acquisition node. All NMR records received the broad CHMO nuclear magnetic resonance spectroscopy term; specific CHMO terms were added only where record-level evidence supported the nucleus, dimensionality, pulse sequence, or experiment family (Table S9; Figure S3(ii)). Refinement covered NMR records in 288 MW and 216 ML studies, including mixed NMR and non-NMR designs. Specific-mapping yields were 95.6% in MW and 99.3% in ML, capturing major experiment families including 1D/2D NMR, 1H/13C NMR, NOESY, CPMG, HSQC, HMBC, COSY, TOCSY, J-spectroscopy, HRMAS, and continuous-wave NMR (Figure S3(ii)). Records lacking sufficient evidence for nucleus, dimensionality, or experiment family were retained under the broad NMR term; acquisition details useful for reuse but unsuitable as ontology terms were stored as raw metadata (Supplementary Methods 3).

### Mapping the ionization source and ion polarity captures high-coverage MS acquisition metadata

Ionization-source mapping was more conservative than analytical-technique mapping because it required direct evidence of the ion-generation mechanism. ML had a larger unresolved fraction than MW. MW studies typically reported the ionization source explicitly through the "ms_type" field, whereas many ML assay files deposited “LC-MS” or “GC-MS” as the analytical technique without a corresponding ion source field. Unresolved ML records, therefore, reflect absent or highly specific source metadata rather than failed normalization (Figure 3C; Figure S5A–B). This makes the ionization source a more evidence-sensitive metadata layer than the analytical technique. The mapped ionization-source landscape was dominated by electrospray ionization (ESI)^23^, followed by electron ionization (EI), consistent with the prominence of LC-MS and GC-MS in public metabolomics (Figure 3D). ESI is the principal ionization source used in LC-MS metabolomics.^22–24,26^ It is well suited to many polar and moderately polar metabolites. In contrast, EI remains the standard ionization method for GC-MS, where its reproducible fragmentation patterns support spectral library matching, including against resources such as NIST^27,28^. Low-frequency sources, including MALDI, nano-ESI, APCI, DESI, chemical ionization, paper spray, SESI, ICP, and field ionization, were retained as distinct terms. Most mapped studies carried a single ionization source. Still, multi-source studies were present in both repositories, driven mainly by EI plus ESI combinations reflecting parallel GC-MS and LC-MS acquisition (Figure S5C–D). The harmonized source layer contained 14 concepts overall (Table S10), of 8 in MW and 11 in ML. Five terms (ESI, EI, MALDI, APCI, DESI) were shared, giving a Jaccard overlap of 35.7% (Figure S5E). Despite the narrow shared vocabulary, these terms covered 99.6% of MW and 98.0% of ML ontology-resolved studies.

Ion polarity achieved higher coverage in ML than in MW at both study and assay levels (Figure 3E). Positive-only acquisitions were more frequent than negative-only across both repositories. Still, combined positive and negative profiles were especially prevalent in ML and common in MW, consistent with the complementary metabolite coverage of dual-polarity LC-MS acquisition^21,29^. These combined profiles did not arise in the same way across repositories. In MW, “positive+negative” study profiles predominantly reflected separate positive- and negative-mode analysis records within the same study. In ML, the same profile could arise from either separate assay files or a single assay file containing explicit dual- or alternating-polarity evidence. This distinction matters in practice. A study-level positive plus negative assignment can mean one of two things: either the same samples were run twice, once in positive mode and once in negative mode as separate acquisitions, or both polarities were acquired within a single run through polarity switching. These are analytically distinct experiments, and collapsing them into a single study-level label would obscure that distinction. Retaining the assay-level polarity structure preserves this information, which is relevant because positive and negative modes detect chemically distinct metabolite classes and often require different instrument settings^30,31^. The ionization source and polarity heatmap confirmed ESI as the dominant source-polarity context in both repositories, with both positive and negative ESI modes well represented (Figure 3F). EI was the next most frequent source, but was predominantly associated with positive-polarity records. Detector-only and non-MS records, including NMR, MR imaging, LC-DAD, and GC-FID, were classified as not applicable and excluded from polarity mapping. A small set of ML assay records contained scan-polarity field values that were not valid polarity terms and mainly contained metadata errors, such as resolution settings or mass ranges. These were retained as unresolved (Tables S12 and S16). Obvious single-token spelling errors were normalized and documented in Table S13.

Ion polarity was harmonized against a PSI-MS-only vocabulary (Table S14-S15). MW resolved to three unique acquisition-polarity IRIs and ML to four, with a vocabulary-level Jaccard overlap of 75%. All mapped MW studies overlapped with ML through shared IRIs, while 2,245 of 2,493 mapped ML studies overlapped with MW. The remaining 248 ML studies carried alternating polarity acquisition (MS:1002833), a PSI-MS term absent from MW.

### Separation-method harmonization resolves analytical acquisition diversity across repositories

Separation-method mapping recovered a high-coverage analytical layer across both repositories while preserving assay and analysis record structure. The resolved landscape was dominated by LC-based methods, particularly reversed-phase LC and HILIC^32^, with GC forming the next major class. Lower-frequency categories captured electrophoresis, normal-phase LC, flow-injection analysis, direct infusion, ion-exchange and ion-pair chromatography, two-dimensional GC^33^, supercritical-fluid chromatography, and explicit no-chromatographic-separation contexts. Broad LC platform descriptors, such as LC, HPLC, and UHPLC, were retained as a separate category rather than reassigned to reversed-phase LC or HILIC, because the deposited metadata did not support a more specific mode assignment. Hence, forcing specificity here would misrepresent the original deposit (Figure 4A). Higher-level grouping of canonical separation method terms is provided in Table S21. At the study level, Figure S6C aggregates unique mapped separation terms per study and therefore reflects multi-method study designs. A study with 5 RP-LC analyses and 1 HILIC analysis is represented the same way as a study with 1 RP-LC and 1 HILIC analysis: both are combined into the study-level combination RP-LC + HILIC. One MW record (ST000163/AN000255), for instance, deposited "Anion exchange/Reversed phase" as a single field and was retained as two separate assignments: ion-exchange chromatography (CHMO:0001014) and reversed-phase LC (CHMO:0001050).

**Figure 4:**
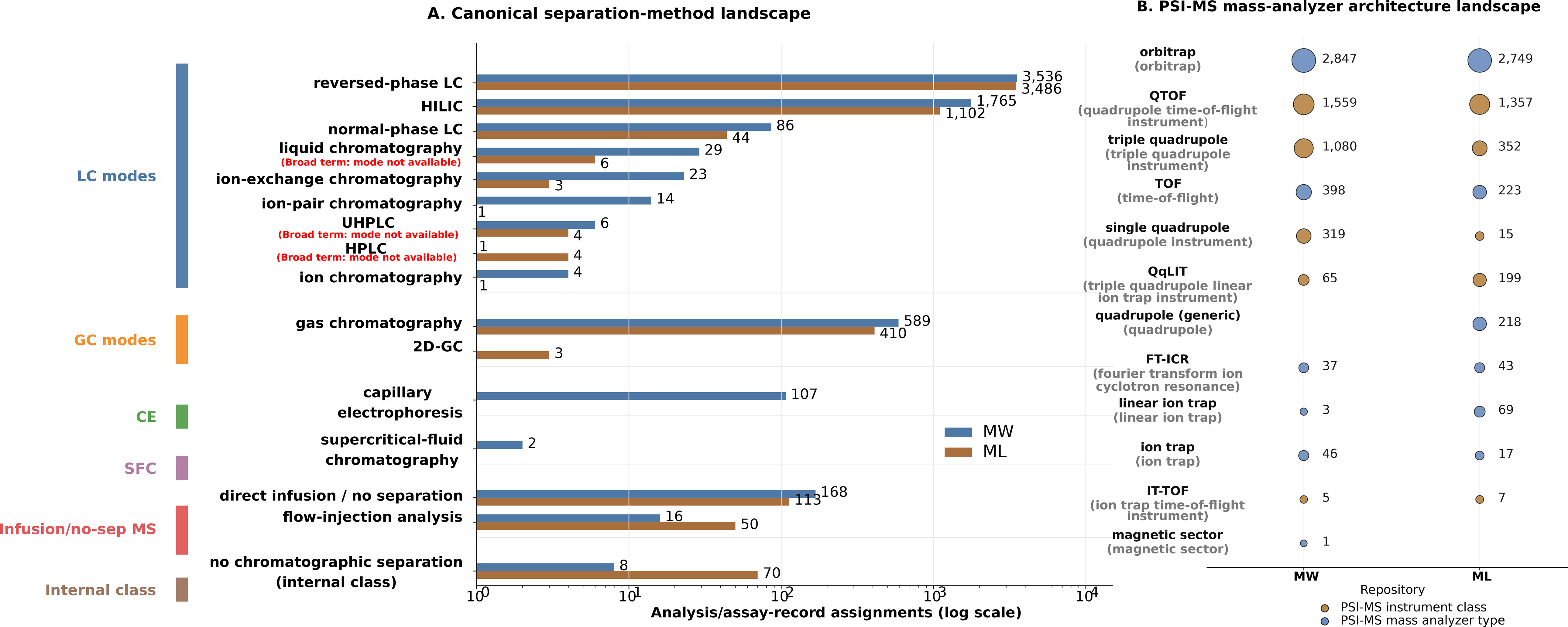
A) Canonical separation-method landscape across MW and ML. Bars show mapped analysis/assay records for each normalized separation-method category on a log scale. Bars omitted where n = 0, for MW 2D-GC or ML’s "capillary electrophoresis" and "supercritical-fluid chromatography". B) PSI-MS mass-analyzer architecture landscape: Bubble matrix showing record-level frequencies of mapped mass-analyzer terms in MW and ML. Bubble size is proportional to the number of mapped analysis/assay records, and color indicates the broader analyzer family. The terms in brackets represent the harmonized labels derived from OLS4.

Mappings resolved primarily to CHMO. Direct infusion was mapped to PSI-MS infusion (MS:1000060) because it represents a defined MS sample-introduction mode rather than a chromatographic or electrophoretic separation. Assays with explicit no-chromatographic-separation evidence, including MALDI-MS imaging, DESI imaging, paper spray MS, PTR-MS, ICP-MS, and SIMS-based workflows, were assigned to a curated internal class, "no_chromatographic_separation", without an IRI. This kept them as valid acquisition contexts and distinguished them from records with genuinely missing separation metadata (Table S20). Primary separation fields accounted for most mappings while secondary fields rescued an additional 48 MW analyses and 237 ML assays that would otherwise have remained unresolved (Figure S6A), utilizing information on column models, chromatography instruments, technology platforms, filename evidence, and study-design descriptors (Figure S6B). The raw-to-canonical mapping is visualized in Figure S7. MW resolved to 14 unique ontology IRIs and ML to 13, with 12 shared, giving a vocabulary-level Jaccard overlap of 80.0% (Table S20). At the study level, 3,752 of 3,800 MW studies and 2,476 of 2,479 ML studies with strict IRI separation overlapped across repositories. The small repository-specific remainder reflected genuine method differences: 48 MW studies were CE-only or SFC-only, and 3 ML studies were 2D-GC-only, none of which have counterparts in the other repository. Studies excluded from the strict IRI overlap carried only the internal no-chromatographic-separation class, remained unresolved after manual review, or lacked an extractable separation term entirely.

### PSI-MS harmonization reveals a shared high-resolution mass-analyzer landscape across repositories

Orbitrap and QTOF dominate the landscape of mapped mass analyzers in both repositories. Quadrupole-based instruments come next, then a shorter tail of TOF, ion-trap, FT-ICR, IT-TOF, and magnetic-sector entries (Figure 4B). Terms were grouped into five families for reporting, but the underlying node keeps the original study-level PSI-MS analyzer and instrument-class mappings (Table S23). Primary analyzer fields provided most of the mappings. Assay-level recovery reached 99.37% in MW (6,354/6,394) and 99.18% in ML (5,226/5,269) (Figure S8A–B). MW’s primary field, "ms_instrument_type," alone supplied 6,329 of those records (99.6% of the MW total). Secondary instrument-name fields added only 25 records, each tied to a specific model. This led to mappings such as Thermo LTQ Ill linear ion trap, Waters Synapt G2 Ill QTOF, Shimadzu LCMS-IT-TOF Ill IT-TOF. In contrast, the primary ML field, "Parameter Value[Mass analyzer]," covered 4,416 records (84.5%), but secondary instrument-model fields were the only usable evidence for another 810 (about 15%), reflecting sparse or generic mass-analyzer parameters in ML metadata.

MW and ML each resolved to 11 unique analyzer terms, 10 of which were shared across a union of 12 (Jaccard 83.3%). The two repository-specific terms are real instrument differences, not mapping gaps. ML alone contributed a broad quadrupole class (MS:1000081; 175 studies), derived from records that only reported generic quadrupole text. MW maintained higher resolution at the single-quadrupole level (287 studies vs. 15 in ML) and a larger presence of triple-quadrupole scans (751 vs. 232). At the same time, ML had more QqLIT assignments (117 vs. 42). MW also held the only mapped magnetic-sector record (MS:1000080), from a single study (ST000565) using a Thermo Element 2 HR-ICP-MS instrument, where the analyzer term was recovered from a "Sector Field" annotation using our pipeline. FT-ICR was present in both repositories at a similar scale (30 MW studies; 27 ML studies), and IT-TOF remained rare (4 MW studies; 2 ML studies). The full PSI-MS mapping, analyzer-family grouping, and repository presence/absence patterns are in Table S23 and Figure S8C.

Studies with more than one analyzer were uncommon (MW 3.80%, ML 6.81%; Figure S8D) and occurred when different assays within a study mapped to different analyzer terms, indicating mixed instrument designs rather than ambiguous mapping. The most frequent analyzer pair overall was Orbitrap with triple quadrupole (49 studies combined: 28 MW, 21 ML), whereas repository-specific top pairs were QTOF with single quadrupole in MW (36 studies) and QTOF with triple quadrupole in ML (26 studies). QTOF is widely used to scan broadly for unknown compounds^34^, and its complementary use with QQQ for pathway profiling is well established^35^. Genuine multi-analyzer strings, in which a single deposited term from a given assay named more than one architecture, were rare: 29 records total (6 MW, 23 ML), most commonly QqLIT plus QTOF in ML (7 records). What remained unmapped was mostly generic MS technique labels, manufacturer-only strings, chromatography or LC platform names, detector-only text, software names, and instrument-setting strings.

### Metabolite identity resolution and differential abundance across repositories

#### RefMet mapping differs between repositories in metabolite-name resolution

RefMet coverage was assessed in studies with at least one structurally valid deposited abundance matrix (4,695 studies: 3,092 MW and 1,603 ML). Coverage was calculated at the level of distinct deposited metabolite-name literals, that is, the exact text strings as deposited, rather than the number of distinct chemical entities, because the same compound can appear under multiple synonymous names or spelling variants across studies. Counting each such literal once, 27.9% resolved to a RefMet identity overall. Resolution was higher in MW than in ML at the unique-deposited-name level: 64.6% versus 12.5%.(Figure 5A; Figure S12A-B). These figures, therefore, reflect how readily deposited names map to RefMet, rather than the size of the underlying chemical space in either repository.

**Figure 5:**
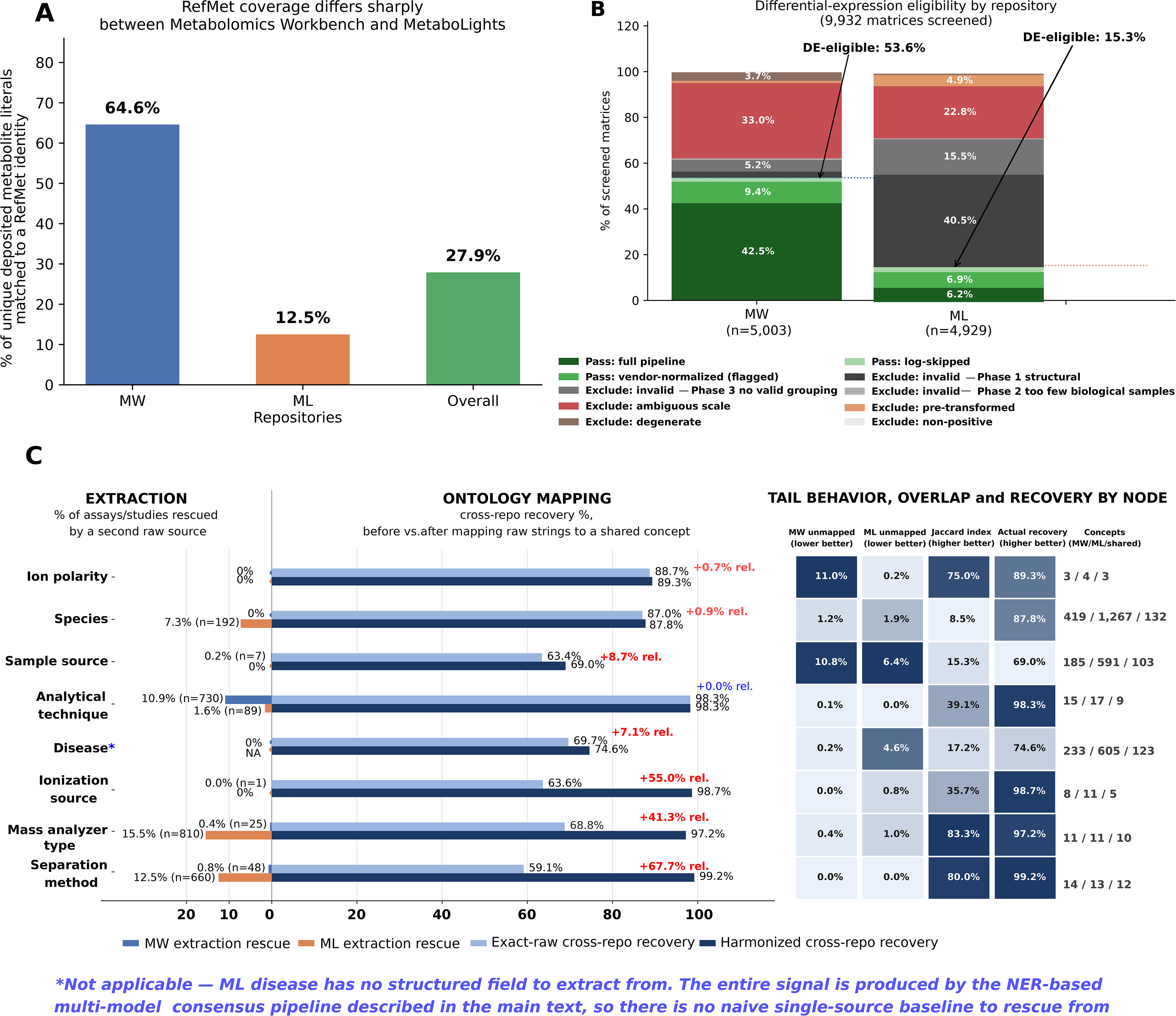
Repository differences in metabolite identity resolution and differential-expression eligibility. A) RefMet coverage at the unique deposited-name level among studies with at least one structurally valid abundance matrix. B) differential-expression eligibility after screening 9,932 matrices, with arrows marking the overall DE-eligible fraction in each repository. C) two harmonization steps per node. The left side shows extraction rescue, the fraction recovered only by a second raw evidence source. The right side shows cross-repository recovery before and after ontology mapping, where exact-raw counts studies found by searching the literal deposited text, and harmonized counts studies found by searching the shared ontology term. Relative gain is calculated as (harmonized−exact-raw)/exact-raw×100. In ML, there is no naive baseline because there is no structured field. D, tail behavior, vocabulary overlap, and recovery by node. MW/ML unmapped is the percentage never resolved to a concept, Jaccard index is the shared share of the combined concept vocabulary, and actual recovery is the percentage of usable studies found by a shared-concept search.

Manual review of the unmatched literals suggested several reasons for this gap (Figure S12C). A representative set of examples from each repository is shown in Table S31. The largest group, 34.9%, consisted of entries that were not compound identifications at all, including untargeted feature labels, m/z values, and arbitrary feature codes that the depositor had never assigned chemical identities. A further 12.1% were lipid entries recorded in detailed acyl-chain shorthand that did not exactly match the simpler standardized RefMet naming format, indicating a naming-detail mismatch rather than a failure to identify the lipid class. Long systematic or IUPAC-style names made up 7.2% and may be recoverable through alternative common synonyms. Another 8.6% consisted of numeric-only entries, catalog or screening-library codes, and ambiguous multi-candidate entries written as slash-separated mixtures. Peptide sequences accounted for a further 4.9%, while 1.9% reflected smaller formatting-related causes, including pipe-delimited strings, RT-suffixed entries, and others. The remaining 30.5% were entries corresponding to named compounds that still could not be resolved after these categories were taken into account. This group is reported conservatively as unresolved rather than being treated as a confirmed gap in RefMet coverage. Where repository-specific breakdowns were possible for the clearly defined categories (Figure S12D), ML’s unmatched literals were dominated by feature-level identifiers, which accounted for 40.8% of ML’s unmatched pool, compared with 0.1% in MW. MW, by contrast, showed greater enrichment for lipid-shorthand entries, which made up 25.2% of its unmatched literals, compared with 9.9% in ML.

#### Structural and metadata gaps jointly limit differential-expression eligibility across repositories

Differential expression eligibility was assessed from the metadata across 9,932 screened matrices (5,003 from 3,216 MW studies; 4,929 from 2,633 ML studies), using a seven-phase workflow that included structural validity checks, biological-sample filtering, factor-based grouping, and classification of the declared measurement scale (Supplementary Methods 7: Quality-tier classification and disposition). Across both repositories, 34.6% of screened matrices (3,433 of 9,932) qualified for testing. Eligibility differed substantially by repository: 53.6% of MW matrices were eligible, compared with 15.3% of ML matrices (Figure 5B). Among eligible matrices, most MW matrices (79.4%) entered the full preprocessing workflow of imputation, log2 transformation, and median centering. In ML, 44.9% of eligible matrices had already been normalized before deposition; these matrices were analyzed through the same downstream pipeline and were retained with a flag in the output.

The difference in eligibility arose from different failure patterns in the two repositories (Figure S13A-B). In MW, most exclusions were due to ambiguity in the declared abundance scale, such as missing or unresolved units or unclear protocol text, even when the underlying data could otherwise be parsed and structured correctly. In ML, the largest source of exclusion was structural failure, followed by ambiguous scale information and failed factor grouping. Thus, the lower eligibility in ML does not reflect the same underlying problem as the higher eligibility in MW. Confounding was common in the full-joint differential analysis: 66.5% of pairwise contrasts differed in two or more experimental factors, and these contrasts accounted for 68.1% of all FDR-significant calls (Figure S13C). In parallel, 20.0% of all tested contrasts included at least one group at the minimum replicate threshold of n=2, with 7.6% at n=2 in both groups (Figure S13D).

### Ontology harmonization converts heterogeneous repository metadata into shared cross-repository query terms

The efficiency of harmonization across both databases was driven by two independent steps: recovery of usable raw metadata and mapping of those values to shared ontology concepts. These steps did not contribute equally across nodes. Across all eight metadata nodes, our extraction pipeline alone rescued a meaningful share of records that a naive single-field lookup would have missed entirely (Table S32; Figure 5C, left titled “EXTRACTION”). In ML, the largest rescue gains came from mass analyzer type, where an instrument-name fallback recovered 15.5% of analyses, and separation method, where secondary fields recovered 12.5%. In MW, the main rescue was the analytical technique node, which recovered 10.9% of analyses that would otherwise have remained unassigned. The disease node in ML is not directly comparable on this metric because there is no structured disease field to rescue from. It is assembled through a multi-model NER pipeline instead. Within that pipeline, keeping single-model evidence after review recovered 360 additional studies (20.4%) beyond the strict two-of-three consensus, but this reflects the consensus threshold rather than extraction rescue. By contrast, ion polarity in both repositories, species in MW, sample source and ionization source in ML, and disease in MW required no rescue because each already came from a single reliable source.

Once raw values were recovered, ontology mapping determined whether they could be matched across repositories, and this is where most of the gain came from (Figure 5C, right, titled “ONTOLOGY MAPPING”). We report the gain both as percentage points and as a relative gain (Figure 5C legend), showing that harmonized retrieval outperformed raw-string matching. The largest gains came from the separation method (59.1% to 99.2%, +67.7% relative), the ionization source (63.6% to 98.7%, +55.0% relative), and mass analyzer type (68.8% to 97.2%, +41.3% relative). These are also the nodes where extraction on MW was already close to complete, so mapping mostly resolved vocabulary differences rather than missing data. On ML, two of these three (separation method and mass analyzer type) still needed substantial extraction rescue (12.5% and 15.5%) before mapping could help.

Ionization source, on the other hand, needed almost no extraction rescue in either repository (0.02% MW, 0% ML), unlike the separation method or mass analyzer type. Sample source improved modestly through mapping, along with a small MW extraction rescue. Ion polarity changed little on both fronts, since both repositories were already close on that term before harmonization. Species showed little mapping gain (+0.9% relative), though it had a real, separate extraction contribution: 192 ML studies (7.3%) recovered through an API fallback that mapping did not need to compensate for. The analytical technique showed no mapping gain, indicating that extraction, and not mapping, was the primary contributor for that node. Disease showed a modest improvement in mapping, but it is not directly comparable because ML lacks a structured disease field. Both repositories also share relatively little disease vocabulary (123 of 715 distinct concepts, 17.2%), indicating that MW and ML are studying different diseases rather than reflecting a mapping shortfall, unlike separation method, ionization source, and mass analyzer type, where 36–83% of the vocabulary is already shared between the two repositories and mapping recovers nearly all of it.

Across all eight metadata nodes, the unresolved studies and terms make up only a small tail of the data (Figure 5D), limited mostly to repository-specific instrument models, disease subtypes, and a few edge cases. Most studies in both repositories map to a shared vocabulary that can be searched across databases, showing that combining extraction with ontology mapping produces real corpus-wide harmonization rather than a partial fix.

Harmonization also improved discovery within each repository, even without cross-repository matching (Figure S14). For each node and repository, we compared the share of studies found using raw text versus harmonized concepts within that repository. The difference was small overall, ranging from no change to an increase of 8.1%, with a median increase of 0.4%, because studies in the same repository already tend to use similar vocabulary. Even so, harmonization reconnected 514 studies across all 16 node-repository combinations that raw-text search would have missed, including 202 of 2,483 ML studies for mass analyzer type, where findability rose from 91.9% to 100% because free-text instrument names split the same concept across multiple strings. Notably, the nodes with the largest within-repository gains (mass analyzer type, separation method, sample source, disease) are the same free-text, unconstrained-vocabulary fields that also showed the largest cross-repository gains, indicating harmonization resolves the same underlying vocabulary fragmentation at two different scales.

## Discussion

Public metabolomics repositories suffer from incomplete, inconsistent, and variably structured metadata, especially for biological context and chromatographic annotation^6,14,36,37^. These limitations reduce discoverability and make cross-study reuse difficult even when datasets are publicly available. Our framework, HARMONY (Harmonized Annotation and Resolution of Metabolomics Ontologies across Node-mapped repository data), addresses this problem by separating the recovery of usable metadata from the mapping of those values onto shared ontology concepts, thereby converting heterogeneous deposition practices into a common query layer without discarding the original evidence.

This separation between metadata recovery and ontology mapping is most evident in the disease node, where the bottleneck is not vocabulary alignment but the recovery of disease context from records with different structures. Disease normalization across clinical vocabularies is a well-studied problem^38^, and prior work has shown that raw disease names can be aligned to structured terminologies such as SNOMED CT^39^ and Disease Ontology^40^ However, harmonization accuracy and hierarchical coverage still vary depending on which terminology is used and how granular the source terms are^41^. We face a similar problem while building our pipeline. In MW, disease is exposed as a study-level endpoint. In ML, the same information must be assembled from study summaries, design descriptors, factor metadata, and ISA-Tab fields. In that setting, ontology lookup alone cannot recover a concept that has not first been extracted from unstructured text and then checked against the study design. Hence, we built a dedicated named-entity recognition and consensus-based extraction workflow with extensive manual validation rather than treating it as a simple extension of the shared ontology layer used for the other nodes. The lower disease recovery observed in ML, therefore, reflects the degree to which disease context is structured at deposition, rather than a shortcoming of ontology mapping itself. In other words, extraction is the first bottleneck. A disease term cannot be mapped if the repository does not expose it clearly enough.

Once extraction is separated from mapping, the value of the ontology layer becomes clear. In nodes such as separation method, ionization source, and mass analyzer type, extraction was already close to complete, but MW and ML still used different labels for the same concepts. Mapping did not recover missing information; it made those values searchable across repositories by converting them into shared terms. Reporting compliance alone does not capture this, because two records can both satisfy minimum reporting requirements^6,12^ and remain invisible to each other if they describe the same concept with different strings. The resulting reduction from many raw strings to a smaller concept set is a direct measure of curation effort, especially in analytical nodes where branded instrument names and shorthand notations mask a much smaller underlying space. Related efforts, such as MetabolomicsHub^42^ are building a shared data model and a common search layer across MetaboLights, Metabolomics Workbench, and GNPS^43^. HARMONY instead focuses on retrospective ontology mapping for a fixed set of 9 study-level nodes in MW and ML, while preserving deposited evidence and linking studies to RefMet metabolite identities and searchable differentially abundant metabolites.

The HARMONY web platform turns this harmonized layer into a practical search system. Instead of asking users to browse ontology tables, it stores the mapped concepts, raw deposited terms, RefMet names, accessions, study metadata, and evidence links in a local indexed database. Searches, therefore, run on a stable version rather than on repeated live calls to MW, ML, or OLS4. Users can search by ontology term, accession, metabolite name, study term, or broader text. Each result keeps links to the original repository record, so the source evidence remains visible. Taken together, HARMONY is not only a metadata-normalization pipeline but also an operational model for connecting public metabolomics repositories without erasing the evidence and uncertainty present in the original deposits. Future updates will expand ontology coverage, improve disease and metabolite-name resolution, refresh the corpus as MW and ML continue to grow, and extend the framework to additional resources such as GNPS.

## Methods

The harmonization nodes were selected to capture the main dimensions along which two deposited studies can be compared. Species, sample source, and disease define the biological setting. In contrast, the analytical technique, separation method, ionization source, ion polarity, and mass analyzer type define the measurement setting that shapes metabolite detection and cross-study comparability. For ontology-backed nodes, OLS4-based lookup^44^ provided programmatic access to candidate terms, labels, synonyms, and identifiers during mapping. Ontology assignments followed a conservative evidence hierarchy rather than blind string matching: exact labels, exact synonyms, repository-provided accessions, and deterministic rules were accepted only when consistent with field provenance and node scope, while non-exact, ambiguous, or high-risk candidates were manually reviewed against raw text, source field, ontology context, and exclusion rules; valid terms lacking a safe match were retained as raw literals rather than forced into an incorrect IRI. RefMet^45^-linked metabolite identifiers and differentially abundant metabolites (DEMs) connect these study-level descriptors to metabolic profiles. Together, these nine nodes support cross-repository search while preserving the original deposited terms from which each mapping was derived.

### Node term extraction and ontology mapping

#### Species

Species labels from MW and ML were mapped to NCBI Taxonomy^19^ (NCBITaxon). Each mapped taxon was represented by an Internationalized Resource Identifier (IRI), corresponding to the web-accessible OBO-format form of an ontology term, e.g., http://purl.obolibrary.org/obo/NCBITaxon_{id}. NCBITaxon was used as the sole target vocabulary for this node because it spans biological taxa from kingdom to strain and is already used as a reference vocabulary within both repository metadata. Raw MW species strings were extracted from the study-level MW species endpoint. For ML, species annotations were first parsed from ISA-Tab sample tables using the “Characteristics[Organism]” column, along with the adjacent “Term Source REF” and “Term Accession Number” fields. Accessions already pointing to NCBITaxon were canonicalized directly to OBO-format IRIs. To avoid false taxon assignments from blanks, quality-control rows, environmental matrices, or placeholder accessions, ISA-Tab organism rows were retained only when the term source or accession supported an NCBITaxon mapping. For ML studies with no organism annotation recovered from ISA-Tab parsing, we performed a second pass using the MetaboLights organisms REST API (GET/ws/studies/{id}/organisms). When multiple taxa were returned for a study, they were split into separate entries, non-biological values were removed, and only terms confidently supported by NCBITaxon were retained. The source of these mappings was recorded as either recovered from the MetaboLights organisms API or directly extracted from the ISA-Tab sample metadata. Species labels were then harmonized to NCBI Taxonomy using an evidence-based, rank-aware matching criterion that preserved the original taxonomic resolution, from broad groups and genera to species, subspecies, varieties, strains, and isolates (Supplementary Methods 1). Repository-provided NCBITaxon accessions were canonicalized directly, whereas raw organism strings were resolved through NCBITaxon-restricted OLS4 lookup using exact label, exact synonym, and taxonomically constrained fallback matches only. Multi-organism records were split into separate aligned taxon entries, and ambiguous aliases or strain-level mappings were excluded or manually corrected to prevent ambiguous species assignments.

### Sample Source

Sample-source terms were extracted from repository metadata using repository-specific fields. For ML, terms came from ISA-Tab sample files (s_*.txt) via the "Characteristics[Organism part]" column and the adjacent "Term Source REF" and "Term Accession Number" fields. For MW, terms were drawn from the REST "source" and "factors" endpoints, with the mwTab "COLLECTION.SAMPLE_TYPE" field used as a fallback. Raw strings were retained for provenance, and normalized strings were used for lookup. Composite fields were split only where delimiters separated genuinely distinct source materials (Supplementary Methods 2; Tables S1–S2). Because sample-source metadata spans tissues, biofluids, cell types, cell lines, blanks, disease context, analytes, procedures, organisms, environmental and food materials, OLS4 searches were restricted to a defined ontology set. Primary targets were UBERON, BTO, CL, CLO/Cellosaurus, NCBITaxon, ENVO, FOODON, and PO; OBI, EFO, and SNOMED CT were accepted only after semantic validation. Disease, chemical, assay, and procedure ontologies were excluded as primary mappings (Table S3). Candidates were scored on label or synonym agreement, ontology tier, specificity, and ancestor evidence (Table S4), then classified into source categories by ontology and parent term (Table S5). Terms describing blanks, QC samples, diseases, analytes, or procedures were retained as raw text but left unmapped. Each retained record carried the raw value, normalized value, split value, mapped IRI, mapped label, ontology, mapping status, evidence type, source category, parent terms, and review notes reflecting the evidence supporting each decision.

### Disease

Disease was treated as a study-level node, with multiple terms retained when the deposited metadata supported more than one concept. For MW, disease terms were taken from the study-level REST disease endpoint, split when multiple terms appeared in a single field, and retained with the original raw value for provenance (Supplementary Methods 6). ML does not have a structured disease endpoint, so disease context was recovered from study summaries, study design descriptors, factor names, factor types, and the Factor Value[] columns in ISA-Tab files (Supplementary Methods 6; Figure S9; Table S25). Disease mentions were identified independently by three models: a BioBERT^46^-based NER model and two instruction-tuned biomedical language models, MedGemma-27B^47^ and GPT-OSS-20B^48^. Decoding was deterministic, and the prompts were constrained so that the models returned only disease terms directly supported by the text. After normalization, terms supported by at least 2 models were considered high-confidence. Single-model candidates were not discarded; they were kept as lower-confidence evidence for review.

All accepted terms were deduplicated, normalized, and mapped through OLS4, using exact label or synonym matches whenever possible. Each candidate was then checked for semantic consistency before inclusion. Terms were kept only when the deposited metadata clearly supported a disease or clinical context. Organism-only labels, non-disease biological or physiological context, method-development studies, environmental stress, diet and exposure studies, microbial ecology, and plant pathology traits were excluded unless the record also showed clear evidence of disease, infection, pathological challenge, or resistance context. Broad labels such as “cancer” or “infection” were retained only when no more specific supported term was available. Valid disease literals without a safe ontology match were kept as raw text rather than forced into an incorrect IRI.

Accepted terms from both repositories were mapped through OLS4 against MONDO, MeSH, DOID, NCIT, and EFO (Table S26). Exact label and exact synonym matches were preferred, and fallback matching was used only when the candidate closely matched the deposited concept. Secondary ontology mappings were retained when they better reflected the metadata than a forced disease ontology term. Each final disease record preserves the mapped IRI and label, ontology, synonyms, parent terms, hierarchy summary, broad fallback mappings, valid unmapped literals, excluded context terms, and evidence-layer status.

### Analytical Technique Extraction and Mapping Methods

#### MetaboLights

Analytical technique labels in ML were derived from assay-level MetaboLights REST metadata (isaInvestigation.studies[].assays[].comments[]), with each assay object treated as a separate analytical record (Figure S3(i)). The primary evidence source per assay was the “Assay Type Label” comment (the badge label shown in the MetaboLights UI), with “Study Assay Technology Platform” (REST field technologyPlatform), assay filename, broad technology type, and study design descriptors used as fallback when the label was generic, missing, or required refinement (Supplementary Methods 3). Raw strings were preserved for provenance and normalized only for matching. Node-specific rules prevented specialized techniques such as LC-DAD, GC-FID, GCxGC-MS, CE-MS, DI-MS, FIA-MS, MALDI-MS, MS imaging, NMR, and MRI from being collapsed into broad LC-MS, GC-MS, or generic MS categories (Figure S3(i)). Accepted terms were mapped to CHMO via OLS4. Secondary ontologies were used only where CHMO lacked a suitable method-level term. Each record retained the raw term, normalized term, ontology IRI, mapped label, evidence source, aliases, synonyms, and curation notes.

#### Metabolomics Workbench

For MW, analytical technique labels were reconstructed from per-analysis metadata via the study-level REST “/analysis” endpoint. Only analysis-level evidence was used, including broad analysis type, analysis summary, chromatography descriptors, instrument and detector descriptors, MS acquisition descriptors, and NMR-specific fields. Study titles, descriptions, mwTab-derived labels, and metabolite-level fields were excluded, as these are not reliably tied to individual analysis records and can collapse platform-specific qualifiers to generic MS or introduce cross-analysis leakage in multi-platform studies. As with ML, controlled rules prevented detector-coupled methods, NMR, DI-MS, FIA-MS, CE-MS, and source-specific MS modalities from being collapsed into a single generic MS modality. The latter was assigned only when analysis-level metadata could not support a more specific term (Supplementary Methods 3). Multiple techniques per study were retained where evidence indicated distinct analytical platforms. Final terms were mapped to CHMO, with secondary ontologies used only where no suitable CHMO term existed.

#### NMR refinement for both databases

NMR experiment-type refinement was applied at the assay or analysis record level rather than as a separate node (Supplementary Methods 3). MW NMR evidence was obtained from the study-level REST “/analysis” endpoint, which contains NMR-specific fields. ML NMR evidence came from assay-level metadata and pulse-sequence fields in assay files. All NMR records received the broad nuclear magnetic resonance spectroscopy CHMO term, with specific terms added only where direct record-level evidence supported the nucleus, dimensionality, pulse sequence, or experiment family.

### Ionization source and ion polarity terms: Extraction and Mapping Methods

Ionization source annotations were extracted at the assay or analysis record level and kept distinct from analytical technique, ion polarity, mass analyzer, separation method, and detector metadata. For MW, each “/analysis” REST endpoint row was treated as a single record, with “ms_type” as the primary evidence field; analysis summary or instrument descriptors were used only when they explicitly named a source. For ML, the ionization source was extracted primarily from deposited assay metadata files (a_*.txt, fetched via the MetaboLights REST assay-file endpoint) using “Parameter Value[Ion source]”, with the next two fields, “Term Source REF” and “Term Accession Number”, aligned in the column order. Where that column was absent, explicit ionization terms in “technologyPlatform” or source-bearing “Assay Type Label” values (e.g., MALDI-MS, DESI-MS) were permitted as fallback. No ionization source was inferred by assumption. A record required explicit source-level evidence to receive an assignment, and adjacent metadata fields were not used as proxies. For instance, an LC-MS record without a deposited ion source field and required fallback entries was left unresolved rather than automatically assigned ESI (Electrospray Ionization). Accepted terms were mapped to PSI-MS, and secondary ontologies were used only for caveated cases lacking a suitable PSI-MS source term. Missing, generic, detector-only, polarity-only, analyzer-only, and non-MS values were left as null or not applicable.

Ion polarity was extracted at the per-assay level before study-level aggregation, to preserve multi-analysis and multi-assay structures in MW and ML, respectively (Supplementary Methods 4). For MW, polarity evidence came exclusively from the “/analysis/ion_mode” REST endpoint, with each returned analysis object treated as a single polarity record. Blank, missing, or UNSPECIFIED values were left unmapped. For ML, "Parameter Value[Scan polarity]" from assay file metadata was the sole evidence source. Because this field can span multiple rows within a single assay file, all row-level values were inspected before assigning an assay-level polarity class. Files with only positive values were assigned positive polarity, files with only negative values negative polarity, and files containing both were treated as dual-polarity records (Table S11). Explicit polarity-switching or alternating-acquisition terms were also accepted as dual-polarity evidence. PSI-MS served as the primary ontology for the ion polarity vocabulary.

### Separation technique: Extraction and Mapping Methods

The separation method was extracted at the assay or analysis record level, since a single study may contain multiple analytical methods, including different chromatographic modes, direct-infusion assays, or no-separation MS workflows. For MW, the primary evidence field was “chromatography_type” from REST-derived records of the “/analysis” endpoint. When this field was empty or uninformative, fallback evidence was drawn solely from the same analysis row, covering column name, chromatography system, analysis summary, MS type, and instrument metadata (Table S17). Study-level “analysis_type” values were not used to fill missing record-level fields, as broad labels such as “LC-MS” may not reflect the separation method of each analysis record. Records were left unresolved where analysis-level evidence was absent or conflicting. For ML, the main evidence fields were "Parameter Value[Column type]", including suffixed versions, "Parameter Value[CE Instrument]", and "Parameter Value[Column model]". Direct-infusion, flow-injection, and no-separation signals were checked in the assay technology platform, type, and filename fields before broader instrument fallbacks were applied. "Parameter Value[Chromatography Instrument]", technology platform, assay type, assay filename, and study-design descriptors were used only as fallback. Study-design descriptors were accepted only when they contained explicit analytical information, such as direct-infusion MS, UPLC-MS, or DESI imaging (Table S17).

Raw strings were normalized to collapse repository-specific abbreviations, column chemistries, instrument shorthand, and no-separation descriptors into canonical separation-method categories before ontology mapping. The full normalization dictionary is provided in Supplementary Methods 5 (Table S18), and manual validation was performed before freezing the node to prevent false-positive mappings (Table S19). Canonical terms were mapped to CHMO. Direct infusion was mapped to PSI-MS infusion (MS:1000060) as it represents MS sample introduction rather than chromatographic separation. A curated "no_chromatographic_separation" class was retained without an IRI for assays performed without chromatography. Broad LC descriptors, such as LC, HPLC, and UHPLC, were not collapsed into mode-specific terms, such as reversed-phase LC or HILIC, without direct record-level evidence. Records outside the scope of chromatographic or electrophoretic separation (for example, CE-MS or Capillary Electrophoresis) were marked not applicable. Those with conflicting evidence were left unresolved.

### Mass analyzer type: Extraction and Mapping Methods

Mass analyzer type was assigned at the analysis or assay level for MW and ML, respectively. Evidence fields followed a fixed hierarchy. In MW, “ms_instrument_type” was the primary field, with “ms_instrument_name” and “analysis_summary” used as fallback only when they contained explicit analyzer or model evidence. In ML, "Parameter Value[Mass analyzer]" was the primary field, with "Parameter Value[Instrument]", technology platform, assay filename, and assay type label as secondary fallback when they contained explicit analyzer or model information (Table S22). Broad technique labels such as LC-MS, GC-MS, or mass spectrometry were not used to infer an analyzer type. Raw strings were normalized for case, hyphens, spacing, and punctuation before lookup, so that equivalent forms such as “QTOF”, “Q-TOF”, and “q tof” were treated identically (Table S24). Specific analyzer architectures and named instrument models were prioritized over broad labels throughout.

All accepted labels were mapped to PSI-MS. Where a direct analyzer term existed, such as Orbitrap, TOF, ion trap, FT-ICR, quadrupole, magnetic sector, that term was used. For architectures where PSI-MS represents the concept as an instrument class rather than a pure analyzer term, such as QTOF, triple quadrupole, IT-TOF, QqLIT, and single quadrupole, the corresponding instrument-class IRI was used instead (Table S23). After mapping, each assigned PSI-MS accession was checked against the official PSI-MS OBO file to ensure the term is still in use (not obsolete) and that the label we used matches the name in the ontology. Placeholder values, including "-", "Other", "Several", "Unspecified", and "Multiple," were treated as missing unless another field in the same record provided usable evidence. Records were marked as not applicable when no analyzer is expected, including NMR, MR imaging, and detector-only assays.

Records were left unresolved when the deposited text was real but not specific, including generic LC-MS or GC-MS labels, chromatography platform names, software or settings strings, manufacturer-only values, ion-mobility-only labels, and instruments pending a separate policy decision, such as Waters Cyclic MS.

### Metabolite name normalization and derivation of differentially expressed metabolites

Deposited metabolite names for all studies were re-annotated against RefMet. For MW, names were retrieved per study from the REST “/metabolites” endpoint and grouped by analysis ID, with a study-level union of unique names taken across all analyses. For ML, every deposited MAF (m_*.tsv) associated with a study was downloaded individually, since a substantial fraction of ML studies carry more than one MAF (1,473 of 2,712 studies had two or more), and the “metabolite_identification” column from each file was pooled into a single study-level union with duplicates across assays collapsed. Unique names were submitted in batches of 5,000 to the Metabolomics Workbench online name-to-RefMet conversion service (name_to_refmet_new_minIDS.php) via POST request, and names returning a valid RefMet identifier were retained as mapped.

Differential abundance was computed for each analytical matrix, directly from deposited raw metabolite tables. The overall workflow is shown in Figure S11. Matrices for MW came from the analysis-level “datatable”, with “mwTab” used as a fallback when no datatable was available. For ML, matrices were obtained from the per-assay MAF files. Each MW analysis (AN######) and each ML MAF was treated as its own independent statistical unit and was never pooled with other analyses from the same study. Non-biological samples, such as quality control, pooled reference, blank, solvent, internal standard, and calibration-standard rows, were removed before any downstream processing, using a rule-based classifier applied consistently across both repositories (Table S30; Supplementary Methods 7: Non-biological sample removal). This classifier worked on sample identifiers, factor strings, and sample-type fields jointly, rather than relying solely on column-name pattern matching.

Each remaining matrix then had to pass a structural validity check, including successful parsing, containing at least one metabolite row and two sample columns, feature distributions showing sufficient variability, and having at least one strictly positive value, before moving on to the next step of quality-tier classification. Per-sample experimental factors were extracted from each repository’s native metadata and combined into a shared grouping variable. Missing-value placeholders were first converted to empty values to prevent the creation of artificial groups. Technical or administrative factors, such as “batch”, “run day”, and “replicate”, and subject-like identifiers, were excluded from grouping in both repositories. Biologically informative factors were preferred when available. Groups were defined from the full set of retained factors when this yielded at least 2 samples per group. Otherwise, a single primary factor was used. Batch correction was not applied because batch annotations across the corpus were sparse, inconsistent, and often entangled with the biological design (Supplementary Methods 7: Experimental factor parsing and joint-group construction).

A conservative quality-tier assessment was performed on each matrix to determine whether it was suitable for a uniform differential-analysis workflow (Supplementary Methods 7: Quality-tier classification and disposition). Deposited metabolite matrices were reported on mixed numeric scales, including raw intensity, concentration, log-transformed data, fold-change data, etc. Hence, we first screened each matrix to determine the abundance scale, using depositor-reported units and protocol text. The value distribution was used only to confirm or challenge the metadata, because distributions can be suggestive but are not reliable enough on their own to decide the input scale. Ambiguous cases where the metadata and observed values didn’t agree clearly were excluded (Supplementary Methods 7: Conditional processing).

Eligible matrices from both repositories were processed with the same workflow. Missing values were imputed using the half-minimum method; measurements were log2-transformed when the deposited scale metadata supported doing so, and each sample was median-centered. Differential analysis was then carried out separately for each replicated pairwise group comparison within a matrix using Welch’s t-test with Benjamini-Hochberg correction, and metabolites with an FDR-adjusted q value below 0.05 were considered significant. Log2 fold change, calculated as the difference between group means, was reported separately (Supplementary Methods 7: Contrast enumeration and statistical testing).

For each comparison, we recorded the minimum arm size, defined as the smaller of the two group sample counts, along with the number of sample-level factors that differed between the groups. Only comparisons with at least two biological samples in each group were tested. Comparisons that differed in more than one factor were flagged as potentially confounded, allowing simpler biological contrasts to be distinguished from those involving multiple simultaneous changes. Results were first stored at the matrix level, since each matrix was treated as a separate analytical unit. They were then carried forward to the study level without collapsing multiple analysis studies into a single differential-analysis result (Supplementary Methods 7: Confound annotation).

### Platform architecture: harmonized data backend and semantic retrieval frontend

The harmonized corpus powers HARMONY, a web platform for searching and comparing public metabolomics studies from MW and ML (https://omicsinharmony.in/). Built with TypeScript/Next.js and a SQLite data store, the platform keeps each record tied to its source accession, original deposited text, ontology mapping, and mapping evidence. The source repository remains the reference point for every result. Search is designed to work in two ways. Exact matches on ontology terms, accessions, RefMet identifiers, and canonical names are always given priority. When no exact match is available, the platform turns to semantic search through two dual encoders. One is trained on controlled-vocabulary synonyms using PubMedBERT^49^ and BiomedBERT^49^ with SapBERT^50^-style metric learning, and the other is trained on free-text study descriptions using a MedGemma^47^-derived contextual encoder. Both produce vector representations that are stored in an HNSW^51^ (Hierarchical Navigable Small World) index for fast similarity search across the full corpus. Results from exact matching, semantic search, and lexical scoring are then combined with ontology-distance and metabolite-overlap evidence, and the final list is re-ranked before being shown to the user. This ranked output supports search, filtering, and “Cross-Study” Comparison. A separate MedGemma-based module can also generate short, evidence-grounded summaries of individual studies. Because this module generates text rather than retrieving records, its output is evaluated separately and is not used as a validated search signal. The complete architecture of the hybrid search, ranking, and comparison framework is shown in Figure S15.

## Software Availability

HARMONY is freely available at https://omicsinharmony.in. The web platform provides ontology-aware search across Metabolomics Workbench and MetaboLights. Users can search by species, disease, sample source, analytical technique, metabolite name, study accession, or broader study terms. Cross-study comparisons are also supported across both repositories by filtering harmonized metadata, shared metabolite identifiers, and differentially abundant metabolites. Results can be explored through universal search, metadata filters, metabolite search, study-comparison views, and the harmonization dashboard. HARMONY is a read-only discovery layer: it does not replace, host, or modify the original repository records, and each result links back to the source study accession. A detailed user manual and a UI walkthrough using actual case studies are available on the website (Harmony: User Guide 1.0&HARMONY Case Studies_1.0). Platform updates can be tracked through the “**What is new**” section on the main landing page. The current version (v1.7.0) is live and is maintained with periodic updates as the underlying repositories grow.

## Acknowledgements

SB acknowledges support from the Prime Minister’s Research Fellowship.

