## Supplementary Table 1 for "Ontology-guided harmonization enables unified discovery of public metabolomics studies within and across repositories"

### Supplementary Methods

#### Supplementary Methods 1: Species mapping to NCBI Taxonomy

After extraction, species strings were standardized for ontology lookup by converting text to lowercase, trimming leading and trailing spaces, collapsing repeated whitespace, and removing punctuation during comparison. The original deposited species string was retained unchanged in the final output to preserve repository provenance. Species mapping was restricted to NCBI Taxonomy (NCBITaxon). When a MetaboLights ISA-Tab record already provided a valid NCBITaxon accession, the accession was directly converted to the corresponding OBO-format IRI. All other raw species strings were queried against OLS4 with the search space restricted to NCBITaxon. Example, for MTBLS423, the pipeline does:
**[native ML deposited]** http://purl.bioontology.org/ontology/NCBITAXON/9606 → [**pipeline normalized**] http://purl.obolibrary.org/obo/NCBITaxon_9606.

Mappings were prioritized according to conservative evidence rules. Exact matches to NCBITaxon labels were accepted first, followed by exact matches to NCBITaxon synonyms. Restricted fallback matches were accepted only when the normalized query matched the returned label or synonym after text standardization, or when a multi-token taxon name shared the same binomial prefix as the returned NCBITaxon label. Common biological aliases, including human, mouse, rat, pig, and cow, were expanded to their corresponding scientific names before lookup. Open-ended fuzzy matches were not accepted unless they satisfied these taxonomic consistency checks.

Multi-species annotations were handled as multiple taxonomic entries rather than being collapsed into a single composite label. For the Metabolomics Workbench, composite species strings were split using common delimiters, including semicolons, commas, slashes, and the conjunction "and," followed by duplicate removal. For MetaboLights, multi-species records were reconstructed from row-wise organism labels when available; organism strings recovered from the MetaboLights organisms API were also split before mapping. The standardized species node retained element-wise correspondence between each deposited raw organism string, its mapped NCBITaxon IRI, canonical label, ontology source, and taxonomic rank. Thus, a study annotated with two organisms, such as Ganoderma lucidum and Ganoderma leucocontextum, retained two raw organism values, two NCBITaxon IRIs, and two canonical labels.

Canonical taxon labels were retrieved from the local OLS4-derived label cache. Taxonomic ranks were **not** inferred from the wording of species labels and were **not** manually assigned. Instead, rank metadata were obtained from NCBI Entrez EFetch taxonomy records and stored with both single-organism and multi-organism species annotations. Search aliases were added only after the primary NCBITaxon mapping was finalized. These aliases were extracted from NCBI taxonomy name classes, including scientific names, synonyms, equivalent names, and common names. Aliases that mapped to more than one current NCBITaxon IRI were excluded from the search layer to reduce false-positive retrieval. Importantly, the alias layer was used only to support search and query expansion; it did not modify the accepted NCBITaxon mapping for any study.

Manual curation was applied to remove false-positive and over-specific mappings when supported by NCBITaxon or OLS evidence. Non-biological strings, environmental matrices, analytical blanks, food matrices without a specific taxonomic node, and community-level descriptors were retained as deposited text but left unmapped. When a deposited species-level organism name had been incorrectly mapped to a strain-level NCBITaxon entry without strain evidence in the source metadata, the mapping was corrected to the appropriate species-level or genus-level NCBITaxon term rather than preserving the over-specific hit.

**Supplementary Methods 2: Sample source mapping principles**

**Normalization, composite splitting, and OLS4 mapping:** Raw source strings were preserved unchanged for provenance, while a separate lookup-normalized form was generated for matching. Normalization consisted of lowercasing the string, trimming leading and trailing whitespace, collapsing repeated whitespace, converting `&` to `and`, replacing underscores and common source delimiters (`/`, `|`, `;`, `,`, `+`) with spaces, removing remaining non-alphanumeric punctuation, and collapsing the result to a single space-delimited token string. Thus, for example, `Culture media, conditioned` became `culture media conditioned`, and `CD8-positive T lymphocyte` became `cd8 positive t lymphocyte`. Composite raw values were split before ontology mapping when the delimiter clearly separated independent source materials. The split output retained an aligned split_index and split_raw_value, allowing multi-source studies to carry multiple source mappings without collapsing them into a single primary term (Table S1-S2). Controlled ontology mapping rules are available in Table S3. Disease, chemical/analyte, assay, procedure, and broad omics ontologies (DOID, MONDO, CHEBI, BAO, HL7, OMIT) were excluded as primary mapping targets because they describe donor clinical state, measured analytes, or assay design rather than the physical, biological,l or environmental material collected — accepting them would conflate sample identity with disease context or analytical purpose. Gene Ontology (GO) was similarly excluded from the general target set, except for GO:0016020 (membrane), which was retained as a curated correction for subcellular membrane fraction preparations where the corresponding UBERON term was biologically over-specific.

**Candidate scoring and acceptance for the sample source node**

For each normalized split term, the pipeline retrieved OLS4 candidates and scored them using lexical evidence, ontology tier, semantic category, and specificity checks. Exact label matches in primary ontologies were preferred, followed by exact synonym matches, normalized label/synonym matches, and constrained token-compatible matches. Candidate scores were penalized if the ontology tier was fallback, if the candidate belonged to a rejected semantic type, or if the candidate added unsupported specificity relative to the raw term (Table S4). The final `mapping_status` was explicitly assigned to every row. Accepted statuses were `accepted_exact_label`, `accepted_exact_synonym`, `accepted_constrained_match`, and `accepted_with_caveat`.

Non-accepted statuses were `rejected_non_source`, `rejected_ambiguous`, `rejected_low_confidence`, `unmapped`, and `missing`. A null mapping was treated as a valid outcome when the raw value was not a defensible sample-source concept.

**Supplementary Methods 3: Analytical technique raw term extraction endpoints for both databases**

**MetaboLights**

MetaboLights records were queried from the public REST endpoint https://www.ebi.ac.uk/metabolights/ws/studies/{MTBLS_ID}. Assay metadata were parsed from isaInvestigation.studies[0].assays[], with one assay object treated as one analytical record. In some cases, isaInvestigation.studies[0].studyDesignDescriptors was used to extract analytical technique terms when they were not available directly.

**Metabolomics Workbench**

For Metabolomics Workbench (MW), analytical-technique labels were reconstructed from the per-analysis REST metadata rather than from study-level summary labels, MWTab-derived labels, metabolite-level fields, existing standardized ontology nodes, or previously assigned ontology values. MW study identifiers were used only to enumerate records for endpoint retrieval. Analytical technique evidence was then extracted from the MW analysis endpoint: https://www.metabolomicsworkbench.org/rest/study/study_id/{STUDY_ID}/analysis

Each returned analysis object was treated as one analytical record. When the endpoint returned a single analysis object, it was parsed as one record; when it returned multiple analysis objects, each new object was retained separately after sorting by analysis ID. The extraction used only fields present within each returned analysis record. These included analysis_id, analysis_summary, analysis_type, chromatography_type, chromatography system, column_name, ms_type, ion_mode, ms_instrument_name, ms_instrument_type, nmr_instrument_type, nmr_experiment_type, spectrometer_frequency, nmr_solvent, and units. These fields were used as complementary evidence sources because MW analysis records often distribute method information across broad analysis types, chromatography descriptors, detector or instrument descriptors, and MS- or NMR-specific fields. Study title, study description, summary.analysis_type, MWTab-derived labels, metabolite-level fields, existing standardized ontology assignments, and external web text were explicitly excluded from analytical-technique assignment.

Raw endpoint strings were preserved unchanged as provenance fields. Normalized strings were generated only for matching and rule evaluation. Normalization harmonized case, spacing, and punctuation variants so that equivalent notations such as LC-MS, LC/MS, and LC MS could be compared consistently. At the same time, the original deposited strings remained available in the row-level output.

**Assignment rules**: NMR was assigned when the analysis type or NMR-specific fields indicated an NMR experiment. Detector-coupled non-MS methods, including LC-FLD and GC-FID, were protected before broader MS rules. Source- or modality-specific MS techniques, including PSI-MS, DESI-MS, MALDI-MS, APCI-MS, and ICP-MS, were evaluated before generic LC/GC rules. Direct-infusion rows were assigned DI-MS when direct-infusion evidence was present, provided no stronger GC-specific evidence was available. FIA-MS, CE-MS, SFC-MS, GC-MS, and LC-MS were then assigned from corresponding chromatography, instrument, column, or method cues. API-MS was retained when atmospheric-pressure ionization was the only specific cue and no APCI/APPI-level assignment was defensible. Generic MS was retained only when analysis_type = MS and the record lacked sufficient chromatography, detector, direct infusion, ionization modality, instrument, or NMR evidence to support a more specific technique. Rows without usable endpoint evidence were marked unresolved rather than inferred from excluded fields. At the study level, unique accepted techniques were aggregated across all analysis records, while retaining all distinct techniques rather than selecting a single primary platform.

**Ontology mapping**: After endpoint-only extraction, normalized technique labels were mapped to ontology IRIs. CHMO was used as the primary analytical-method ontology because it provides method-level terms for LC-MS, GC-MS, NMR, CE-MS, MALDI-MS, APCI-MS, ICP-MS, detector-coupled chromatographic methods, and related analytical techniques. Secondary ontology terms were used only when CHMO lacked a suitable method-level term. DI-MS was retained using a TransformON/TFO term with an explicit caveat because a clean CHMO assay-level DI-MS term was not available. PSI-MS was retained using the HUPO-PSI MS controlled vocabulary because the resolved term represented a source-specific MS modality. Terms mapped to component methods rather than complete coupled techniques, such as FIA-MS and SFC-MS, were retained with caveats rather than treated as exact full-method equivalents.

**NMR experiment-type refinement**

NMR experiment-type mapping was treated as a refinement of the analytical-technique node and was applied only to records whose broad analytical technique indicated nuclear magnetic resonance spectroscopy. It was not used as an ionization source, for ion polarity, for mass analyzer type, or for separation method. The assignment unit followed the analytical-technique record-level structure: either a single MW/analysis record or a single ML assay record, with study-level summaries generated solely by aggregating record-level NMR terms.

For MW, NMR evidence was extracted from direct /analysis fields, including analysis_type, analysis_summary, nmr_experiment_type, nmr_instrument_type, spectrometer_frequency, nmr_solvent, and units. NMR records were identified when these fields indicated NMR (nuclear magnetic resonance) or a recognizable NMR experiment family, such as NOESY, CPMG, HSQC, HMBC, COSY, TOCSY, J-resolved/JRES, HRMAS/MAS, or MR imaging.

For ML, NMR evidence was extracted from assay-level ISA metadata and cached assay-file metadata, including assay filename, assay type label, technology type, technology platform, Parameter Value[Pulse sequence name], Parameter Value[Instrument], Parameter Value[Magnetic field strength], Parameter Value[NMR Probe], Parameter Value[Solvent], Parameter Value[NMR tube type], Parameter Value[Number of transients], Parameter Value[Temperature], and relevant protocol names or support text. The assay-file pulse-sequence field was treated as the strongest ML evidence for experiment-family assignment. Shared protocol descriptions were retained as support text but were not used to assign every NMR method mentioned in a protocol block to every assay.

All NMR acquisition records were assigned the broad CHMO parent term nuclear magnetic resonance spectroscopy (CHMO:0000591). More specific CHMO terms were added only when direct record-level evidence reported the nucleus, dimensionality, pulse sequence, or experiment family. Vendor pulse-program names were retained as raw metadata and mapped only to a parent experiment family when defensible; for example, a NOESY pulse program was mapped to the NOESY family rather than treated as a separate vendor-specific ontology term. Probe nuclei were not used alone to infer the NMR nucleus, so a probe string such as 1H-13C/15N/D remained probe metadata unless the experiment field itself supported 1H or 13C NMR.

NMR-related acquisition details that were useful for reuse but not suitable as experiment-type ontology terms were retained as raw or literal metadata. These included FT-NMR evidence, spectrometer frequency or magnetic field strength, instrument model, probe, solvent, tube type, temperature, number of transients, vendor pulse-program strings, and shared protocol descriptions.

**Supplementary Methods 4: Ionization source and ion polarity extraction and mapping for both databases**

**Ionization source:** The ionization source was reconstructed independently from repository-level evidence. The node was restricted to MS ionization sources or methods, such as electrospray ionization, electron ionization, APCI, APPI, MALDI, DESI, paper spray ionization, proton-transfer-reaction ionization, and related source concepts. It was not allowed to absorb metadata for ion polarity, analytical technique, mass analyzer, separation method, detector type, or instrument model.

For Metabolomics Workbench, the extraction unit was one row returned by the /rest/study/study_id/{STUDY_ID}/analysis endpoint. The native ms_type field was treated as the primary ionization-source field. The analysis_type field was used as context to distinguish MS from non-MS analyses. When ms_type was blank, generic, other, or multiple, only explicit ionization-source strings in analysis_summary, ms_instrument_name, or ms_instrument_type were used as fallback evidence. chromatography_type was retained as a context variable but was not used to infer the ionization source. Similarly, ion_mode was retained only as polarity provenance and was not mapped as an ionization source.

For MetaboLights, the full study metadata were first used to identify assay records and their corresponding assay metadata filenames. Each assay metadata file was then parsed, and Parameter Value[Ion source] was treated as the primary ionization-source field. Adjacent Term Source REF and Term Accession Number columns were aligned by column position to retain any depositor-provided controlled-vocabulary evidence. The technology platform was used only as fallback evidence when it explicitly contained an ionization-source term. Assay type labels were used only when they named a source-bearing method, such as MALDI-MS, DESI-MS, APCI-MS, PTR-MS, SESI-MS, or paper spray MS. Broad technology type and scan polarity were not used as ionization-source evidence.

Across both repositories, mappings were deliberately conservative. ESI was not inferred from LC-MS, EI was not inferred from GC-MS, and source terms were not inferred from polarity, chromatography, analyzer type, or instrument family alone. Non-MS and detector-only records, including NMR, LC-DAD, LC-FLD, GC-FID, UV, fluorescence, and related detector assays, were marked as not applicable for this node. Placeholder, missing, unspecified, analyzer-only, polarity-only, or acquisition-mode-only values were retained as null or unresolved.

Accepted terms were mapped primarily to PSI-MS because it provides controlled source-level terms for MS ionization methods. CHMO or other secondary vocabularies were used only when the deposited value was method-level, and no suitable PSI-MS source-level term was available. For each mapping, the final table retained the raw value, normalized value, source field, evidence text, mapped IRI, mapped label, ontology prefix, mapping status, confidence category, caveat reason where applicable, and hierarchy/search aliases. Exact IRIs were used for harmonized matching, while parent and hierarchy terms were used only for broader search and visualization.

Rows with no deposited ion-source evidence were retained as null. Broad API-class MW rows were retained only as atmospheric-pressure-ionization-class evidence and were not over-resolved to ESI, APCI, or APPI. DuoSpray and Agilent Jet Stream cases were mapped to electrospray ionization only when supporting evidence justified this assignment. Detector-only GC-FID and fluorescence rows were marked not applicable. One GCxGC-TOF case with inconsistent API metadata was corrected to electron ionization based on the instrument and analytical context. SIMS primary-beam descriptors were not forced into the standard ionization-source vocabulary and were retained as policy-sensitive unresolved cases pending a dedicated SIMS ontology policy.

**Ion polarity:** For Metabolomics Workbench, the extraction unit was one analysis row from the official study analysis endpoint, and polarity was assigned only from raw_endpoints.analysis.ion_mode. For MetaboLights, the extraction unit was a single assay file; study-level REST metadata were used to enumerate assays, and row-level assay-file metadata were then parsed to recover values for Parameter Value[Scan polarity]. Assay filenames, technology platform strings, analytical technique labels, ionization-source labels, chromatography fields, instrument metadata, and study-design descriptors were retained as provenance or applicability context. Still, they were not used to infer polarity.

Ion polarity was mapped to the PSI-MS acquisition-polarity branch because the mapped entity represents an analysis or assay acquisition record rather than an individual scan. Positive-only records were mapped to positive polarity acquisition (MS:1000077), negative-only records to negative polarity acquisition (MS:1000076), direct positive-and-negative records to mixed polarity acquisition (MS:1003774), and explicit alternating or switching records to alternating polarity acquisition (MS:1002833). For compatibility with scan-level representations, positive and negative acquisition records could also be linked to the older PSI-MS terms positive scan (MS:1000130) and negative scan (MS:1000129). Still, the acquisition polarity terms were used as the final study/assay-level nodes. Values that described scan mode, resolution, mass range, or missing information were deliberately left unmapped. NMR, MR imaging, LC-DAD, GC-FID, UV, fluorescence, and other non-MS or detector-only assays were treated as outside the scope of the ion-polarity node and were not mapped to polarity IRIs. At the study level, ion polarity was stored as the deduplicated union of all mapped acquisition-polarity concepts observed across the study's analysis or assay records. Record-level provenance was retained so that positive-plus-negative study profiles could be traced either to separate acquisition records or to a single mixed/alternating record. Ionization-source and ion-polarity combinations were generated only after both nodes had been independently mapped from their respective direct evidence fields; ionization-source terms were not used to infer polarity.

#### Supplementary Methods 5: Separation method and mass analyzer term harmonization and mapping

Raw separation strings were preserved in the row-level extraction table and were not overwritten by normalized labels. Normalization was used only for classification and ontology mapping, so that repository-specific abbreviations, column chemistries, platform shorthand, and assay-file descriptors could be collapsed into a controlled set of canonical separation-method categories.

Reversed-phase LC evidence included direct terms such as "RP" and "reverse phase". It reversed-phase, as well as column-chemistry terms commonly used to indicate reversed-phase chromatography, including C18, C8, PFP, ODS, and HSS T3. HILIC evidence included direct HILIC strings and related column descriptors such as amide and ZIC-pHILIC. Broad LC evidence, such as LC, LC-MS, or liquid chromatography, was retained as liquid_chromatography only when a more specific LC mode could not be resolved. Similarly, UPLC/UHPLC and HPLC were retained as implementation-level LC descriptors and were not treated as equivalent to mode-specific terms such as reversed-phase LC or HILIC.

GC evidence included direct GC or GC-MS labels and GC column models such as DB-5, HP-5, VF-5, and RTX-5. Explicit multidimensional GC evidence, including GCxGC, GC x GC, 2D-GC, and Pegasus 4D, was mapped separately to two-dimensional GC. Capillary electrophoresis evidence included CE, CE-MS, and explicit capillary-electrophoresis strings.

Flow-injection analysis and direct infusion were treated as distinct, non-chromatographic-separation acquisition contexts. FIA, FIAMS, and flow injection were mapped to flow_injection_analysis, whereas DI, DIMS, direct infusion, shotgun lipidomics, NanoMate, and TriVersa NanoMate were mapped to direct_infusion_no_separation. Direct infusion was retained as a real sample-introduction mode rather than treated as missing separation metadata, because these records describe sample delivery directly to the ion source without chromatographic or electrophoretic separation.

Explicit no-chromatography or no-separation MS contexts, including MALDI, DESI, MS imaging, SESI, paper spray, ICP-MS, PTR-MS, and related source-driven acquisitions, were mapped to the internal reporting class no_chromatographic_separation only when no chromatographic or electrophoretic separation evidence was present. This class was retained without an ontology IRI because it represents the absence of chromatographic separation rather than a positive separation method.

#### Supplementary Methods 6: disease-node extraction and ontology mapping

The disease node was implemented as a study-level metadata node. A study could retain multiple disease or disease-context terms when the deposited metadata supported multiple conditions, infections, disease models, or clinical endpoints. The final standardized JSON representation separated strict disease ontology mappings from broad fallback mappings, valid unmapped disease-like literals, excluded non-disease context terms, and non-extractable studies. This layered design was used to avoid overrepresenting weak disease evidence while preserving relevant context for search and audit.

##### Metabolomics Workbench disease extraction

Metabolomics Workbench disease terms were extracted from the official REST disease endpoint: **https://www.metabolomicsworkbench.org/rest/study/study_id/{study_id}/disease.** For each study, the disease endpoint was queried, the returned term was cached, and disease strings were extracted directly from the response. Multiple disease entries were retained, and pipe- or semicolon-separated strings were split into separate term objects. Empty endpoint responses were treated as **missing_from_mw_rest_endpoint**, not as failed ontology mapping. MW raw disease values were then mapped through OLS4. Exact labels and exact synonyms were preferred. Depositor variants and abbreviations were normalized only when the intended concept was clear, such as "Diabetes" to "diabetes mellitus", "NASH" to "nonalcoholic steatohepatitis", "ALS" to "amyotrophic lateral sclerosis", and spelling variants such as "Alzheimers disease" to "Alzheimer's disease". The accepted ontology was determined from the returned IRI namespace, because OLS results can include imported terms. Disease-centric ontology mappings were favored, but secondary ontology mappings were retained when they represented the submitted MW disease field more closely than a forced disease term.

##### MetaboLights disease extraction

MetaboLights does not provide a structured disease endpoint like MW, so disease terms were extracted from study-level and ISA-derived metadata instead. We used the study ID, title, summary, design descriptors, and their accessions, Study Factor Name, Study Factor Type, and Factor Value fields to determine whether a disease or condition was actually present. Study descriptions and design descriptors provided the main free-text evidence. At the same time, factor fields were used to check whether a candidate term reflected a real study condition, model, endpoint, or disease context rather than background text. Raw strings were retained for provenance, and normalized strings were used only for splitting, comparison, and ontology matching.

Disease candidates were generated independently by three extractors: a token-level NER model based on BioBERT, fine-tuned on the NCBI Disease corpus; and two instruction-tuned biomedical language models, MedGemma-27B and GPT-OSS-20B. The models were prompted to extract only disease terms directly supported by the text, preserve literal forms and abbreviations, and return “None” when no disease was present. Positive, negative, and explicit no-disease examples were included to prevent the models from inferring disease from pathway names, treatments, organisms, environmental stress, diet, exposure, or model-system language. Decoding was deterministic for reproducibility. Pipe-separated outputs were split by study before reconciliation so that each candidate could be reviewed separately.

The three outputs were then reconciled with a two-of-three consensus rule applied to cleaned, normalized strings. Single-extractor terms were kept as lower-confidence candidates rather than discarded. The reconciled set, including both consensus terms and single-extractor candidates, was then reviewed manually against the same evidence fields used during extraction. Single-extractor terms were promoted only when the metadata clearly supported them. Terms that reflected background context, organism-only labels without disease evidence, non-disease study types, and other residual terms were excluded. Multiple disease labels were retained when the study design supported more than one concept, and valid disease-like terms without a safe ontology match were kept as unmapped literals rather than forced into an incorrect assignment.

**Ontology mapping and layered disease representation**

Accepted disease terms were mapped using OLS4, with disease-centric vocabularies as the primary target sets: MONDO, MeSH, DOID, NCIT, and EFO. Exact label and exact-synonym matches were preferred. Fallback matches were accepted only when the mapped term faithfully represented the deposited disease concept. ML final mapped disease terms were restricted to the primary disease ontology set. In contrast, MW allowed a small number of secondary ontology context terms when the submitted endpoint value itself was not a classical disease term but was faithfully represented elsewhere.

The final disease-node schema used layered mapping statuses: “strict_accepted_mapped” or “mapped_strict_ontology” represented the main disease ontology layer. “broad_fallback_mapped” represented broad but evidence-supported fallback handles, such as cancer or infection, when no specific supported disease was available. “valid_unmapped_literal” represented disease-like literals were retained without accepted IRIs. “excluded_context_or_fragment” represented fragments, organism-only labels, method/background terms, and false-positive leakage. Non-extractable was used for ML studies in which no disease or condition was supported after the evidence review. “missing_from_mw_rest_endpoint” was used when the MW REST disease endpoint returned no disease value.

##### Sanity checking and finalizing the disease node

After OLS4 mapping, disease mappings were reviewed for semantic consistency between the raw string and the mapped ontology label. Broad labels were checked to ensure that their generality was intentional rather than a failed attempt to map a more specific term. Unmapped literals were reviewed to determine whether they lacked a safe ontology match or whether a correct term could be recovered through manual hierarchy browsing. Unsafe mappings, including organism-only labels, gene or protein-like matches, chemicals, anatomy terms, or non-disease process terms, were excluded from the strict disease layer.

The final standardized JSON disease blocks were regenerated for both repositories. For both MW and ML, the disease-node values included raw values, mapped values, strict/secondary/unmapped layers, mapping status, aliases, synonyms, parent labels, hierarchy terms, and OLS URLs. For ML, additional fields included are extractability labels, strictly accepted mapped terms, broad fallback terms, valid unmapped literals, excluded context terms, non-extractable categories, raw literals, ontology metadata, synonyms, parent labels, hierarchy labels, OLS URLs, and UI filtering booleans.

##### Supplementary Methods 7: Differential abundance analysis of metabolites

**Non-biological sample removal:** Sample-level removal of quality-control, pooled-reference, blank, solvent, internal-standard, and calibration-standard rows was performed using a rule-based classifier operating jointly on sample identifiers, deposited factor strings, and sample-type fields, applied identically to both repositories before any quality-tier screening or factor-based grouping. Narrow analytical-control abbreviations occurring within otherwise biological factor strings (for example, an MW factor value indicating a field blank) were also expanded and excluded at this stage. In contrast, generic biological labels such as an experimental control arm were retained as valid biological samples rather than treated as analytical controls.

**Experimental factor parsing and joint-group construction:** For ML, factor columns were identified as all sample-sheet headers matching "Factor Value[...]" (case-insensitive), normalized by stripping the wrapper syntax, lowercasing, and replacing spaces with underscores. The factor table was joined to the matrix columns using the sample-sheet sample identifier column, and the resulting dataset was restricted to samples present in the current MAF. For MW, factor values were retrieved from the study-wide "factors.json" record (sourced from the REST factors endpoint), parsed from pipe-separated key-value strings, and joined to matrix columns using the local sample identifier rather than the datatable "Class" column, which was found to be incomplete relative to the "factors" endpoint in a subset of studies. The sample-source field, when present, was included as an additional factor dimension. In addition to name- and value-based exclusion of technical and subject-identifier factors (below), all literal missing-value tokens present in deposited factor fields (NA, N/A, -, unknown, nan, ND, "not available", and equivalent case-insensitive variants) were normalized to an empty value before joint-group construction in both repositories, so that samples sharing a missing-value token in a given factor were not incorrectly pooled into a spurious shared group.

Factor names matching a fixed technical or administrative set (replicate, batch, analysis time, run day, column number, and composite variants) were excluded from candidate grouping variables in both repositories; this filter was initially ML-only and was extended to MW after auditing showed MW could otherwise select a purely technical field as its sole grouping variable. A complementary value-level check excluded factors whose values matched per-subject identifier patterns (e.g., sequential patient or donor codes), regardless of field name, because subject identity was found in biologically named fields in some studies. Another filter was applied to factors with very high value cardinality and low per-group replication. Among factors passing both filters, those carrying recognized biological or experimental semantics were ranked high. The full joint combination of retained factors defined groups wherever it yielded at least two replicated groups, falling back to a single primary factor otherwise. Samples with no usable factor values were excluded from the contrast definition. The complete deposited grouping, including unreplicated groups, was retained in matrix-level output for audit.

**Quality-tier classification and disposition:** The quality-tier scheme distinguished three structurally different kinds of evidence, kept separate by design. Tier 1 comprised hard, non-corroborating screens: rejection for any negative abundance value or a near-constant feature distribution — either of which excluded a matrix outright regardless of any other evidence. Tier 2 comprised declared-metadata classification, parsing depositor-reported units (MW) or data-transformation protocol text (ML) against a fixed keyword taxonomy: raw intensity, concentration, vendor-normalized (e.g., probabilistic quotient or total-ion-current normalized), log-transformed, ratio or fold-change, already-scaled, and square-root-scale. Tier 3 comprised a distributional heuristic (a z-score-like signature check on the observed value distribution) used exclusively to corroborate or override the Tier 2 classification; Tier 3 evidence was never sufficient, on its own, to admit a matrix into the eligible pool in the absence of supporting Tier 2 metadata.

A ratio/fold-change classification (Tier 2) that lacked corroboration from Tier 1 (no negative values) or a z-score signature (Tier 3) was not excluded by default. Because this classification was found, on manual review, to arise predominantly from protocol-text language describing downstream statistical analysis rather than the actual deposited abundance scale, an uncorroborated ratio/fold-change label was instead demoted to the next most defensible class present as co-occurring evidence for that matrix, following a fixed conservative-first priority order (already-scaled, already-square-root, already-log, vendor-normalized, raw intensity, concentration) applied identically to Metabolomics Workbench and MetaboLights matrices. This demotion rescued a substantial subset of matrices that would otherwise have been lost in an ambiguous, excluded state despite carrying perfectly usable scale evidence elsewhere in the same metadata record. Square-root-scale deposits, whether classified directly from Tier 2 evidence or reached via this demotion path, were excluded outright in this version of the pipeline; no inverse square-root or log-of-square-root transformation was attempted to rescue these matrices, given their low overall prevalence in the corpus and the added risk of introducing an undocumented, non-standard transformation step. Matrices lacking any Tier 2 classification, or with an irreconcilable conflict between Tier 2 and Tier 3 evidence, were excluded as ambiguous rather than routed to manual curation.

Briefly, these were the overall nine conditions for DE-eligibility

1. **(Phase 1) Structural validity** — matrix parses; ≥1 metabolite row; ≥2 sample columns; numeric coercion succeeds; ≥1 strictly positive value
2. **(Phase 2) Non-empty after QC** — ≥2 biological samples remain after non-biological (QC/blank/pool/standard) sample removal
3. **(Phase 3) Valid factor grouping** — full-joint or primary-factor fallback yields ≥2 replicated groups (n≥2/arm); technical/administrative and subject-identifier factors excluded from candidates
4. **(Phase 4) No hard numeric red flags** — no negative abundance values; not near-constant/degenerate; not all-nonpositive
5. **(Phase 5) Declared scale present (Tier 2)** — units/protocol text classifies to a known scale (raw intensity, concentration, vendor-normalized, already-log); not absent/unclassifiable
6. **(Phase 6) Distributional corroboration (Tier 3)** — not a z-score-like signature (hard exclude); distribution doesn't contradict the declared scale
7. **(Phase 6) Not a hard-excluded scale class** — not already-scaled; not already-sqrt (both excluded outright in v1)
8. **(Phase 6) Ratio/fold-change resolved, not uncorroborated** — uncorroborated ratio_or_fc demoted to a supported class via fixed priority order, not left ambiguous
9. **(Phase 7) Final disposition ∈ eligible set** — "pass_full_pipeline", "pass_log_skipped", or "pass_vendor_norm_flagged" (everything else — exclude_invalid, exclude_ambiguous, exclude_pretransformed, exclude_degenerate, exclude_nonpositive -> is not DE-eligible)

**Conditional preprocessing**. Matrices classified as raw intensity or concentration were imputed for missing and zero values using half the minimum positive value observed per metabolite, log2-transformed, and sample-wise median-centered on the log2 scale. Matrices that were already log-transformed were median-centered on the deposited scale, without a second logarithmic transformation. Matrices classified as vendor-normalized were carried through the identical imputation, log2-transformation, and median-centering procedure as raw-intensity matrices, but retained an explicit annotation flagging the pre-existing depositor-applied normalization for downstream interpretation.

**Contrast enumeration and statistical testing.** For each eligible matrix, we tested all unordered pairwise comparisons between replicated groups after removing non-biological samples, requiring at least two biological samples per group. To keep the analysis tractable, we set a maximum of 5,000 replicated pairwise comparisons per matrix. Matrices exceeding this limit, typically due to highly fragmented designs or very large profiling schemes, were not tested; instead, they were recorded as skipped with an explicit machine-readable reason, so that every excluded matrix remained visible in the audit trail. Within each tested comparison, metabolites missing in more than 50% of samples in either group were omitted from that comparison.

Differential abundance for the remaining metabolites was assessed with Welch's two-sample t-test, which does not assume equal variances. We chose this test rather than the Student's t-test, as used in tools such as MetENP, because the data span many independent studies, instruments, and sample types, in which within-group variance is unlikely to be uniform. Raw p-values were adjusted within each contrast and matrix using the Benjamini-Hochberg procedure; adjustment was not pooled across contrasts, matrices, or studies. Metabolites with an adjusted FDR q-value below 0.05 were considered significant, and the nominal p-value threshold of 0.05 was also retained for descriptive reference. Log2 fold change was defined as the difference between group means on the scale used for testing. Statistical testing and multiple-testing correction were implemented in Python 3 with SciPy.

**Confound annotation.** Each pairwise comparison was annotated with the number of joint-group factors that differed between the two groups, along with a binary flag indicating whether the comparison was confounded when two or more factors changed simultaneously. We also recorded the minimum arm size for each comparison and whether either group met the minimum replicate threshold of $n$=2. At the matrix level, we summarized the numbers of clean and confounded comparisons separately, along with the numbers of FDR-significant metabolites in each category.

**Example of confounding factors from a given study:**
**Study/Analysis:** ST000403 / AN000643
**The full factor set in that matrix is:**

- cell_type
- glucose_labelling
- sample_source
- timepoint

**Now, if we compare the following cases:**

**Not confounded**
Group 1: cell_type:Mature erythrocyte | glucose_labelling:C13 glucose | sample_source:Cells | timepoint:1 hour
Group 2: cell_type:Mature erythrocyte | glucose_labelling:C13 glucose | sample_source:Cells | timepoint:20 hours
**Only timepoint changes: n_factors_differing = 1**
**is_confounded = false**

**Confounded**
Group 1: cell_type:Mature erythrocyte | glucose_labelling:C13 glucose | sample_source:Cells | timepoint:1 hour
Group 2: cell_type:Mature erythrocyte | glucose_labelling:unlabelled | sample_source:Cells | timepoint:20 hours
**glucose_labelling and timepoint both change**
**n_factors_differing = 2**
**is_confounded = true**

**More strongly confounded**

Group 1: cell_type:Mature erythrocyte | glucose_labelling:C13 glucose | sample_source:Cells | timepoint:1 hour
Group 2: cell_type:Reticulocyte | glucose_labelling:unlabelled | sample_source:Cells | timepoint:20 hours
**cell_type, glucose_labelling, and timepoint all change**
**n_factors_differing = 3**
**is_confounded = true**

**So the rule is:**

- 1 differing factor => clean comparison
- 2 or more differing factors => confounded comparison

sample_source stays constant in these examples, so it does not contribute to confounding.

### Supplementary Tables

**Table S1: Normalization and splitting rules for composite terms for the sample source**

| Rule | Handling |
|---|---|
| Semicolon, pipe, plus | Split when they separated the source terms |
| Slash | Split only when both sides were clear biological entities, not abbreviations |
| Comma | Split only when it looked like an enumerated list of independent short source phrases |
| and | Split only when it connects short independent source phrases |
| Protected phrases | Not split when the phrase represented a single source concept, such as large intestine, breast milk, cell culture medium, conditioned medium, or whole body minus foot. |

**Table S2: Example compound phrases split into multiple raw terms before ontology mapping**

| Example phrase | Study ID | DB | Raw composite | Split terms |
|---|---|---|---|---|
| Plasma, Liver -> Plasma and Liver | ST000487 | MW | Plasma, Liver | Plasma; Liver |
| leaf;root -> leaf and root | MTBLS4286 | ML | leaf;root | leaf; root |
| HeLa cell;NIH-3T3 cell -> HeLa cell and NIH-3T3 cell | MTBLS78 | ML | HeLa cell;NIH-3T3 cell | HeLa cell; NIH-3T3 cell |
| thorax;abdomen -> thorax and abdomen | MTBLS180 | ML | thorax;abdomen | thorax; abdomen |
| Cecal contents and Ileal contents -> Cecal contents and Ileal contents | MTBLS12825 | ML | Cecal contents and Ileal contents | Cecal contents; Ileal contents |

**Table S3: Controlled OLS4 ontology mapping rules for sample source**

| Ontology tier | Ontologies | Role in sample-source mapping |
|---|---|---|
| Primary | UBERON, BTO, CL, CLO, Cellosaurus, NCBITaxon, ENVO, FOODON, PO | Preferred targets for anatomy, tissues, biofluids, cell types, named cell lines, whole organisms, environmental materials, food/material matrices, and plant anatomical structures, respectively |
| Secondary | OBI, EFO, SNOMED CT | Accepted only after semantic validation when the term represented a specimen, body substance, anatomical structure, cell/cell line, organism, culture material, or biological/material source |
| Fallback/high-risk | MESH, NCIT | Accepted only after explicit review when no better primary or secondary mapping existed, and the concept was semantically a source material |
| Rejected as primary | DOID, MONDO, CHEBI, BAO, HL7, OMIT | Disease, chemical/analyte, assay, procedure, administrative, or broad omics concepts were not accepted as primary sample-source mappings |
| Curated exception | GO | Not part of the general target set. Used only for the final membrane correction to GO:0016020, because the previous UBERON membrane organ mapping was biologically over-specific for a subcellular membrane fraction |

**Table S4: Candidate scoring for source node**

| Evidence type | Interpretation | Examples |
|---|---|---|
| exact_label | Normalized raw value matched the candidate label | Blood → blood / UBERON_0000178; Intestine → intestine / UBERON_0000160 |
| exact_synonym | Normalized raw value matched a candidate synonym | Stool → feces / UBERON_0001988; Cerebrum → telencephalon / UBERON_0001893 |
| normalized_label or normalized_synonym | Match became exact after safe punctuation or hyphen normalization | B-cells → B cell / CL_0000236; HCT-116 cell → HCT 116 cell / CLO_0003665 |
| constrained_token_match | Raw and candidate shared the meaningful head term, but the match required review for specificity | Fibroblast cells → fibroblast / CL_0000057; Maize starchy endosperm → endosperm / PO_0009089 |
| source_iri_verified | Supported by a deposited source IRI if enabled; for this freeze, native source IRIs were kept as provenance and not used automatically | Example provenance case: ML urine had source/candidate BTO_0001419, but final primary mapping was revised to UBERON_0001088 |
| no_valid_candidate | No candidate passed semantic validation | retention index → rejected as assay/procedure; blank → rejected as analytical blank/QC; Worms → retained unmapped/ambiguous |

**Table S5: Assignment rules for candidate ontologies to their respective source types**

| Final category | Assignment logic |
|---|---|
| anatomical_source | UBERON/BTO anatomical structures, organs, tissues, or reviewed anatomical fallback terms |
| biofluid | UBERON/BTO/SNOMED/NCIT/MESH body fluids or lavage/body-substance terms when semantically valid |
| cell_type | CL terms and reviewed cell-type terms |
| cell_line | CLO, Cellosaurus, or reviewed named cell-line/cultured-cell mappings |
| organism | NCBITaxon organism-level or whole-organism descriptors |

| Final category | Assignment logic |
|---|---|
| environmental_material | ENVO environmental materials such as soil, seawater, sediment, and wastewater |
| food_or_material_matrix | FOODON terms and reviewed beverage/food/material matrices |
| plant_structure | PO plant anatomical structures such as leaf, root, flower, seed, shoot, and fruit |
| culture_or_growth_medium | OBI/BTO/NCIT/MESH/EFO culture media, growth media, and culture-derived supernatants when source-like |
| specimen_or_sample_processing_descriptor | Cell pellets, extracellular vesicles, organoids, lysate-like material, sorted cells, xenografts, microsomes, mitochondria, and other source-bearing specimen/material descriptors |
| analytical_blank_or_qc | Blank, QC, pooled-QC, standard, and reference-control terms that were not biological source materials |
| disease_context | Disease-only strings or disease ontology concepts rejected as primary sample-source mappings |
| analyte_or_chemical | CHEBI/chemical/analyte terms or strings such as RNA, DNA, protein, bile acid, phospholipid, solvent, or reference compound |
| assay_or_procedure | Procedure, assay, measurement, device, administrative, or clinical information concepts |
| ambiguous | Under-specified source strings or concepts where a precise biological source could not be defensibly asserted |
| missing | Blank, placeholder, or not-applicable source strings |

**Table S6: Species-mapping audit and taxonomic coverage summary**

| Section | Metric or audit item | MW | ML | Overall / details | Interpretation |
|---|---|---|---|---|---|
| Repository coverage | Total studies included in species node | 4,118 | 2,712 | — | Initial study volume for taxonomic normalization. |
| Mapping success | Studies with at least one mapped NCBITaxon IRI | 4,058 | 2,586 | 98.5% (MW) / 95.4% (ML) | High taxonomic coverage was maintained across both metabolomics repositories. |

| Section | Metric or audit item | MW | ML | Overall / details | Interpretation |
|---|---|---|---|---|---|
| Unmapped curation | Raw species strings present but unmapped | 50 | 49 | — | Unmapped records were manually reviewed to assess biological validity. |
| Unmapped curation | Absent or non-usable deposited species metadata | 10 | 77 | — | Records with missing metadata were retained as explicit null species entries. |
| Unmapped curation | Representative null species entries | — | — | blank_sample, seawater, food mixtures | Mapping was avoided to prevent unsupported inference for non-organismal strings. |
| Taxonomic diversity | Unique mapped NCBITaxon IRIs | 419 | 1,267 | — | MetaboLights exhibited a significantly broader taxonomic vocabulary than Workbench. |
| Taxonomic diversity | Study coverage by the top five taxa | 82.4% | 58.9% | — | Workbench studies were highly concentrated around common model organisms. |
| Distribution structure | Singleton taxon frequency | — | 76.2% | 965 singleton IRIs in ML | MetaboLights demonstrated a pronounced long-tail taxonomic distribution. |
| Vocabulary depth | Taxa required for 90% cumulative study coverage (90% tier) | 25 | 127 | — | ML required a larger set of distinct taxa to cover its high-frequency space. |
| Vocabulary evidence | Exact OLS label matches (90% tier) | 25 | 101 | — | Exact labels exclusively supported Workbench high-frequency mappings. |
| Vocabulary evidence | Synonym-level mappings (90% tier) | 0 | 5 | — | Synonym rescue accounted for only a small fraction of high-frequency ML IRIs. |

| Section | Metric or audit item | MW | ML | Overall / details | Interpretation |
|---|---|---|---|---|---|
| Vocabulary evidence | Constrained token-compatible matches (90% tier) | 0 | 9 | — | Limited fallback matching was necessary for select high-frequency terms. |
| Vocabulary evidence | Reviewed or caveated mappings (90% tier) | 0 | 12 | Typos and strain descriptors | Manual review resolved inconsistencies in major repository descriptors. |
| Complexity metrics | Studies with multiple mapped NCBITaxon IRIs | 85 | 257 | — | Multi-species records were more prevalent in the MetaboLights repository. |
| Taxonomic rank | Primary rank distribution | Species -level | Species -level / Varied | Strain, genus, and no-rank | Rank preservation allowed capturing varied taxonomic granularity without forcing. |
| Cross-DB overlap | Common NCBITaxon IRIs | 132 | 132 | Shared across repos | The overlap in taxonomic vocabulary across databases was modest. |
| Cross-DB overlap | Jaccard overlap coefficient | 8.49% | 8.49% | IRI level | Most taxonomically unique IRIs were specific to a single repository. |
| Cross-DB linkage | Studies linked to shared taxa | 3,854 | 2,063 | — | Commonly mapped taxa facilitated broad connectivity across both databases. |

**Table S7: Sample-source ontology harmonization results**

| Section | Metric or audit item | MW | ML | Overall / details | Interpretation |
|---|---|---|---|---|---|
| High-frequency vocabulary coverage | Sample-source IRIs required to reach 90% cumulative mapped-study frequency | 24 | 55 | — | ML required more distinct source IRIs to cover the high-frequency mapped-study space, supporting a broader long-tail source vocabulary. |
| High-frequency vocabulary evidence | Exact OLS label matches within the 90% frequency tier | 17 | 41 | — | Most high-frequency mappings were directly supported by exact OLS label evidence. |
| High-frequency vocabulary evidence | Constrained token-compatible matches within the 90% frequency tier | 2 | 5 | — | A small subset required controlled token-compatible matching. |
| High-frequency vocabulary evidence | Reviewed or caveated mappings within the 90% frequency tier | 5 | 9 | — | Reviewed mappings were retained separately from exact mappings. |
| High-frequency vocabulary evidence | Synonym-only mappings within the 90% frequency tier | 0 | 0 | — | The high-frequency vocabulary was not driven by synonym-only rescue. |
| Unique-term resolution evidence | Exact label mappings among unique accepted raw source terms | — | — | 797 / 955, 83.5% | Exact label matching dominated the unique-term mapping layer. |
| Unique-term resolution evidence | Manually curated mappings among unique accepted raw source terms | — | — | 94 / 955, 9.8% | Manual review was required for a minority of terms, mainly |

| Section | Metric or audit item | MW | ML | Overall / details | Interpretation |
| --- | --- | --- | --- | --- | --- |
| | | | | | curation-sensitive or edge-case mappings. |
| Unique-term resolution evidence | Constrained token-compatible mappings among unique accepted raw source terms | — | — | 64 / 955, 6.7% | Constrained matching was used as a controlled fallback rather than as the main mapping route. |
| Ontology tier support | Primary OBO ontology mappings among unique accepted terms | — | — | 572 / 955, 59.9% | Primary sample-source-relevant OBO ontologies supported the most accepted unique terms. |
| Ontology tier support | Secondary ontology mappings among unique accepted terms | — | — | 236 / 955, 24.7% | Secondary ontologies were retained after semantic review. |
| Ontology tier support | Reviewed fallback ontology mappings among unique accepted terms | — | — | 147 / 955, 15.4% | Fallback ontology mappings were retained only after review. |
| Row-level mapping decision | Accepted exact label | — | — | 6,410 / 7,951, 80.6% | Exactly accepted mappings dominated the row-level table. |
| Row-level mapping decision | Accepted exact synonym | — | — | 5 / 7,951, 0.1% | Synonym-only acceptance was rare. |
| Row-level mapping decision | Accepted constrained match | — | — | 122 / 7,951, 1.5% | Constrained matches were retained as a distinct evidence class. |
| Row-level mapping decision | Accepted with caveat | — | — | 581 / 7,951, 7.3% | Caveated mappings were kept queryable but were not treated as exact mappings. |

| Section | Metric or audit item | MW | ML | Overall / details | Interpretation |
|---|---|---|---|---|---|
| Row-level mapping decision | Rejected non-source | — | — | 189 / 7,951, 2.4% | Non-source deposits were explicitly retained as rejected mapping decisions. |
| Row-level mapping decision | Unmapped | — | — | 639 / 7,951, 8.0% | Unmapped rows were mostly ambiguous, underspecified, or unsupported by valid sample-source concepts. |
| Row-level mapping decision | Missing | — | — | 5 / 7,951, 0.1% | Missing rows were retained as explicit null decisions. |
| Residual rejected or null terms | Ambiguous or underspecified values | — | — | 641 rows | Largest residual class, reflecting deposits where a precise biological source could not be assigned. |
| Residual rejected or null terms | Specimen or processing descriptors | — | — | 87 rows | Retained as unresolved or rejected when not source-specific enough. |
| Residual rejected or null terms | Blank, QC, or control terms | — | — | 45 rows | Kept as non-biological source decisions rather than mapped to tissues or fluids. |
| Residual rejected or null terms | Analyte or chemical labels | — | — | 31 rows | Rejected as sample-source mappings. |
| Residual rejected or null terms | Disease-context strings | — | — | 23 rows | Disease-only terms were not treated as sample-source concepts. |

| Section | Metric or audit item | MW | ML | Overall / details | Interpretation |
|---|---|---|---|---|---|
| Residual rejected or null terms | Missing values | — | — | 5 rows | Retained as explicit missing values. |
| Residual rejected or null terms | Assay or procedure metadata | — | — | 1 row | Rejected as non-source metadata. |
| Representative curation decision | IRI-namespace validation | — | — | cerebellum in MTBLS1410 | CL-indexed brain atlas hit was remapped to UBERON anatomical structure. |
| Representative curation decision | False taxon removal | — | — | Adipose Pannus in ST003198 | NCBITaxon arthropod-genus hit was rejected, and only a caveated tissue-level source was retained. |
| Representative curation decision | Exposure or context rejection | — | — | Chow diet in ST003425 | FOODON dog-breed lexical hit was rejected because diet is not a biological sample source. |
| Representative curation decision | Depositor typo rescue | — | — | Extracellular vesicles in ST004002 | The spelling variant was corrected and mapped to extracellular vesicles, with a caveat. |
| Representative curation decision | Placeholder cleanup | — | — | Other in ST004443 | Placeholder value was set to missing or null because it was semantically vacuous. |
| Representative curation decision | Overspecificity control | — | — | membrane in MTBLS12808 | UBERON membrane-organ over-specificity was rejected, and caveated membrane material was retained. |

| Section | Metric or audit item | MW | ML | Overall / details | Interpretation |
|---|---|---|---|---|---|
| Representative curation decision | RNA substring bug fix | — | — | culture supernatant in MTBLS11049 | The supernatant was retained as a culture-derived source and was not rejected due to the internal RNA substring. |
| Representative curation decision | Blank or QC cleanup | — | — | blank sample in MTBLS10568 | Retained as a rejected non-source with no biological IRI assigned. |
| Representative curation decision | Assay or procedure rejection | — | — | retention index in MTBLS8356 | Retained as rejected non-source because it represents analytical metadata. |
| Representative curation decision | Disease-qualified material | — | — | Colorectal Cancer Cells in ST003029 | Mapped to malignant cell source with caveat and not to disease ontology. |
| Representative curation decision | Disease-only rejection | — | — | colorectal cancer in MTBLS13095 | Rejected as a disease-only term for sample-source mapping. |
| Representative curation decision | Reviewed fallback acceptance | — | — | Exhaled Breath Condensate in ST001437 | Accepted as a reviewed SNOMED specimen/body-substance concept. |

**Table S8: Distribution of both shared and unique mapped analytical technique terms across MW and ML (green shade -> shared terms)**

| Mapped label | Ontology | IRI | MW | ML | Shared |
|---|---|---|---|---|---|
| capillary electrophoresis-mass spectrometry | CHMO | CHMO_0000702 | yes | yes | yes |
| desorption electrospray ionisation mass spectrometry | CHMO | CHMO_0000484 | yes | yes | yes |
| direct injection mass spectrometry | TFO | c_bfzxXDMU | yes | yes | yes |
| flow-injection analysis | CHMO | CHMO_0002891 | yes | yes | yes |
| gas chromatography-flame ionisation detection | CHMO | CHMO_0001736 | yes | yes | yes |
| gas chromatography-mass spectrometry | CHMO | CHMO_0000497 | yes | yes | yes |
| liquid chromatography-mass spectrometry | CHMO | CHMO_0000524 | yes | yes | yes |
| matrix-assisted laser desorption-ionisation mass spectrometry | CHMO | CHMO_0000519 | yes | yes | yes |
| nuclear magnetic resonance spectroscopy | CHMO | CHMO_0000591 | yes | yes | yes |
| atmospheric pressure chemical ionisation mass spectrometry | CHMO | CHMO_0000473 | yes | | |
| inductively coupled plasma mass spectrometry | CHMO | CHMO_0000538 | yes | | |
| liquid chromatography-fluorescence detection | CHMO | CHMO_0001742 | yes | | |
| mass spectrometry | CHMO | CHMO_0000470 | yes | | |
| paper spray ionization | MS | MS_1003235 | yes | | |
| supercritical-fluid chromatography | CHMO | CHMO_0001003 | yes | | |
| 2D-GC-MS | TFO | c_ZS6Stfx2 | | yes | |
| imaging mass spectrometry | CHMO | CHMO_0000053 | | yes | |
| ion mobility spectrometry-mass spectrometry | CHMO | CHMO_0000499 | | yes | |
| isotope ratio mass spectrometry | CHMO | CHMO_0000506 | | yes | |
| liquid chromatography-photodiode array detection | CHMO | CHMO_0001738 | | yes | |

| Mapped label | Ontology | IRI | MW | ML | Shared |
|---|---|---|---|---|---|
| proton transfer reaction mass spectrometry | CHMO | CHMO_0000570 | | yes | |
| secondary electrospray ionisation mass spectrometry | CHMO | CHMO_0000490 | | yes | |
| ultra-performance liquid chromatography-mass spectrometry | CHMO | CHMO_0000715 | | yes | |

**Table S9: Distribution of both shared and unique mapped NMR terms across MW and ML**

| Canonical key | Mapped label | Ontology | Accession | Mapping note |
|---|---|---|---|---|
| nmr_spectroscopy | nuclear magnetic resonance spectroscopy | CHMO | CHMO:0000591 | Broad parent assigned to every NMR record |
| 1d_nmr | one-dimensional nuclear magnetic resonance spectroscopy | CHMO | CHMO:0000592 | Explicit 1D or pulse-sequence evidence |
| 1h_nmr | 1H nuclear magnetic resonance spectroscopy | CHMO | CHMO:0000593 | Experiment/type evidence only; not probe-only evidence |
| 13c_nmr | 13C nuclear magnetic resonance spectroscopy | CHMO | CHMO:0000595 | Experiment/type evidence only; not probe-only evidence |
| 2d_nmr | two-dimensional nuclear magnetic resonance spectroscopy | CHMO | CHMO:0000598 | Explicit 2D evidence |
| noesy | nuclear Overhauser enhancement spectroscopy | CHMO | CHMO:0000607 | NOESY family mapping; raw pulse program retained |
| cpmg | Carr-Purcell-Meiboom-Gill pulse sequence | CHMO | CHMO:0000718 | CPMG family mapping; raw pulse program retained |

| | | | | |
|---|---|---|---|---|
| hsqc | heteronuclear single quantum coherence | CHMO | CHMO:0000604 | HSQC family mapping |
| hmbc | heteronuclear multiple bond coherence | CHMO | CHMO:0000601 | HMBC family mapping |
| tocsy | total correlation spectroscopy | CHMO | CHMO:0000605 | TOCSY/DIPSI family mapping |
| cosy | correlation spectroscopy | CHMO | CHMO:0000599 | COSY family mapping |
| j_spectroscopy | J-spectroscopy | CHMO | CHMO:0000606 | J-resolved/JRES mapping |
| hrmas | high-resolution magic angle spinning | CHMO | CHMO:0000618 | HRMAS/MAS mapping when explicit |
| cw_nmr | continuous-wave nuclear magnetic resonance spectroscopy | CHMO | CHMO:0000713 | Kept distinct from FT-NMR |
| mri | magnetic resonance imaging | MMO | MMO:0000019 | Magnetic-resonance imaging branch; not routine NMR spectroscopy |

**Table S10: Distribution of both shared and unique mapped ionization source terms across MW and ML**

| Mapped label | Ontology | IRI | MW | ML | Shared |
|---|---|---|---|---|---|
| atmospheric pressure chemical ionization | PSI-MS | MS_1000070 | yes | yes | yes |
| desorption electrospray ionization | PSI-MS | MS_1002011 | yes | yes | yes |
| electron ionization | PSI-MS | MS_1000389 | yes | yes | yes |
| electrospray ionization | PSI-MS | MS_1000073 | yes | yes | yes |
| matrix-assisted laser desorption ionization | PSI-MS | MS_1000075 | yes | yes | yes |
| atmospheric pressure ionization | PSI-MS | MS_1000240 | yes | | |

| Mapped label | Ontology | IRI | MW | ML | Shared |
|---|---|---|---|---|---|
| inductively coupled plasma | PSI-MS | MS_1000059 | yes | | |
| paper spray ionization | PSI-MS | MS_1003235 | yes | | |
| chemical ionization | PSI-MS | MS_1000071 | | yes | |
| field ionization | PSI-MS | MS_1000258 | | yes | |
| nanoelectrospray | PSI-MS | MS_1000398 | | yes | |
| secondary electrospray ionization | PSI-MS | MS_1003775 | | yes | |
| glow discharge ionization | PSI-MS | MS_1000259 | | yes | |
| ionspray mass spectrometry | CHMO | CHMO_0000486 | | yes | |
| heated electrospray ionization | unresolved | not accepted | | yes | |
| rapid evaporative ionization | unresolved | not accepted | | yes | |

Note: glow discharge ionization and ionspray mass spectrometry are proxy/caveated mappings for deposited ML terms, not exact source-label matches.

**Table S11: Raw value normalization dictionary for ion polarity terms**

| Repository | Raw value from direct field | Reporting class | Mapped label | PSI-MS accession | Handling |
|---|---|---|---|---|---|
| MW | POSITIVE | positive_only | positive polarity acquisition | MS:1000077 | accepted direct polarity |
| MW | NEGATIVE | negative_only | negative polarity acquisition | MS:1000076 | accepted direct polarity |
| MW | POSITIVE and NEGATIVE | both_positive_and_negative | mixed polarity acquisition | MS:1003774 | accepted direct dual-mode polarity |

| Repository | Raw value from direct field | Reporting class | Mapped label | PSI-MS accession | Handling |
|---|---|---|---|---|---|
| ML (Typos from depositors) | positive, Positive, positive scan, postive, postivie | positive_only | positive polarity acquisition | MS:1000077 | accepted direct polarity or normalized typo |
| ML (Typos from depositors) | negative, negative scan, negtive | negative_only | negative polarity acquisition | MS:1000076 | accepted direct polarity or normalized typo |
| ML | positive \| negative | negative, negative | positive, Positive and negative, positive or negative` | both_positive_and_negative | mixed polarity acquisition |
| ML | alternating, first positive and then negative, and equivalent direct switching strings | switching_or_alternating | alternating polarity acquisition | MS:1002833 | accepted direct switching evidence |

**Table S12: Unresolved MetaboLights scan-polarity values**

| Study | Assay file | Raw Parameter Value[Scan polarity] | Classification | Why unresolved |
|---|---|---|---|---|
| MTBLS10439 | a_MTBLS10439_LC-MS_positive_hilic_metabolite_profiling.txt | Full ms resolution: 60000 | Metadata-placement error | Resolution metadata, not polarity |
| MTBLS1051 | a_MTBLS1051_GC-MS_Batch1_GC-MS___metabolite_profiling.txt | none | Unavailable/non-informative | No assignable polarity evidence |
| MTBLS1051 | a_MTBLS1051_GC-MS_Batch2_GC-MS___metabolite_profiling-1.txt | none | Unavailable/non-informative | No assignable polarity evidence |
| MTBLS12537 | a_MTBLS12537_GC-MS_positive__metabolite_profiling.txt | SIM | Metadata-placement error | Scan/acquisition mode, not polarity |
| MTBLS12563 | a_MTBLS12563_GC-MS___metabolite_profiling.txt | SCAN/SIM | Metadata-placement error | Scan/acquisition mode, not polarity |
| MTBLS12692 | a_MTBLS12692_GC-MS___metabolite_profiling.txt | SCAN/SIM | Metadata-placement error | Scan/acquisition mode, not polarity |
| MTBLS12759 | a_MTBLS12759_LC-MS___metabolite_profiling.txt | not available | Explicitly unavailable | No assignable polarity evidence |
| MTBLS12759 | a_MTBLS12759_LC-MS___metabolite_profiling-1.txt | not available | Explicitly unavailable | No assignable polarity evidence |
| MTBLS12759 | a_MTBLS12759_LC-MS___metabolite_profiling-2.txt | not available | Explicitly unavailable | No assignable polarity evidence |
| MTBLS12936 | a_MTBLS12936_LC-MS_negative_reverse-ph | 70-1050 | Metadata-placement error | Mass range, not polarity |

| Study | Assay file | Raw Parameter Value[Scan polarity] | Classification | Why unresolved |
|---|---|---|---|---|
| | ase_metabolite_profiling.txt | | | |
| MTBLS12951 | a_MTBLS12951_LC-MS___metabolite_profiling-1.txt | Not Available | Explicitly unavailable | No assignable polarity evidence |
| MTBLS12951 | a_MTBLS12951_LC-MS___metabolite_profiling.txt | Not Available | Explicitly unavailable | No assignable polarity evidence |

**Table S13: Typos in ion polarity entries within ML**

| Study | Raw value | Normalized polarity |
|---|---|---|
| MTBLS10263 | negtive | negative |
| MTBLS10446 | postive | positive |
| MTBLS11965 | postive | positive |
| MTBLS13378 | postivie | positive |
| MTBLS13502 | postive | positive |
| MTBLS9522 | postive | positive |
| MTBLS9835 | negtive | negative |

**Table S14: Ontology terms for acquisition polarity node all mapped using PSI-MS**

| Reporting class | Final ontology label | PSI-MS IRI |
|---|---|---|
| positive_only | positive polarity acquisition | MS:1000077 |
| negative_only | negative polarity acquisition | MS:1000076 |
| both_positive_and_negative | mixed polarity acquisition | MS:1003774 |
| switching_or_alternating | alternating polarity acquisition | MS:1002833 |

**Table S15: Unique values → ontology mappings for the ion polarity node**

| **MW unique raw values** | | | | |
|---|---|---|---|---|
| **Raw value** | **Reporting class** | **Final mapped label** | **IRI** | **Records** |
| POSITIVE | positive_only | positive polarity acquisition | MS:1000077 | 3,428 |
| NEGATIVE | negative_only | negative polarity acquisition | MS:1000076 | 2,417 |
| POSITIVE and NEGATIVE | both_positive_and_negative | mixed polarity acquisition | MS:1003774 | 4 |
| UNSPECIFIED | unknown_or_missing | none | none | 547 |
| [blank] | not_applicable or unknown_or_missing | none | none | 296 |
| **ML unique raw values** | | | | |
| positive, Positive, positive scan, postive, postivie | positive_only | positive polarity acquisition | MS:1000077 | 2,671 |
| negative, negative scan, negtive | negative_only | negative polarity acquisition | MS:1000076 | 2,102 |
| positive \| negative, negative \| positive, Positive and negative, positive or negative | both_positive_and_negative | mixed polarity acquisition | MS:1003774 | 48 |
| alternating, alternating \| positive \| negative, negative \| alternating, positive \| alternating, positive \| negative \| alternating, first positive and then negative | switching_or_alternating | alternating polarity acquisition | MS:1002833 | 424 |

**Table S16: ML unresolved / non-mapped raw values**

| Raw values | Handling | Records |
|---|---|---|
| [blank] | missing or not applicable depending on assay type | 330 |
| not available, Not Available, none | unresolved non-polarity value | 7 |
| SCAN/SIM, SIM, Full ms resolution: 60000, 70-1050 | unresolved non-polarity value | 5 |

**Table S17: Source records used for separation-method extraction**

| Repository | Extraction unit | Main source | Primary evidence fields |
|---|---|---|---|
| MW | One /analysis row | REST-derived MW raw analysis records | chromatography_type, with same-row fallbacks from column, chromatography system, analysis summary, MS type, and instrument metadata |
| ML | One ISA assay object or assay file | REST/ISA-derived ML assay metadata and assay-file rows | Parameter Value[Column type], CE instrument, column model, chromatography instrument, assay type, technology platform, assay filename, and explicit study-design descriptors |

**Table S18: Normalization examples**

| Raw evidence pattern | Canonical term |
|---|---|
| RP, reverse phase, reversed phase, C18, C8, PFP, ODS, HSS T3 | reversed_phase_lc |

| | |
|---|---|
| HILIC, hydrophilic interaction, amide, ZIC-pHILIC | hilic |
| LC, liquid chromatography, LC-MS | liquid_chromatography |
| UPLC, UHPLC, ultra-performance liquid chromatography | uhplc |
| HPLC | hplc |
| GC, GC-MS, DB-5, HP-5, VF-5, RTX-5 | gas_chromatography |
| GCxGC, GC x GC, 2D-GC, Pegasus 4D | two_dimensional_gc |
| CE, CE-MS, capillary electrophoresis | capillary_electrophoresis |
| FIA, FIAMS, flow injection | flow_injection_analysis |
| DI, DIMS, direct infusion, shotgun lipidomics, NanoMate, TriVersa NanoMate | direct_infusion_no_separation |
| MALDI, DESI, MS imaging, SESI, paper spray, ICP-MS, PTR-MS, when no chromatographic or electrophoretic separation was present | no_chromatographic_separation |

**Table S19: Manual audit and freeze-stage rule fixes for separation method**

| Audit fix | Reason |
|---|---|
| Removed broad WAX ion-exchange matching | Prevented GC WAX-phase columns from being misclassified as ion-exchange chromatography |
| Ran chromatography checks before non-applicable checks | Prevented valid chromatography evidence from being suppressed by neighboring modality text |
| Treated Unspecified, Other, Multiple, and placeholders as non-informative | Allowed fallback only when another direct field supplied clear evidence |
| Supported both flat and numbered MW /analysis layouts | Ensured all MW analysis rows were parsed consistently |
| Added bare CE and CE-MS patterns | Recovered capillary electrophoresis records |

| | |
|---|---|
| Collected suffixed ML column fields | Captured fields such as Parameter Value[Column type 1] and related variants |
| Used all ML column-type values per assay | Preserved multi-column or multi-method assay evidence |
| Moved DI/FIA checks before broad instrument checks | Prevented direct infusion and flow injection from being swallowed by the broad LC/MS context |
| Prevented FIA-MS from being classified as DI-MS | Kept flow injection distinct from direct infusion |
| Added explicit ML study-design descriptor fallback | Rescued records with direct analytical descriptors but no assay-file separation field |
| Added paper spray and secondary electrospray no-separation contexts | Prevented explicit no-separation MS source rows from being treated as missing |
| Mapped direct infusion to PSI-MS MS:1000060 | Completed ontology mapping for direct-infusion sample introduction |

**Table S20: Ontology terms for separation method node all mapped using CHMO and PSI-MS**

| Canonical term | Final ontology label | Ontology | CURIE |
|---|---|---|---|
| reversed_phase_lc | reversed-phase liquid chromatography | CHMO | CHMO:0001050 |
| hilic | hydrophilic interaction chromatography | CHMO | CHMO:0002262 |
| liquid_chromatography | liquid chromatography | CHMO | CHMO:0001004 |
| uhplc | ultra high performance liquid chromatography | CHMO | CHMO:0000795 |
| hplc | high-performance liquid chromatography | CHMO | CHMO:0001009 |
| gas_chromatography | gas chromatography | CHMO | CHMO:0001002 |
| two_dimensional_gc | two-dimensional gas chromatography | CHMO | CHMO:0002886 |

| | | | |
|---|---|---|---|
| capillary_electrophoresis | capillary electrophoresis | CHMO | CHMO:0001024 |
| normal_phase_lc | normal-phase liquid chromatography | CHMO | CHMO:0001051 |
| ion_exchange_chromatography | ion-exchange chromatography | CHMO | CHMO:0001014 |
| ion_chromatography | ion chromatography | CHMO | CHMO:0002874 |
| ion_pair_chromatography | ion-pair chromatography | CHMO | CHMO:0002280 |
| supercritical_fluid_chromatography | supercritical-fluid chromatography | CHMO | CHMO:0001003 |
| flow_injection_analysis | flow-injection analysis | CHMO | CHMO:0002891 |
| direct_infusion_no_separation | infusion | PSI-MS | MS:1000060 |
| no_chromatographic_separation | no chromatographic separation | none | none |

**Table S21: Higher-level grouping of canonical separation-method terms for interpretation**

| **Higher-level group** | **Canonical terms** |
|---|---|
| LC mode-specific methods | reversed_phase_lc, hilic, normal_phase_lc, ion_exchange_chromatography, ion_pair_chromatography, ion_chromatography |
| Broad/platform LC descriptors | liquid_chromatography, hplc, uhplc |
| GC methods | gas_chromatography, two_dimensional_gc |
| Electrophoretic separation | capillary_electrophoresis |
| Specialized chromatography | supercritical_fluid_chromatography |
| Sample-introduction/no-separation MS contexts | flow_injection_analysis, direct_infusion_no_separation |
| Internal no-chromatography reporting class | no_chromatographic_separation |

**Table S22: Evidence hierarchy for mass-analyzer type**

| Repository | Evidence tier | Field | Use |
|---|---|---|---|
| MW | Primary | ms_instrument_type | Direct analyzer/model evidence |
| MW | Secondary | ms_instrument_name | Instrument-model fallback |
| MW | Secondary | analysis_summary | Explicit analyzer/model fallback only |
| ML | Primary | Parameter Value[Mass analyzer] | Direct assay-file analyzer field |
| ML | Secondary | Parameter Value[Instrument] | Instrument-model fallback |
| ML | Secondary | Technology platform | Used only for explicit analyzer/model evidence |
| ML | Secondary | Assay filename | Used only for explicit analyzer/model evidence |
| ML | Secondary | Assay type label | Used only for explicit analyzer evidence |

**Table S23: PSI-MS ontology mapping and higher-level grouping for mass-analyzer type**

| Analyzer family | Internal canonical term | PSI-MS label | Accession | PSI-MS branch |
|---|---|---|---|---|
| Orbital/FT high-resolution | fourier_transform_ion_cyclotron_resonance | fourier transform ion cyclotron resonance | MS:1000079 | mass analyzer type |
| Orbital/FT high-resolution | orbitrap | orbitrap | MS:1000484 | mass analyzer type |

| | | | | |
|---|---|---|---|---|
| TOF and hybrid TOF | quadrupole_time_of_flight | quadrupole time-of-flight instrument | MS:1003763 | instrument class |
| TOF and hybrid TOF | time_of_flight | time-of-flight | MS:1000084 | mass analyzer type |
| TOF and hybrid TOF | ion_trap_time_of_flight | ion trap time-of-flight instrument | MS:1003764 | instrument class |
| Quadrupole architectures | triple_quadrupole | triple quadrupole instrument | MS:1003762 | instrument class |
| Quadrupole architectures | single_quadrupole | quadrupole instrument | MS:1003950 | instrument class |
| Quadrupole architectures | quadrupole | quadrupole | MS:1000081 | mass analyzer type |
| Quadrupole architectures | hybrid_triple_quadrupole_linear_ion_trap | triple quadrupole linear ion trap instrument | MS:1003765 | instrument class |
| Ion-trap family | ion_trap | ion trap | MS:1000264 | mass analyzer type |
| Ion-trap family | linear_ion_trap | linear ion trap | MS:1000291 | mass analyzer type |
| Magnetic sector | magnetic_sector | magnetic sector | MS:1000080 | mass analyzer type |

**Table S24: Normalization and special-case rules**

| Evidence pattern | Final handling |
|---|---|
| Q Exactive, Exploris, Orbitrap Fusion, Lumos, ID-X, LTQ Orbitrap | orbitrap |
| QTOF, Q-TOF, TripleTOF, ZenoTOF, micrOTOF-Q, timsTOF, Synapt, Xevo G2, maXis | quadrupole_time_of_flight |
| GC-TOF, CE-TOF, MALDI-TOF, PTR-TOF, reflectron TOF | time_of_flight |
| QQQ, QqQ, TSQ, Xevo TQ-S, API 2000, Agilent 7000 series | triple_quadrupole |
| QTRAP, Q Trap, QTRAP 6500, QQQ and LIT | hybrid_triple_quadrupole_linear_ion_trap |
| LTQ-FT, FTMS, FT-ICR | fourier_transform_ion_cyclotron_resonance |
| Broad quadrupole only | quadrupole |
| Single-quadrupole model names | single_quadrupole |
| Stellar, Waters Cyclic MS | left unresolved pending policy |

**Table S25: ML curation rules for retaining or excluding disease terms**

| Evidence field | Role |
|---|---|
| Study title | Strong evidence when disease is part of the title |
| Study summary/description | Primary free-text evidence for disease, infection, clinical condition, model, or endpoint |
| Study design descriptors | Semi-structured disease/model/condition evidence |
| Descriptor accessions | Support for descriptor interpretation |
| Study Factor Name | Indicates disease/status/grouping variables |
| Study Factor Type | Supports interpretation of factor names |

| | |
|---|---|
| Factor Value[] column names | Identifies disease-related grouping columns |
| Factor Value[] values | Confirms disease/control, infected/uninfected, resistant/susceptible, or model groups |

**Table S26: Disease-node ontology policy**

| **Mapping layer** | **Description** | **Use in main disease analyses** |
|---|---|---|
| Strictly accepted mapped | Specific or defensible disease/condition mapped to MONDO, MeSH, DOID, NCIT, or EFO | Yes |
| Broad fallback mapped | Broadly supported handle, such as cancer, infection, or disease | Report separately; optional fallback |
| Secondary ontology mapped | Non-disease ontology context is retained only when faithful to the deposited MW value | Context only |
| Valid unmapped literal | Disease-like literal without a safe accepted IRI | Search/review only |
| Excluded context or fragment | Fragment, organism-only, method/background, non-disease term | No |
| Non-extractable | ML study with no supported disease/condition label | No |
| MW missing endpoint | MW endpoint returned no disease value | No |

**Table S27: Disease-node coverage and mapping layers**

| **Metric** | **MW** | **ML** |
|---|---|---|
| Total studies | 4,118 | 2,712 |
| Studies with disease evidence | 2,466 (59.88%) | 1,369 (50.48%) |
| Studies without extractable disease evidence | 1,652 (40.12%) | 1,343 (49.52%) |

| Metric | MW | ML |
|---|---|---|
| Studies with a strictly accepted disease IRI | 2,458 (59.69% of all; 99.68% of disease-evidence studies) | 1,184 (43.66% of all; 86.49% of disease-extractable studies) |
| Studies with a strict or broadly accepted IRI | 2,462 (59.79% of all; 99.84% of disease-evidence studies) | 1,306 (48.16% of all; 95.40% of disease-extractable studies) |
| Studies with disease literature but no accepted IRI | 4 (0.10% of all; 0.16% of disease-evidence studies) | 63 (2.32% of all; 4.60% of disease-extractable studies) |

**Table S28: Categorization of non-extractable disease terms for ML**

| Category | Meaning |
|---|---|
| method_resource_or_workflow_without_disease | Method development, software, spectral libraries, annotation workflows, QC/resource papers. |
| plant_agronomy_food_or_non_disease_trait | Plant/food/agronomy studies about cultivars, fruit, crops, leaves, fermentation, growth traits, not plant disease. |
| diet_nutrition_exposure_or_intervention_without_disease | Diet, supplementation, feeding, compound exposure, drug treatment, or intervention studies without a disease endpoint. |

| | |
|---|---|
| microbial_ecology_or_biofilm_without_host_disease | Microbiome, microbial community, bacterial adaptation, biofilm, resistance, in the context of no explicit host disease. |
| normal_biology_development_or_trait_without_disease | Development, genotype, mutant/wild-type comparisons, growth, reproduction, normal phenotype/trait studies. |
| environmental_or_abiotic_stress_without_disease | Drought, heat, salt, hypoxia, oxidative stress, temperature, osmotic stress, water deficit. |
| cell_line_or_in_vitro_perturbation_without_disease | Cell culture, overexpression, induction, mitochondrial/cellular perturbation without disease assignment. |
| healthy_reference_control_or_baseline_metabolome | Healthy/control/reference/baseline metabolome studies with no disease case group. |
| exercise_physiology_or_performance_without_disease | Exercise, training, rugby/athletic performance, and physical activity physiology without a disease endpoint. |
| insufficient_or_generic_metadata_for_disease | Metadata is too vague or fragmented to assign disease safely. |

**Table S29. Manual ML Disease-Node Curation Examples**

| Study | Initial issue | Final handling | Rationale |
|---|---|---|---|
| MTBLS12886 | Infectious diarrhea evidence was initially under-recovered despite an explicit diarrhea factor/value structure and ETEC context. | Reclassified as disease-extractable. Strictly mapped diarrhea and ETEC infection; retained infectious diarrhea and enterotoxigenic Escherichia coli infection as valid unmapped review literals. | Factor Value[Diarrhea] and the study text supported a real infectious diarrhea condition. |
| MTBLS1708 | Axolotl forelimb regeneration contained disease-like terms such as wound/healing/blastema. | Rejected as non-disease-extractable; retained regeneration, wound healing, and blastema as context terms. | The study concerns normal regeneration biology, not a disease or clinical condition. |
| MTBLS5868 | Cell overgrowth and osmotic stress were initially disease-like but arose from an in vitro mechanistic perturbation study. | Rejected as non-disease-extractable; retained cell overgrowth, osmotic stress response, and cell senescence as context terms. | Study design, factors, or sample grouping supported no disease assignment. |
| MTBLS6758 | Biofilm and organism terms leaked into disease candidates, including Biofilms, Mycobacterium, and Mycobacterium tuberculosis. | Retained supported disease concept tuberculosis; moved biofilm/organism-only terms to excluded/context terms. | Mentions of biofilms and organisms are not disease terms unless there is explicit host-disease evidence. |

| | | | |
|---|---|---|---|
| MTBLS8966 | Acinetobacter baumannii infection and antimicrobial resistance were biologically relevant but lacked a safe, accepted, and precise IRI. | Kept broad fallback bacterial infection; retained Acinetobacter baumannii infection and antimicrobial resistance as valid unmapped review literals. | Preserves useful disease/infection/AMR context without forcing unsafe ontology mappings. |

**Table S30: Classifier rules for flagging non-biological samples for DEM analysis**

| Expanded meaning | Raw terms are treated as this class |
|---|---|
| Field blank | fieldblank, field_blank |
| Generic blank | blk, blank |
| Process blank | pblk, processblank, process_blank |
| Method blank | mblk, methodblank, method_blank |
| Reagent blank | rblk, reagentblank, reagent_blank |
| Solvent blank | sblk, solventblank, solvent_blank |
| Quality control | qc, qcs |
| Pooled QC | pooledqc, pooled_qc, poolqc, pool_qc |
| Internal standard | istd, internalstd, internalstandard, internal_standard |
| Reference material | nist |
| System suitability | sst |
| Long-term reference | ltr |
| Study reference dilution series | srd |
| Biological recovery reference | biorec |

**Table S31: Representative examples showing why some deposited literals didn't give valid RefMet mappings**

| Category | Repository | Representative unmatched examples |
|---|---|---|
| Untargeted feature ID / m/z literal | MW & ML | Unknown14 (ST003206); ID14326 (ST004153); Unknown 8 (6.07ppm) (ST001706); Unknown2.16to2.17ppm (ST002417); Unknown 11 (9.36ppm) (ST001924) CPD0-1656(ST002066,ST003053) ID14326 (ST004153)<br>Neg_3-Phenyllactic acid_166.0628_2.249_ExRate0 (ST002517)<br>Neg_3-Phenyllactic acid_166.0628_2.249_ExRate2 (ST002517)<br>Neg_3-Phenyllactic acid_166.0628_2.249_ExRate3 (ST002517)<br>99.85655 (MTBLS13574) |
| Residual unresolved named compound | MW | "(+)-2,7-Dideoxypancratistatin" — ST002444<br>"(1,3-diphenylpropoxy)sulfonic acid" — ST003179, ST003565<br>"(10E,12Z)-Hexadeca-10,12-dienoyl-CoA" — ST003378"<br>(10E,15Z)-9,12,13-Trihydroxy-10,15-octadecadienoic acid" — ST004200<br>"(12E)-14-Pentyloxacyclotetradec-12-ene-2,11-dione" — ST004200 |
| Residual unresolved named compound | ML | "noise" — MTBLS2396<br>Δ9tetrahydrocannabinol — MTBLS315<br>α -allenylagmatine — MTBLS315<br>α-D-arabinofuranose — MTBLS315<br>α-D-arabinopyranose — MTBLS315 |
| Chain-composition shorthand / lipid-like | MW | "1,2-dilinoleoyl-GPC (18:2/18:2)" — ST002909, ST003520<br>"1,2-dilinoleoyl-GPE (18:2/18:2)*" — ST002909, ST003520<br>"1,2-dipalmitoyl-GPC (16:0/16:0)" — ST002909, ST003520<br>"1,2-dipalmitoyl-GPE (16:0/16:0)*" — ST002909, ST003520<br>"CL(1'-[18:2(9Z,12Z)/0:0],3'-[18:2(9Z,12Z)/0:0])" — ST003378 |
| Chain-composition shorthand / lipid-like | ML | (20S,24E)-20,26-Dihydroxy-24-dammaren-3-one; ... ;MG(i-24:0/0:0/0:0) — MTBLS12654<br>(2E)-Piperamide-C5:1 — MTBLS10376, MTBLS12931, MTBLS1693, MTBLS2145, MTBLS4437<br>(2E,4E,8E)-Piperamide-C9:3 — MTBLS11941, MTBLS13169, MTBLS13197, MTBLS13914<br>(2E,4E,8E)PiperamideC9:3 — MTBLS12809<br>(2E,6E)-Piperamide-C7:2 — MTBLS11752, MTBLS11941, MTBLS4437 |

| | | |
|---|---|---|
| Long systematic / IUPAC name | MW | "(+)-3,4-Dihydro-3,8-dihydroxy-3-methylbenz(a)anthracene-1,7,12(2H)-trione" — ST004432<br>"(-)-Epigallocatechin 3-gallate 7-glucoside 4""""-glucuronide" — ST003768<br>"(10E,12E,23E,47E)-14,18,19,22,31,33,40,41-Octahydroxy-... hydrogen sulfate" — ST004200<br>"(13R,17R)-17-[(1R,2S)-1,2-dihydroxyheptyl]-1,5,10-triazabicyclo[11.4.0]heptadec-15-en-11-one" — ST004432<br>"(13Z)-N-[(2S,3R,4E)-1,3-Dihydroxy-4-hexadecen-2-yl]-13-docosenamide" — ST004200 |
| Long systematic / IUPAC name | ML | α-(2,6-anhydro-3-deoxy-D-arabino-heptulopyranosid)onate 7-phosphate — MTBLS1569, MTBLS630, MTBLS751<br>α-D-xylosyl-(1->6)-β-D-glucosyl-(1->4)-β-D-glucose — MTBLS630<br>'(6R)-2-hydroxy-2-methyl-6-... heptyl hydrogen sulfate' — MTBLS12340<br>'2-((4R)-4-... pentanamido)ethane-1-sulfonic acid' — MTBLS12340<br>((1R,2R,5R)-2-(2,6-Dimethoxy-4-(2-methyloctan-2-yl)phenyl)-... methanol — MTBLS12310, MTBLS9100 |
| Peptide sequence | MW | Ala Ala Ala Arg Phe — ST002405<br>Ala Arg Asp Pro Val — ST002405<br>Ala Arg Glu Arg — ST002405<br>Ala Glu Asn Arg — ST004088<br>Ala Glu Lys Asp — ST004088 |
| Peptide sequence | ML | Ala Ala Ala Arg Phe — MTBLS10453, MTBLS13575<br>Ala Ala Ala Asp — MTBLS13958<br>Ala Ala Ala Gln — MTBLS4437<br>Ala Ala Ala Glu — MTBLS13958, MTBLS4437<br>Ala Ala Ala His — MTBLS13958, MTBLS4437 |
| Numeric-only literal | MW | No examples in this repository for this category. |
| Numeric-only literal | ML | "1" — MTBLS2396"10" — MTBLS2396"100" — MTBLS2396"101" — MTBLS2396"102" — MTBLS2396 |
| Catalog / code-like literal | MW | A0A0C4DH29 — ST003666<br>A4UGR9 — ST003666<br>A6H8Y1 — ST003666<br>A6NDG6 — ST003666 |

| | | |
|---|---|---|
| | | AA861 — ST003622 |
| Catalog / code-like literal | ML | A145015 — MTBLS338<br>A159003 — MTBLS338<br>A171005 — MTBLS338<br>A174001 — MTBLS338<br>A178003 — MTBLS338 |
| Slash-mixture / multi-candidate literal | MW | "2,3-Pyridinedicarboxylic Acid/Quinolinic Acid" — ST002797<br>"3D-3,5/4-trihydroxycyclohexane-1,2-dione" — ST004432<br>"5-O-Methyl-2,3,5/4,6-pentahydroxycyclohexanone" — ST003160, ST003179, ST003565<br>"F-1,6/2,6-DP" — ST002876<br>"PC(DiMe(11,3)/DiMe(9,5))" — ST003378 |
| Slash-mixture / multi-candidate literal | ML | (+)-Piperitone/d-Piperitone — MTBLS5943<br>(+/)2Hydroxy4(methylthio)butanoic acid — MTBLS12809<br>(+/)Ibipinabant — MTBLS12809<br>(+/)Pelletierine — MTBLS12809<br>(+/-) 5-iPF2alpha-VI-(d11) — MTBLS11844, MTBLS1490 |
| Pipe-delimited multi-value string | MW | 1,5,6,7-TETRAHYDRO-4H-INDOL-4-ONE\|2-PHENYLACETAMIDE\|N-BENZYLFORMAMIDE — ST001671, ST001688<br>17\|A-Carboxy-17\|A-formyloxy Dexamethasone — ST004389<br>6\|A-Hydroxy-7\|A-(thiomethyl)spirolactone — ST004459<br>ALPHA-GALACTOSE 1-PHOSPHATE\|ALPHA-GLUCOSE 1-PHOSPHATE\|GLUCOSE-6-PHOSPHATE\|MANNOSE 6-PHOSPHATE — ST001671, ST001688<br>sp\|O75223\|GGCT_HUMAN Gamma-glutamylcyclotransferase — ST001072 |
| Pipe-delimited multi-value string | ML | (+)-7-(2-hydroxy-3-methyl-butyl)-6-methoxy-9-methyl-9H-[1,3]dioxolo[4,5-h]quinolin-8-one\|(+)-lunidine\|Lunidin\|Lunidine — MTBLS12719<br>(+)-Pinoresinol 4-O-[beta-D-Glucopyranosyl-(1\|right\|2)-... ] — MTBLS11800<br>(+)-clavaminol C\|(2R,3S)-2-acetylaminododecan-3-ol\|clavaminol C — MTBLS12764<br>(+-)-1,1'-Binaphthyl-carbonsaeure-8\|... — MTBLS12764<br>(-)-dihydropertusaric acid\|(-)589-Pertusarinic acid\|... — MTBLS12719 |

| | | |
|---|---|---|
| Character-encoding artifact | MW | "(1'S,2'S)-3',11'-dihydroxy-... 0Â?,â·]dodecan]-5'-en-4'-one" — ST002775<br>"(14-{3,4,5,11,16,17,18-heptahydroxy-...})oxidanesulfonic acid" — ST003378<br>"(3Î_5Î±,22Î_25S)-Spirosolan-3-yl 4-O-Î_-D-glucopyranosyl-Î_-D-galactopyranoside" — ST004200<br>"1,2,3,4-Tetrahydro-Î²-carboline-3-carboxylic acid" — ST002775<br>"2',4',11-trioxaspiro[...]-5,7,10-triol" — ST002776 |
| Character-encoding artifact | ML | (+)-Cycloolivil 4'-O-¦Â-D-glucopyranoside — MTBLS10453<br>(+)-Lyoniresinol 2¦Á-O-¦Â-D-glucopyranoside — MTBLS10453<br>(+)-pinoresinol-¦Â-D-glucoside; (+)-pinor — MTBLS11479<br>(+)-pinoresinol-Î²-D-glucoside; (+)-pinor — MTBLS10002<br>(+)-¦Á-Tocopherol — MTBLS10453 |
| Short acronym / token literal | MW | 0DH — ST004416<br>0DP — ST004416<br>0DPH — ST004416<br>1-AG — ST000257, ST000549, ST000593, ST000597, ST000659<br>1MVR — ST002576 |
| Short acronym / token literal | ML | -- — MTBLS12521, MTBLS12922, MTBLS13477, MTBLS4820, MTBLS5132<br>-ESA — MTBLS2384<br>1-AG — MTBLS10002, MTBLS11203, MTBLS12826, MTBLS13109, MTBLS13367<br>1-U — MTBLS10782<br>10cb — MTBLS12777 |
| Explicit unknown placeholder | MW | Unknown — ST001284, ST001285, ST001626, ST001946, ST002113<br>Unknown (carbon number 13) — ST004032<br>Unknown (carbon number 15) — ST004032<br>Unknown (carbon number 26) — ST004032<br>Unknown (carbon number 5) — ST004032 |
| Explicit unknown placeholder | ML | UNKNOWN_1 — MTBLS5163<br>Unknown — MTBLS1015, MTBLS10255, MTBLS10450, MTBLS10856, MTBLS12573<br>Unknown 045(100) 59(89.0) 74(67.8) — MTBLS627 |

| | | |
|---|---|---|
| RT-suffix literal | MW | (+)-Echinoisoflavanone_RT2 — ST004088<br>(-)-Matairesinol 4'-[apiosyl-(1->2)-glucoside]_RT1 — ST004088<br>(-)-Neolinderatin_RT2 — ST004088<br>(10E,15Z)-9,12,13-Trihydroxyoctadeca-10,15-dienoic acid_RT2 — ST004088<br>(20S)-3beta-Hydroxychola-5,16-dien-24-oic Acid_RT1 — ST004088 |
| RT-suffix literal | ML | No examples in this repository for this category. |
| Spreadsheet-formula error | MW | No examples in this repository for this category. |
| Spreadsheet-formula error | ML | #N/A — MTBLS12905<br>#NAME? — MTBLS11382, MTBLS12252, MTBLS12337, MTBLS12958, MTBLS12959 |

**Table S32: Extraction sources confirmed through evidence**

| Node | Repo | Primary evidence field(s) | Fallback evidence field(s), in order | Total |
|---|---|---|---|---|
| Species | MW | species endpoint raw value (1) | none | 1 |
| Species | ML | Characteristics[Organism] (1) | metabolights_organisms_api recovery call (1) | 2 |
| Sample source | MW | REST source endpoint, Sample source (1) | REST factors endpoint sample_source (1) → mwTab COLLECTION.SAMPLE_TYPE key (1) | 3 |
| Sample source | ML | Characteristics[Organism part] (1) | none | 1 |
| Ionization source | MW | ms_type (1) | ms_instrument_name (1) | 2 |
| Ionization source | ML | Parameter Value[Ion source] (1) | none | 1 |
| Ion polarity | MW | ion_mode (1) | none | 1 |
| Ion polarity | ML | Parameter Value[Scan polarity] (1) | none | 1 |

| **Node** | **Repo** | **Primary evidence field(s)** | **Fallback evidence field(s), in order** | **Total** |
|---|---|---|---|---|
| Separation method | MW | chromatography_type (1) | column_name, chromatography_system, analysis_summary, ms_type, ms_instrument_type (5) | 6 |
| Separation method | ML | Parameter Value[Column type], Parameter Value[Column model] (2) | Parameter Value[Chromatography Instrument], Study Assay Technology Platform, technologyPlatform/filename, Study Assay File Name, Study Design Descriptors (5) | 7 |
| Mass analyzer type | MW | ms_instrument_type (1) | ms_instrument_name (1) | 2 |
| Mass analyzer type | ML | Parameter Value[Mass analyzer] (1) | Parameter Value[Instrument] (1) | 2 |
| Analytical technique | MW | chromatography_type, ms_type, analysis_type, analysis_summary (4) | ms_instrument_name, ms_instrument_type (2) | 6 |
| Analytical technique | ML | Assay Type Label (1) | Study Assay Technology Platform, Study Assay File Name, Study Design Descriptors (Keywords) (3) | 4 |
| Disease | MW | REST disease endpoint (1) | none | 1 |
| Disease | ML | no structured field — manual curation pipeline using the following fields:<br>1. name<br>2. description<br>3. study_design<br>4. curator_keywords<br>5. study_id / accession-level context | — | 0 automated fields |

### Supplementary Figures

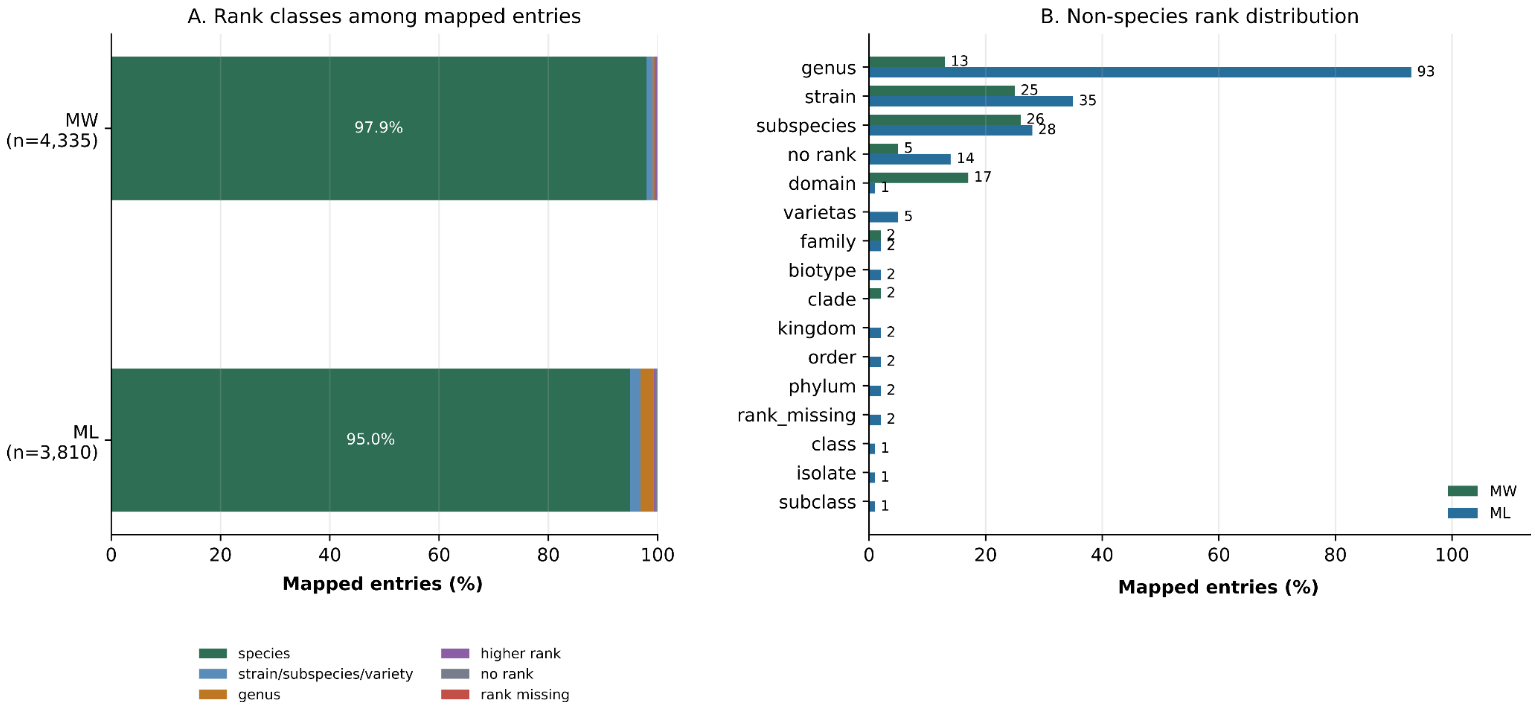


**Figure S1:** ***NCBITaxon rank distribution for species mappings across both MW and ML. A) Overall distribution across different ranks. B) Individual rank representation (other than species) across all mapped entries***

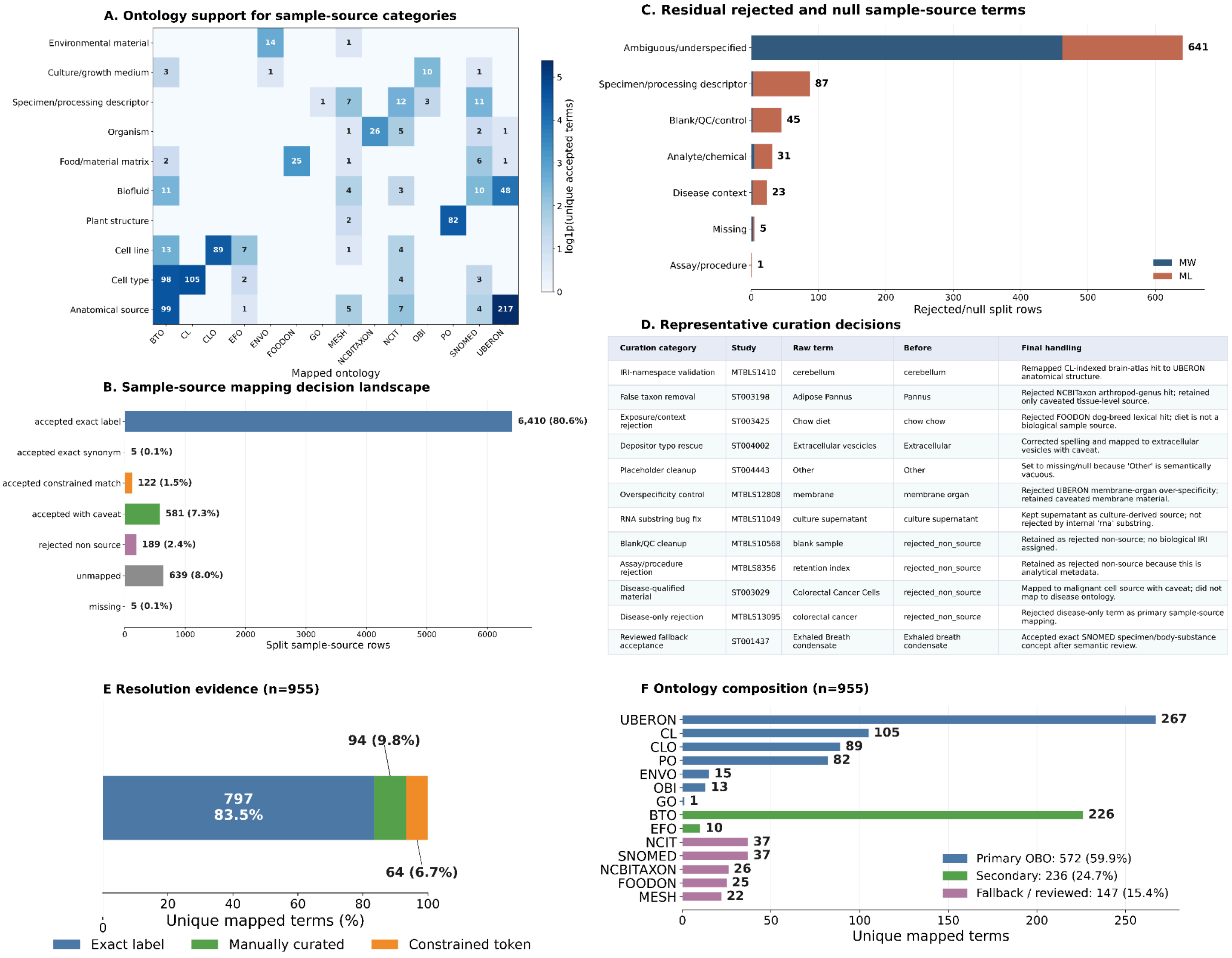


| Curation category | Study | Raw term | Before | Final handling |
|---|---|---|---|---|
| IRI-namespace validation | MTBLS1410 | cerebellum | cerebellum | Remapped CL-indexed brain-atlas hit to UBERON anatomical structure. |
| False taxon removal | ST003198 | Adipose Pannus | Pannus | Rejected NCBITaxon arthropod-genus hit; retained only caveated tissue-level source. |
| Exposure/context rejection | ST003425 | Chow diet | chow chow | Rejected FOODON dog-breed lexical hit; diet is not a biological sample source. |
| Depositor typo rescue | ST004002 | Extracellular vescicles | Extracellular | Corrected spelling and mapped to extracellular vesicles with caveat. |
| Placeholder cleanup | ST004443 | Other | Other | Set to missing/null because 'Other' is semantically vacuous. |
| Overspecificity control | MTBLS12808 | membrane | membrane organ | Rejected UBERON membrane-organ over-specificity; retained caveated membrane material. |
| RNA substring bug fix | MTBLS11049 | culture supernatant | culture supernatant | Kept supernatant as culture-derived source; not rejected by internal 'rna' substring. |
| Blank/QC cleanup | MTBLS10568 | blank sample | rejected_non_source | Retained as rejected non-source; no biological IRI assigned. |
| Assay/procedure rejection | MTBLS8356 | retention index | rejected_non_source | Retained as rejected non-source because this is analytical metadata. |
| Disease-qualified material | ST003029 | Colorectal Cancer Cells | rejected_non_source | Mapped to malignant cell source with caveat; did not map to disease ontology. |
| Disease-only rejection | MTBLS13095 | colorectal cancer | rejected_non_source | Rejected disease-only term as primary sample-source mapping. |
| Reviewed fallback acceptance | ST001437 | Exhaled Breath condensate | Exhaled breath condensate | Accepted exact SNOMED specimen/body-substance concept after semantic review. |

**Figure S2:** *(A) Ontology support for accepted sample-source categories, showing how different ontologies contributed to anatomical, biofluid, cell, plant, organism, environmental, food/material, and specimen descriptors. (B) Distribution of final mapping decisions across split sample-source entries. (C) Residual rejected or null entries by category and database. (D) Representative curation decisions illustrating false-positive removal, typo rescue, disease/context handling, blank/QC cleanup, and reviewed fallback mappings. (E) Resolution evidence for unique mapped terms, grouped by matching strategy. (F) Ontology composition of unique mapped terms, grouped by ontology tier (primary OBO, secondary, and fallback/reviewed).*

(i)

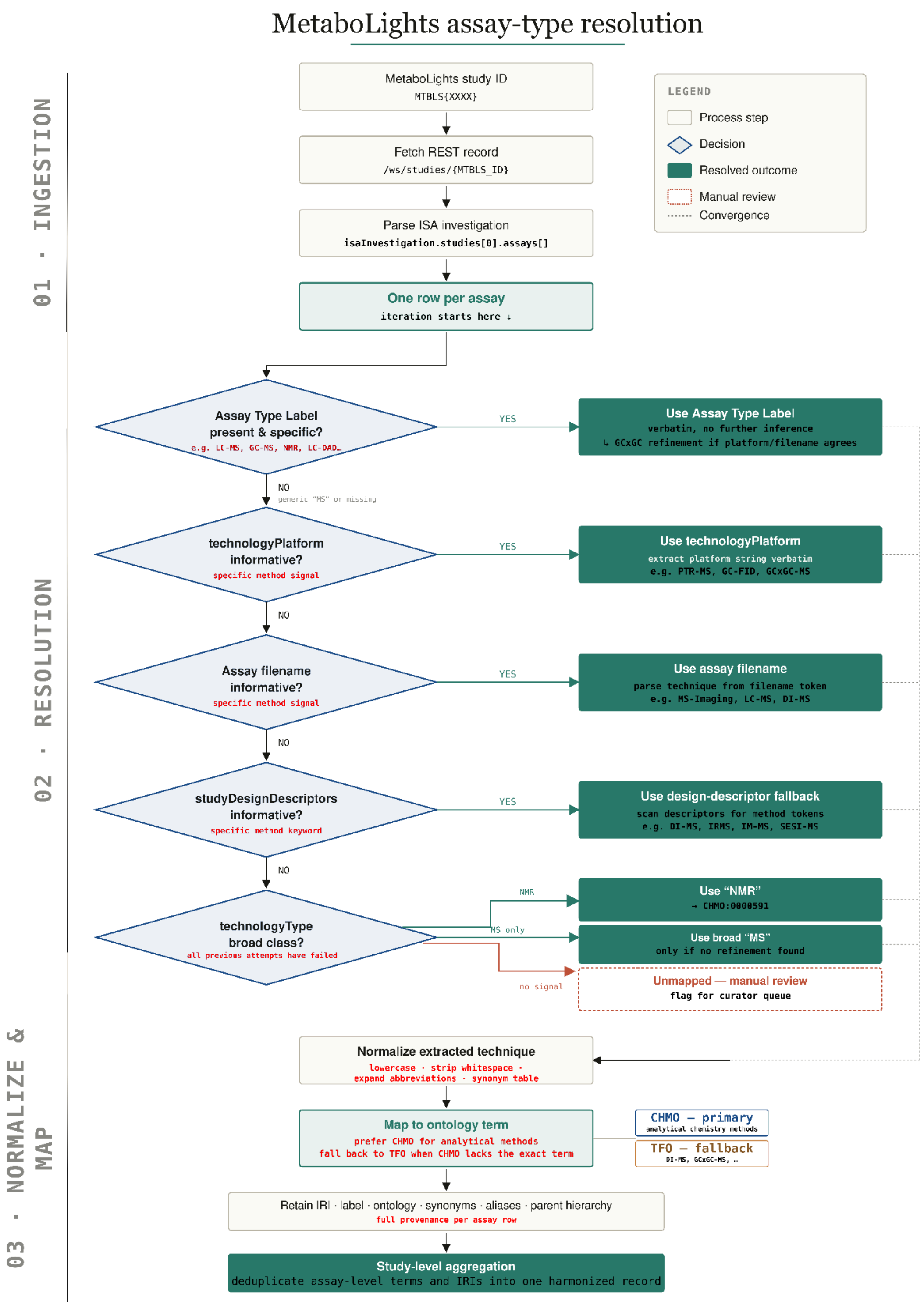
MetaboLights assay-type resolution
01 · INGESTION
02 · RESOLUTION
03 · NORMALIZE & MAP
MetaboLights study ID
MTBLS{XXXX}
Fetch REST record
/ws/studies/{MTBLS_ID}
Parse ISA investigation
isaInvestigation.studies[0].assays[]
One row per assay
iteration starts here ↓
LEGEND
Process step
Decision
Resolved outcome
Manual review
Convergence
Assay Type Label
present & specific?
e.g. LC-MS, GC-MS, NMR, LC-DAD…
YES
Use Assay Type Label
verbatim, no further inference
↳ GCxGC refinement if platform/filename agrees
NO
generic "MS" or missing
technologyPlatform
informative?
specific method signal
YES
Use technologyPlatform
extract platform string verbatim
e.g. PTR-MS, GC-FID, GCxGC-MS
NO
Assay filename
informative?
specific method signal
YES
Use assay filename
parse technique from filename token
e.g. MS-Imaging, LC-MS, DI-MS
NO
studyDesignDescriptors
informative?
specific method keyword
YES
Use design-descriptor fallback
scan descriptors for method tokens
e.g. DI-MS, IRMS, IM-MS, SESI-MS
NO
technologyType
broad class?
all previous attempts have failed
NMR
Use "NMR"
→ CHMO:0000591
MS only
Use broad "MS"
only if no refinement found
no signal
Unmapped — manual review
flag for curator queue
Normalize extracted technique
lowercase · strip whitespace ·
expand abbreviations · synonym table
Map to ontology term
prefer CHMO for analytical methods
fall back to TFO when CHMO lacks the exact term
CHMO – primary
analytical chemistry methods
TFO – fallback
DI-MS, GCxGC-MS, …
Retain IRI · label · ontology · synonyms · aliases · parent hierarchy
full provenance per assay row
Study-level aggregation
deduplicate assay-level terms and IRIs into one harmonized record

(ii)

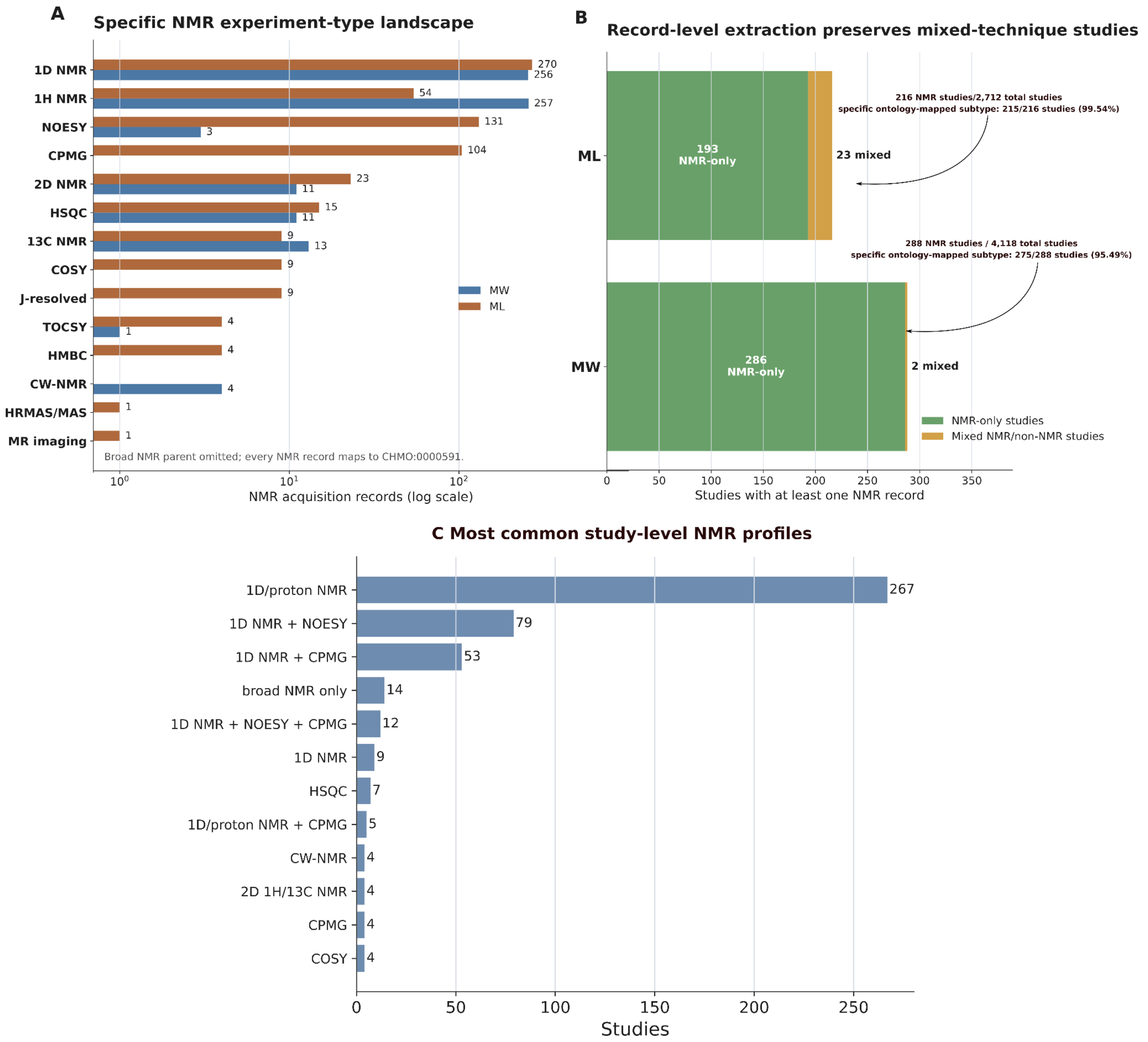


**Figure S3:** *(i) Workflow for resolving analytical technique terms for all assays belonging to ML; (ii) NMR experiment-type harmonization at the acquisition-record level. (A) Distribution of ontology-mapped NMR experiment-type terms across MW analysis records and ML assay records, excluding the broad parent NMR spectroscopy term. (B) Study-level NMR context after record-level extraction, separating NMR-only studies from studies containing both NMR and non-NMR records. (C) Most frequent study-level NMR profiles after aggregating mapped record-level NMR attributes.*

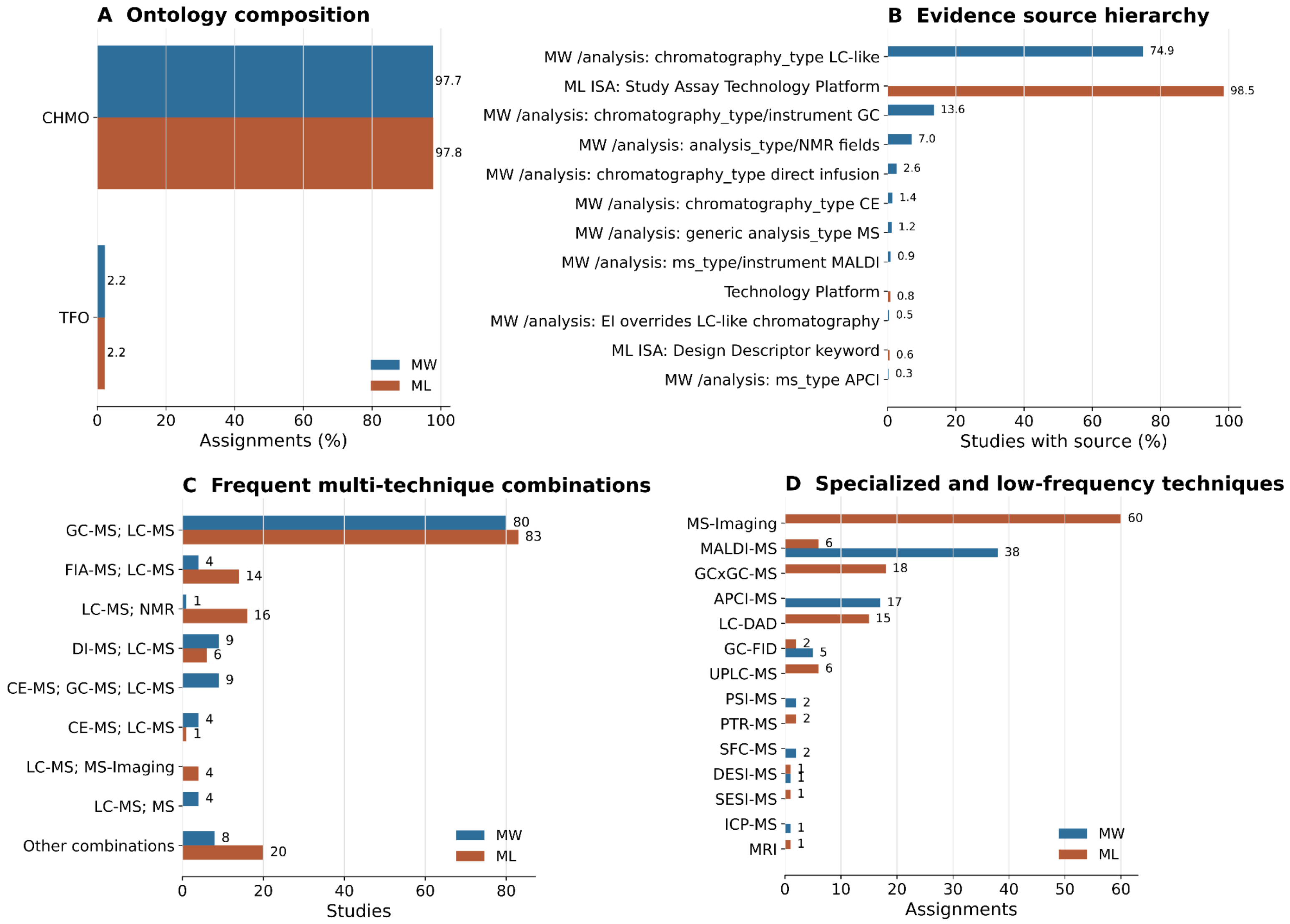


**Figure S4: Analytical-technique ontology and evidence summary**

*(A) Ontology composition of final MW and ML analytical-technique assignments, showing that nearly all terms mapped to CHMO, with TFO used for selected direct-injection/GCxGC terms. (B) Evidence-source hierarchy used for extraction, comparing MW `/analysis` endpoint fields with ML ISA-Tab assay descriptors. (C) Most frequent multi-technique study combinations across repositories. (D) Specialized and low-frequency analytical techniques were retained after harmonization.*

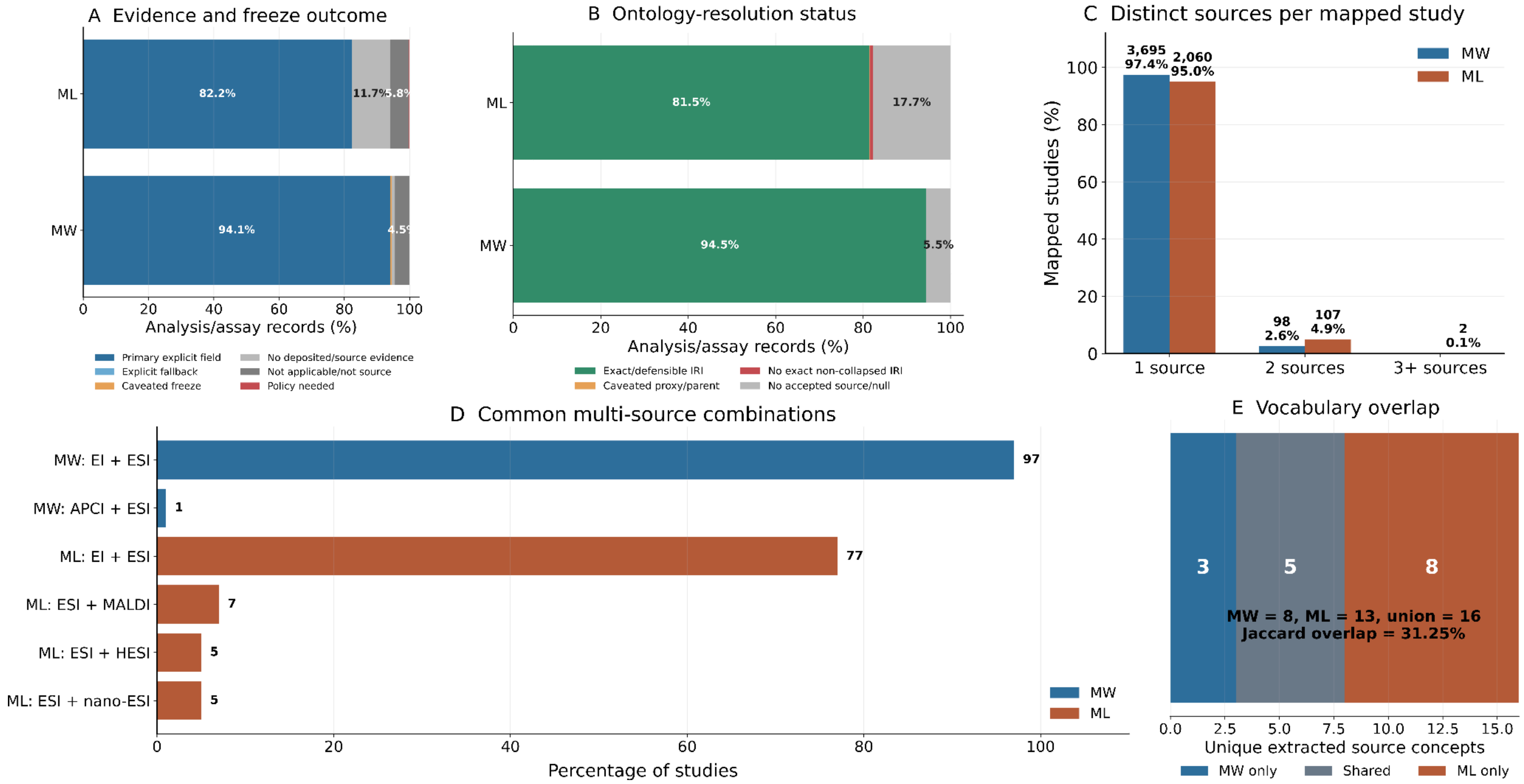


**Figure S5:** ***(A) Evidence and freeze outcome for ionization-source extraction across MW analysis records and ML assay records. (B) Ontology-resolution status showing exact/defensible IRI mappings, caveated proxy mappings, unresolved non-collapsed terms, and null/unaccepted records. (C) Number of distinct ionization-source terms per mapped study, showing how often studies contain multiple source types. (D) Most common multi-source combinations observed across studies, separated by repository. (E) Vocabulary overlap between MW and ML after harmonization, showing repository-specific and shared ionization-source concepts.***

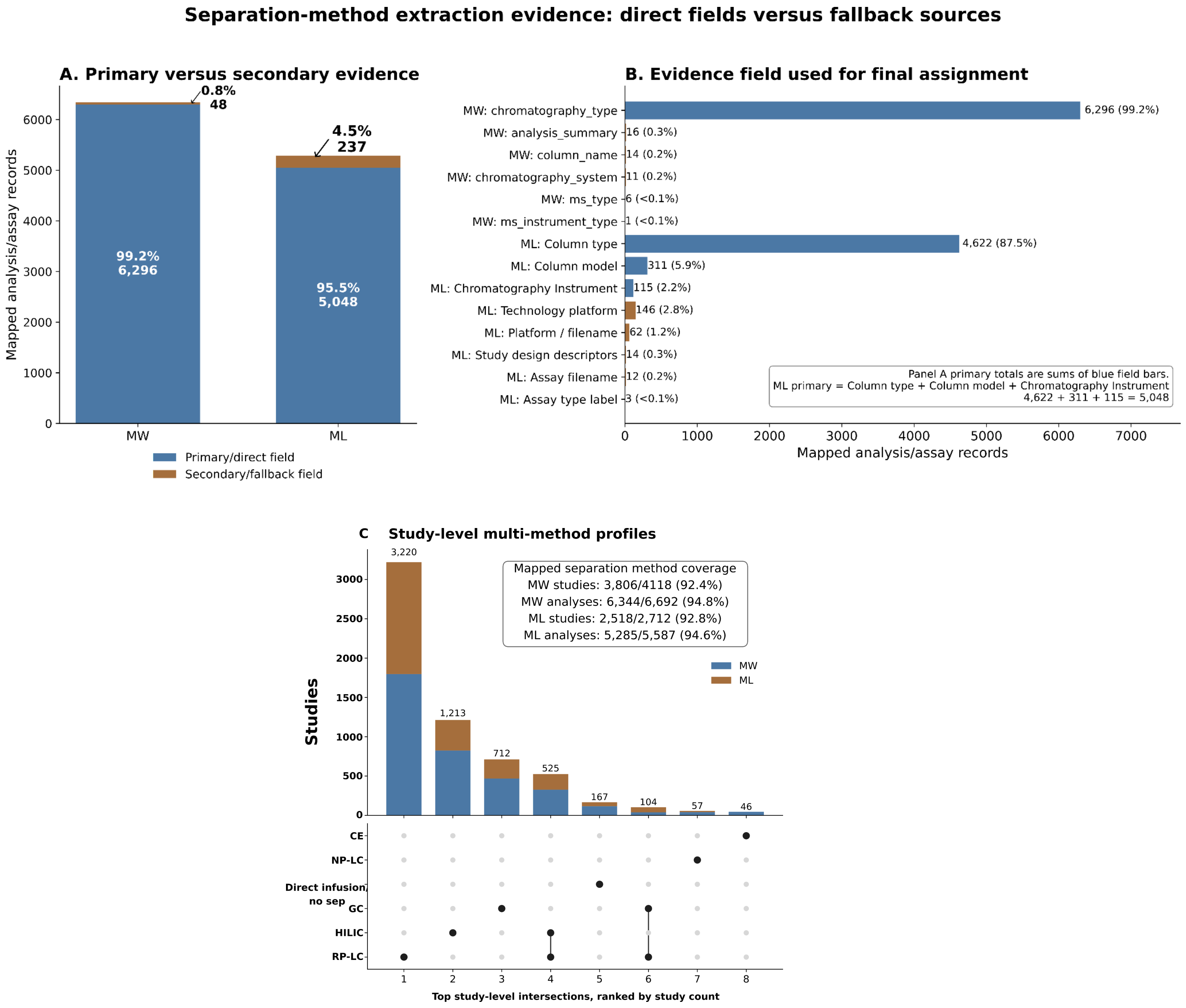


**Figure S6:** ***Evidence sources used for separation-method assignment. Panel A compares mappings derived from primary fields with mappings recovered through secondary or fallback evidence. Panel B shows the specific evidence fields contributing to final assignments. Secondary evidence recovered valid separation-method records that would otherwise have remained unresolved, while manual audit rules constrained fallback use to prevent over-interpretation. C) Study-level multi-method separation profiles. UpSet-style intersections show the most frequent single- and multi-method study profiles after aggregating record-level assignments; inset reports study- and record-level mapping coverage.***

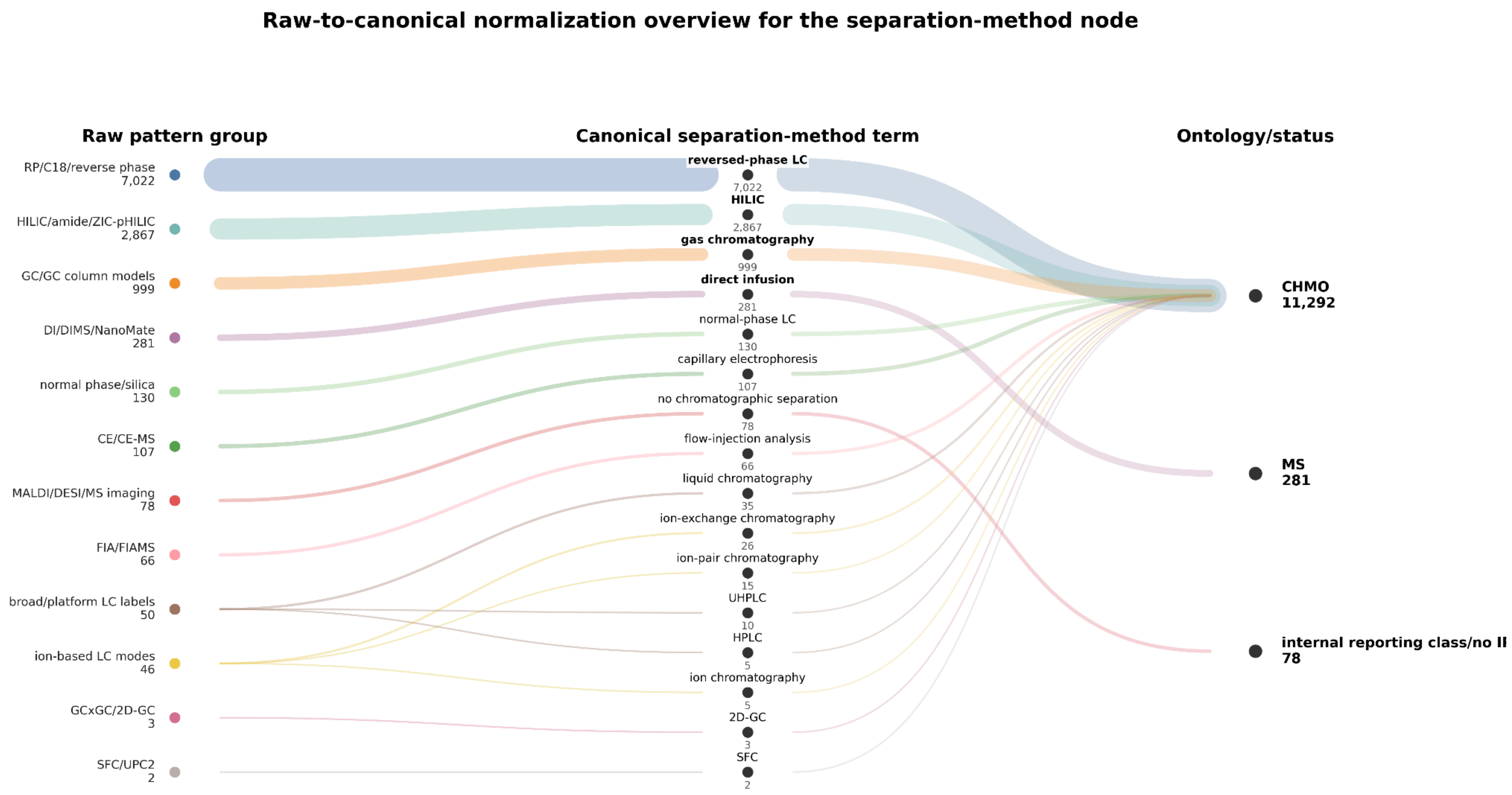


**Figure S7:** *Raw-to-canonical normalization overview for the separation-method node. Alluvial links show how grouped raw evidence patterns were normalized to canonical separation-method terms and then assigned to ontology/status categories. Link width is proportional to record-level assignments. Most terms mapped to CHMO, direct infusion mapped to PSI-MS, and explicit no-chromatographic-separation contexts were retained as an internal reporting class without an IRI. Raw-pattern groups are visualization-level audit summaries and do not replace the preserved row-level raw values.*

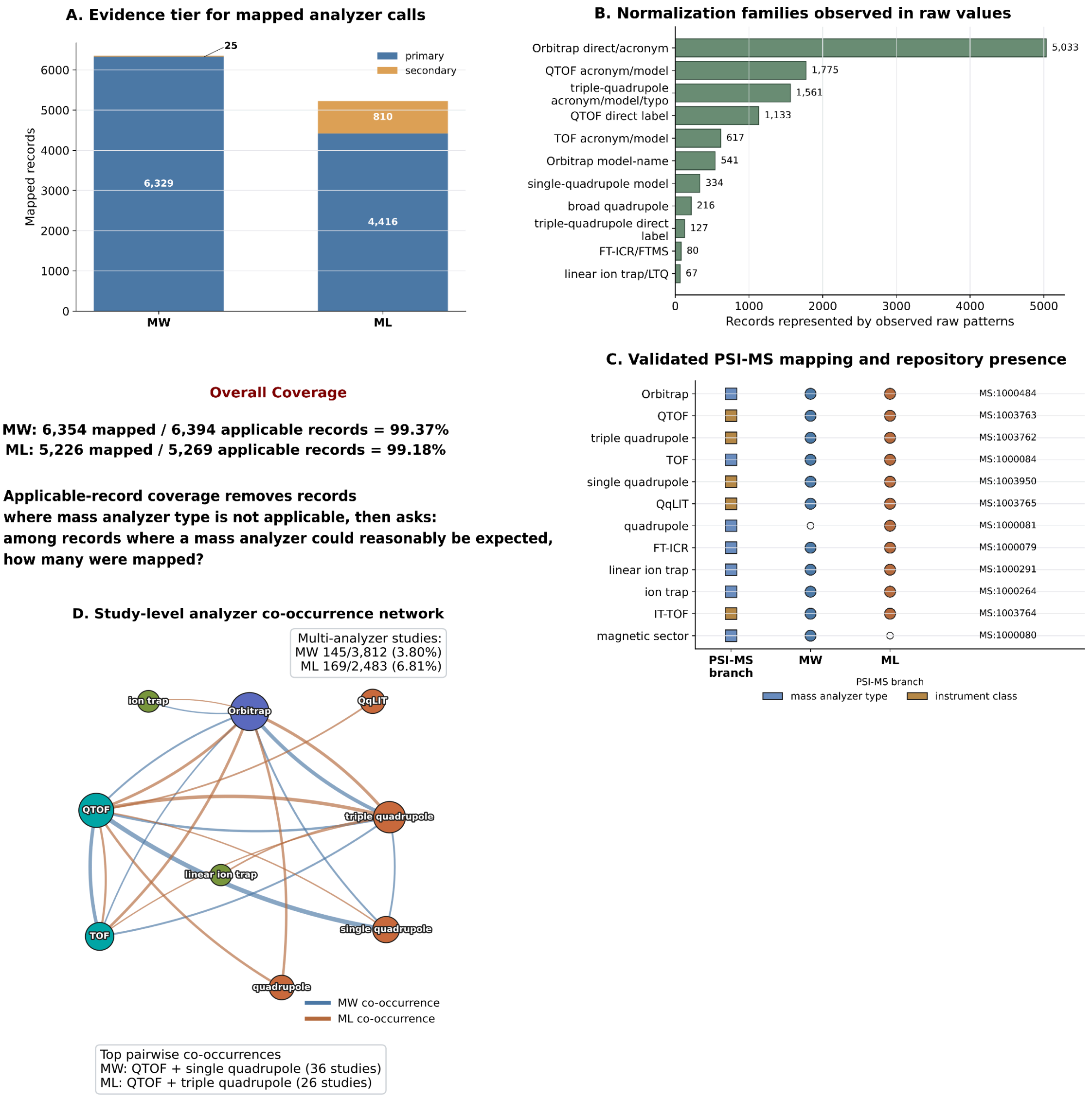


**Figure S8:** ***Supplementary audit of mass-analyzer type mapping.***

A) Contribution of primary and secondary evidence fields to final analyzer assignments. B) Normalization-family summary showing the main raw evidence groups collapsed into canonical mass-analyzer terms. C) PSI-MS term presence and repository overlap summary. Formal PSI-MS labels, accessions, branches, and analyzer-family groupings are reported in Table S23. D) Study-level analyzer co-occurrence network: Network of mass-analyzer terms that co-occur within the same study after aggregating assay-level assignments. Nodes represent analyzer terms; node size indicates how many studies (MW + ML combined) use that analyzer type at the study level. Size is scaled with a square-root transform, so very common types (Orbitrap, QTOF) do not visually dominate the whole panel; node colors match the analyzer-family groups in panel C; edge color indicates the repository; and edge thickness reflects the number of studies sharing that analyzer pair.

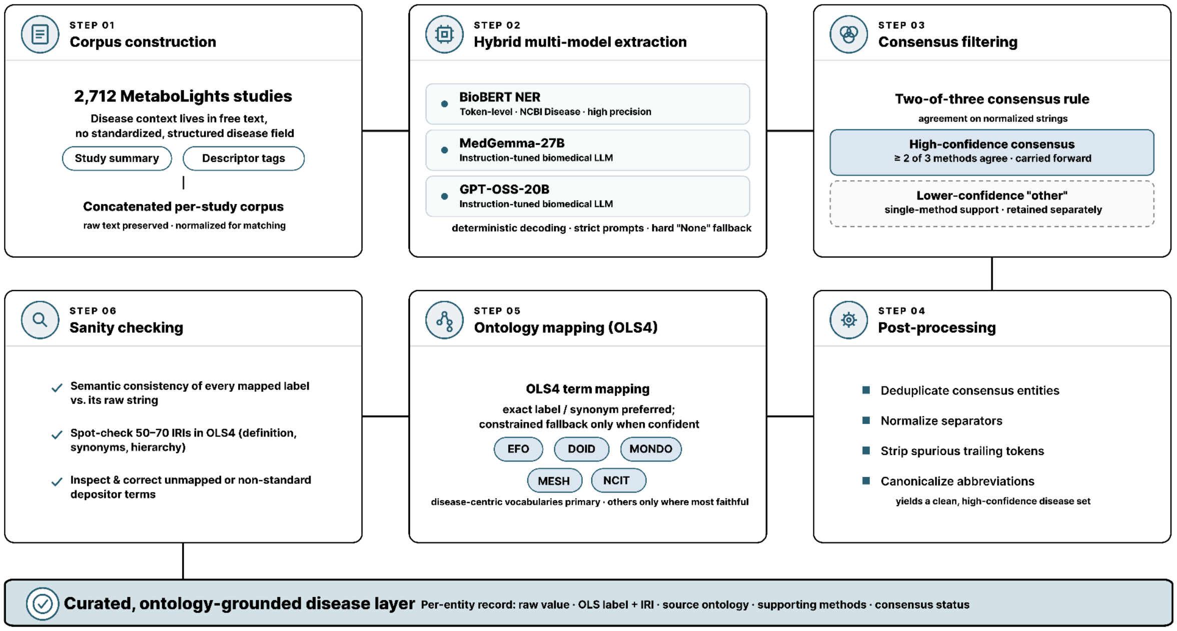


**Figure S9:** ***Methodology for disease node extraction from MetaboLights.***

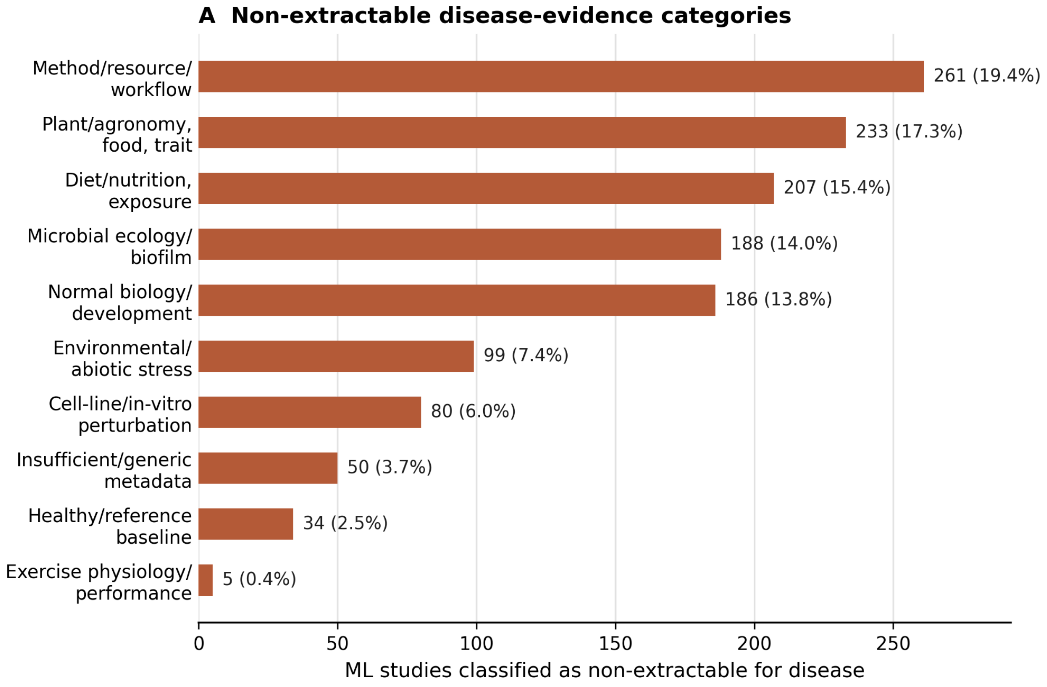


**Figure S10:**

**Breakdown of non-extractable disease terms from ML**

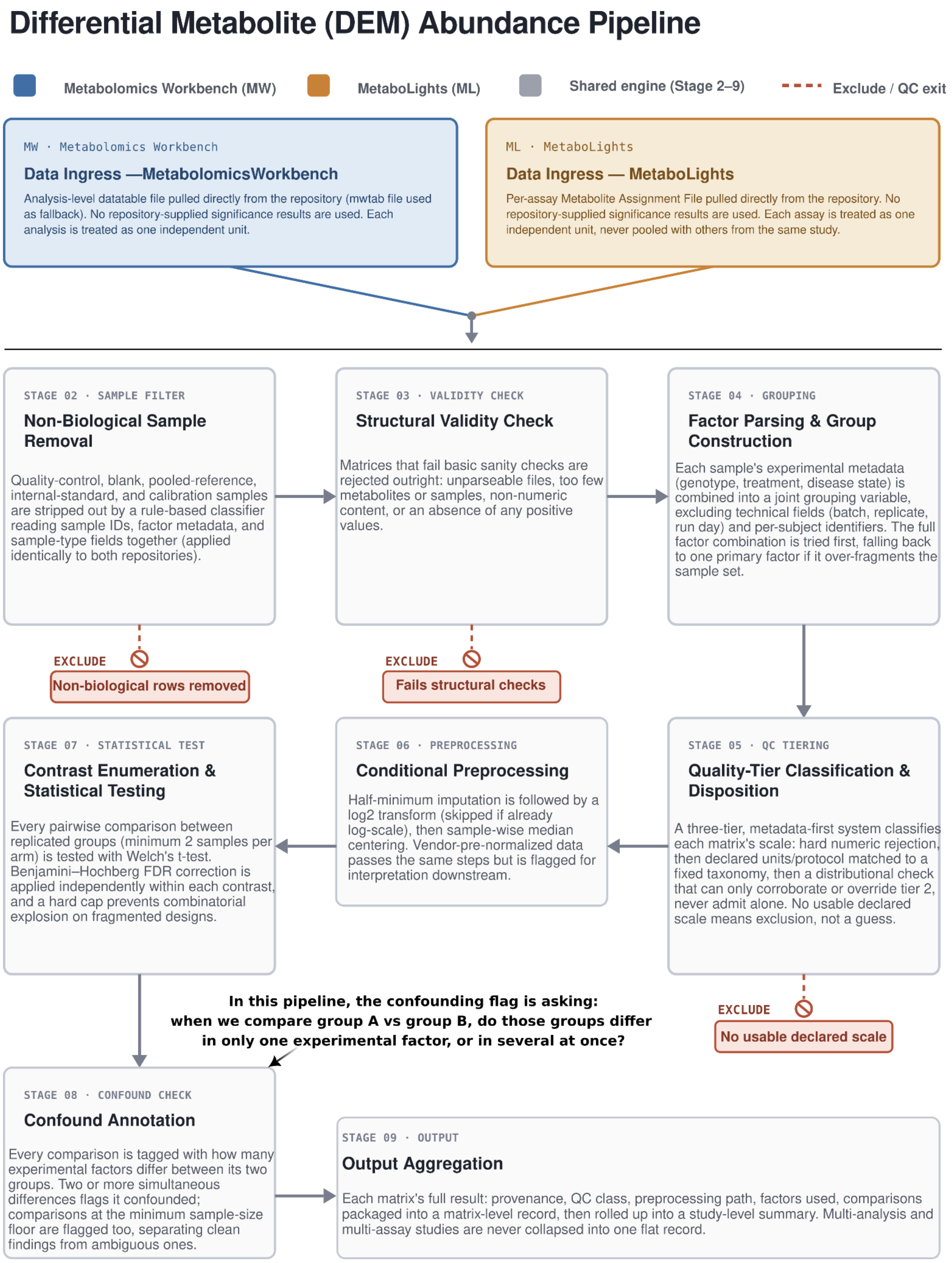


**Figure S11: Overall workflow explaining each step of the differential metabolite (DEM) abundance pipeline.**

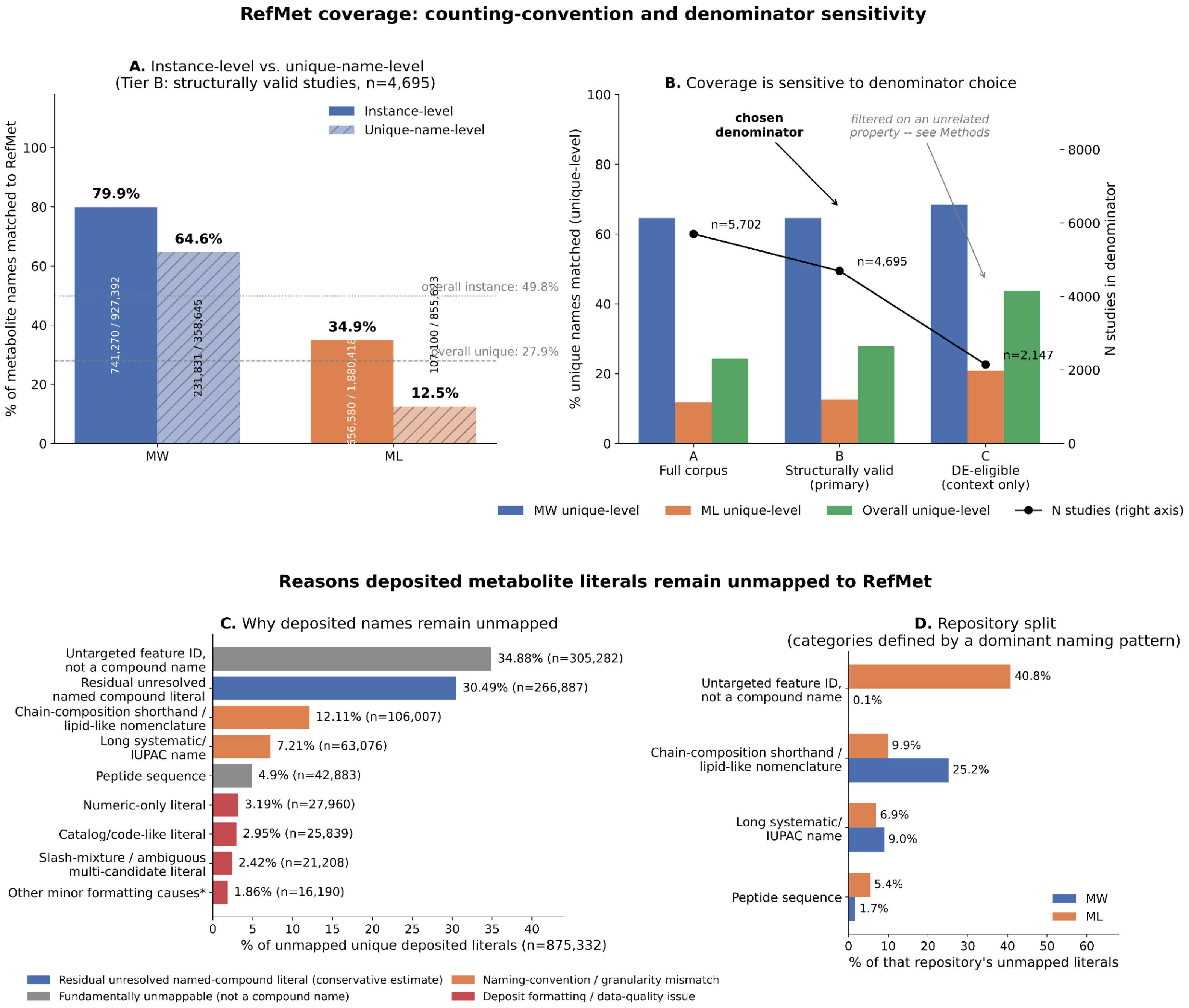


**Figure S12:** *RefMet mapping coverage: counting conventions, denominator sensitivity, and causes of non-mapping. (A) MW/ML coverage under two counting conventions (Tier B denominator, n=4,695 studies): instance-level = every metabolite-row occurrence counted, so repeated common metabolites count multiple times; unique-name-level = each distinct deposited name counted once, regardless of how often it recurs. (B) Unique-name-level coverage across denominator tiers A (full corpus), B (structurally valid, primary), C (DE-eligible); line shows N studies per tier. (C) Breakdown of unmapped unique literals (n=875,332) by cause, colored by mechanism. (D) Repository split for the four categories with an unambiguous naming-pattern definition.*

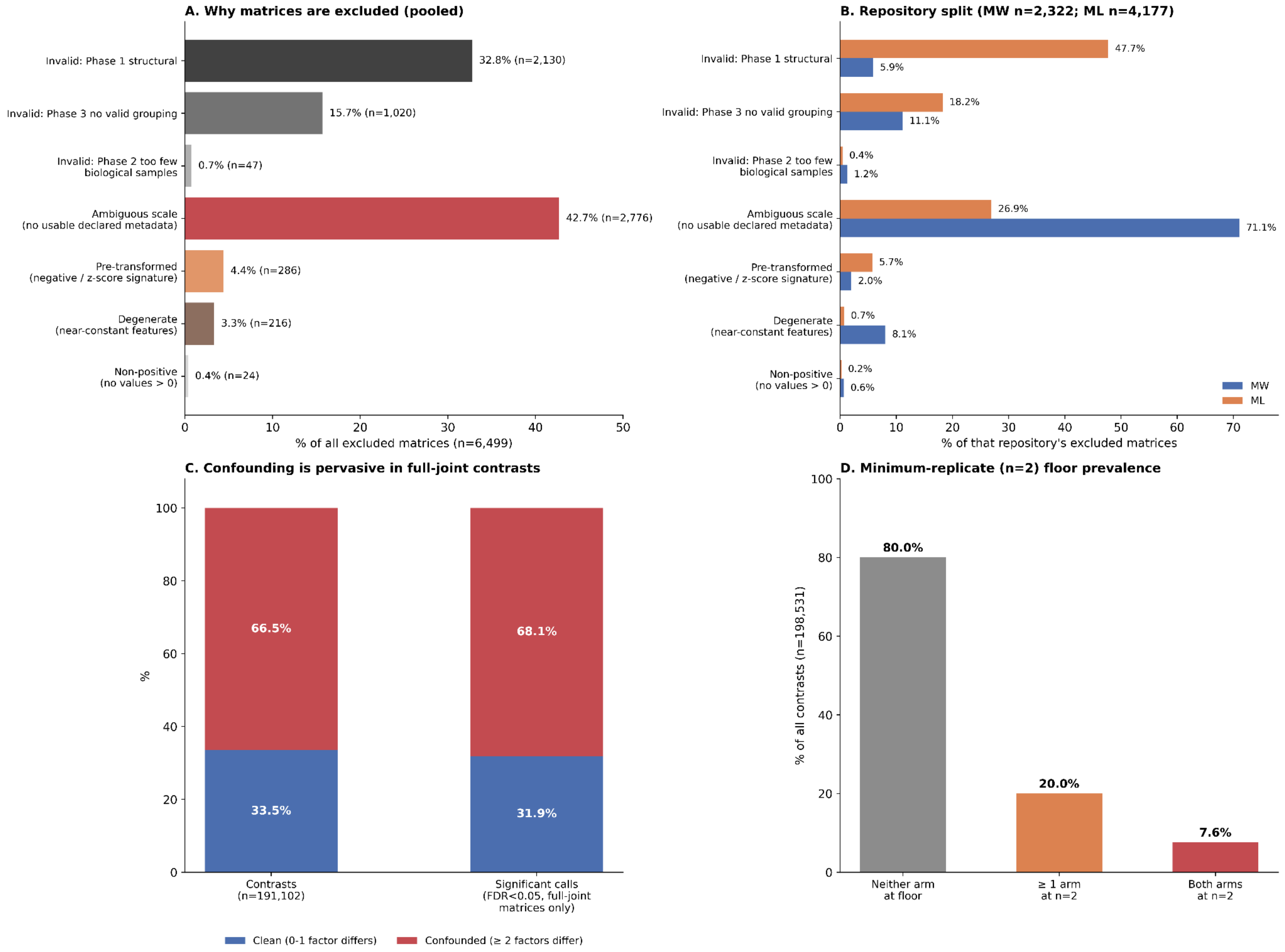


**Figure S13:** *Exclusion causes and result-reliability diagnostics. (A) Reasons deposited matrices were excluded from differential expression eligibility, pooled across repositories (n = 6,499 excluded matrices); color indicates the originating pipeline phase. (B) Same categories split by repository, as % of each repository's own excluded matrices (MW n = 2,322; ML n = 4,177). (C) Among full-joint pairwise contrasts with ≥2 experimental factors (n = 191,102), the fraction differing on a single factor ("clean") versus ≥2 factors simultaneously ("confounded"), compared against the fraction of FDR-significant calls contributed by each group. (D) Across all tested pairwise comparisons (n = 198,531), this panel shows how often the analysis involved the minimum allowed group size of 2 samples: in neither group, in one group, or in both groups.*

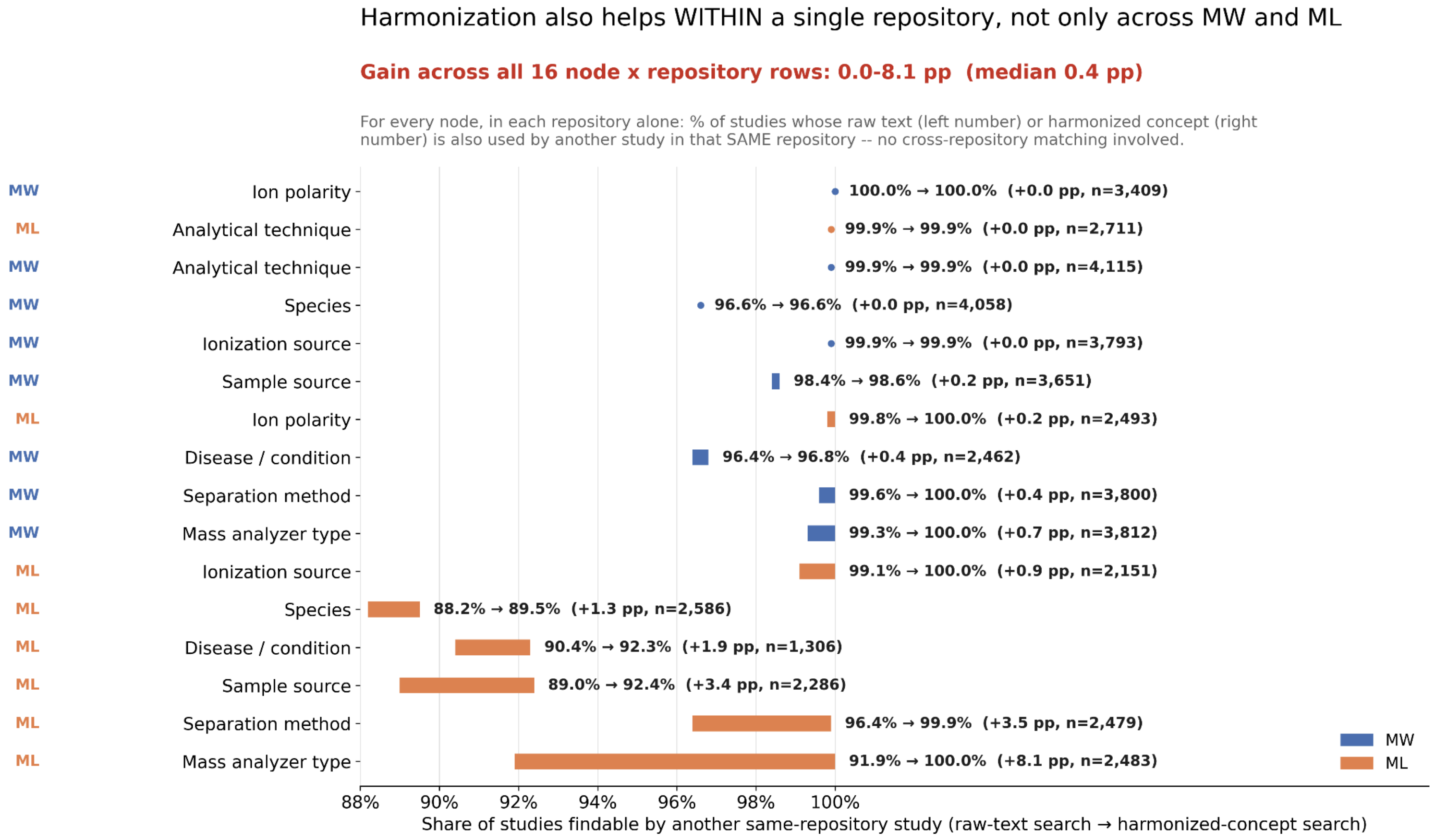


**Figure S14:** ***Ontology mapping also improves discovery within a single repository, independent of cross-repository matching.*** *For each of the 8 metadata nodes, computed separately within MW and within ML (16 rows total), the percentage of studies in that repository that share their raw text (left value) or their harmonized concept (right value) with at least one other study in the same repository, without MW-ML comparison is involved. Bars show the gain from raw-text to concept-based findability; dots mark rows with negligible gain. n = number of mapped studies for that node/repository. Rows are sorted by gain, largest first. pp = percentage points demonstrating the differences between two percentages.*

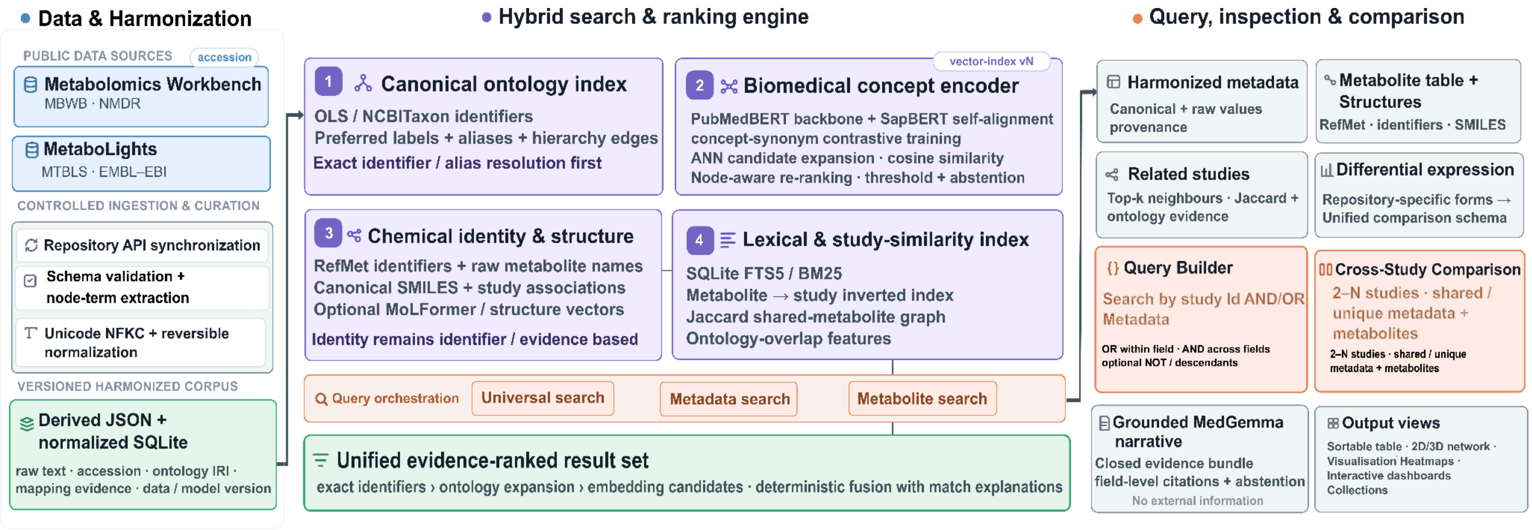


**Figure S15:** ***Complete platform architecture depicting the harmonized data backend and semantic retrieval frontend***
